# The impact of rare germline variants on human somatic mutation processes

**DOI:** 10.1101/2021.11.14.468508

**Authors:** Mischan Vali Pour, Ben Lehner, Fran Supek

## Abstract

Somatic mutations are an inevitable component of ageing and the most important cause of cancer. The rates and types of somatic mutation vary across individuals, but relatively few inherited influences on mutation processes are known. We performed a comprehensive gene-based rare variant association study with diverse mutational processes, using human cancer genomes from over 11,000 individuals of European ancestry. By combining burden and variance tests, we identify 207 associations involving 15 somatic mutational phenotypes and 42 genes that replicated in an independent data set at a FDR of 1%. We associated rare inherited deleterious variants in novel genes such as *MSH3*, *EXO1*, *SETD2*, and *MTOR* with two different forms of DNA mismatch repair deficiency, and variants in genes such as *EXO1*, *PAXIP1*, and *WRN* with deficiency in homologous recombination repair. In addition, we identified associations with other mutational processes, such as *APEX1* with APOBEC-signature mutagenesis. Many of the novel genes interact with each other and with known mutator genes within cellular sub-networks. Considered collectively, damaging variants in the newly-identified genes are prevalent in the population. We suggest that rare germline variation in diverse genes commonly impacts mutational processes in somatic cells.

## Introduction

Cancer is primarily a disease of mutations, alterations in the DNA sequence which result from replication errors and/or exogenous or endogenous DNA damaging agents^1^. Genomic instability via an increased rate of mutagenesis is a major enabling mechanism of cancer^2^, because it decreases the time needed to accrue the typically 2–10 somatic mutations in driver genes that are needed to initiate tumorigenesis^3, 4^. Thus, identifying genetic determinants of the variability of somatic mutation rates is important for understanding and predicting variation in cancer risk among individuals as well as for determining the mechanisms responsible for tumorigenesis. Moreover, many of the most effective cancer therapies target vulnerabilities associated with defects in specific repair pathways or mutation processes and many widely used therapeutics are themselves highly mutagenic^5–7^.

During the last decade, large-scale sequencing efforts have greatly enabled the analysis of somatic mutations in tumor genomes, both via whole-exome^8^ and whole-genome sequencing, either from primary^9^ or metastatic tumors^10^. These studies have identified driver genes and mutations^4, 11–13^ and also highlighted the abundance of ‘passenger’ mutations. Passenger mutations do not confer a selective advantage to the cancer cell and can be used to infer the sources of mutations in that particular individual and their tumor^1, 14^, either exogenous (chemicals, radiation) or endogenous (e.g. DNA replication errors, spontaneous deamination of cytosine)^15^.

Diverse mutation types have been analyzed in cancer genomes^14, 16^ including single base substitutions (SBS)^17, 18^ and the trinucleotide they are embedded in^17, 18^, double base substitutions (DBS)^19^, small deletions and indels (indels)^19^, copy number alterations (CNAs)^20^ and other structural variants (SVs)^21^. The extracted mutational patterns (often referred to as “mutational signatures”) capture biological, technical, and, in many cases, unknown sources of variation^14, 21^. In addition to the number and type of mutations, the regional distribution of mutations can also be informative about the activity of mutational processes^16^. For instance, in tumor genomes in which DNA mismatch repair (MMR) is impaired, there is reduced enrichment of mutations in late replicating regions (where presumably this pathway is normally less active or accurate)^16, 22^. Besides replication timing, the distribution of mutations also associate with locations of chromatin marks (e.g. H3K36me3^23, 24^ and H3K9me3^25^, the direction of DNA replication (leading vs lagging strand)^26, 27^, the direction of transcription (transcribed vs. untranscribed strand)^26^, chromatin accessibility (e.g. DNase I hypersensitive sites)^28^, CTCF/cohesin binding sites^29, 30^, and the inactive X-chromosome^31^. Moreover, the mitochondrial genome carries mutational patterns which differ from those in the nuclear genome^32, 33^.

While the catalogues of variation in somatic mutational patterns and rates between individuals are substantial^18, 19, 24, 26, 34, 35^, the extent to which this is determined by inherited genetic variants is less well understood. Examples of inherited variants that influence mutation processes include variants that cause familial cancer syndromes^36^. These include rare damaging germline variants (RDGVs) in the MMR genes *MSH2*, *MSH6*, *PMS2,* and *MLH1* that predispose to early-onset cancer of the colorectum and other organs (Lynch syndrome)^37^. Variants causing Lynch syndrome have been associated with several somatic mutational patterns^38^, most prominently short indels at microsatellite regions^19^, but also a relative enrichment of mutations in early replicating regions^22^, a replicative strand asymmetry^39^, and an increased number of mutations in several SNV-based cancer signatures^18^ due to the inefficient repair of base-base mismatches and smaller DNA loops. In addition, individuals with damaging variants in the genes *BRCA1*, *BRCA2, PALB2*, and *RAD51C* have an increased risk of breast, ovarian, pancreatic, and prostate cancer, and have distinct somatic mutational patterns^40, 41^ such as SBS Signature 3 mutations^17^, deletions at microhomology-flanked sites^17^, a copy number signature^20^ and several rearrangement based signatures^21, 42^. The products of these genes function in the repair of DNA double-strand breaks (DSBs) via homologous recombination, and impairment of this pathway necessitates repair via other, more error-prone mechanisms such as microhomology-mediated end joining, which create certain mutational patterns^43^. Finally, tumor genomes from individuals born with pathogenic variants in *TP53* frequently have complex chromosomal rearrangements (so-called chromothripsis)^44^.

These known examples illustrate how rare inherited variants can affect somatic mutation rates in humans^38^, and have motivated recent analyses aiming to identify additional variants associated with specific somatic mutational patterns. In a whole-genome pan-cancer association study^9^, a previously reported association^45, 46^ of a common deletion polymorphism in the coding region of *APOBEC3B*, altering APOBEC-signature mutagenesis, was replicated, and another nearby QTL locus associating with APOBEC burden was seen^9^. Known associations of rare variants in *BRCA1* and *BRCA2* with somatic CNA phenotypes were recapitulated^9^. In addition, an association between RDGVs in the DNA glycosylase *MBD4* with an increase of C>T mutations at CpG sites was reported^9^, which was also found in several independent studies^47, 48^. Furthermore, in a breast-cancer-specific study, the association of RDGVs with APOBEC and deficient homologous recombination (dHR) SNV signatures was investigated across ancestries, without detecting hits significant in both ancestries^49^.

These examples illustrate how genome-wide analyses can in principle be used to discover new germline determinants of human somatic mutation processes. In model organisms, genetic screens have revealed that mutations in many different genes influence mutation processes^50, 51^.

Here, we perform a comprehensive rare variant association study using sequencing data from three large-scale projects and identifiy novel genes associating with diverse somatic mutational processes. We use a gene-based testing approach combining a burden test and a variance test, two dimensionality reduction methods to define mutational phenotypes, considered multiple models of inheritance and multiple *in silico* variant prediction tools. We report 207 replicating associations involving 15 somatic mutational phenotypes and 42 genes, and an additional 149 associations involving 24 phenotypes and 44 genes at a more permissive false discovery rate. Rare inherited variants in a diverse set of genes therefore contribute to inter-individual differences in somatic mutation accumulation.

## Results

### Somatic mutation phenotypes in 15,000 human tumors

To capture inter-individual variation in somatic mutation processes, we extracted 56 mutational features from ∼15,000 tumor genomes analyzed as part of the Cancer Genome Atlas Program (TCGA)^8^, the Pan-Cancer Analysis of Whole Genomes (PCAWG)^9^ and the Hartwig Medical Foundation (Hartwig) study^10^. These features included different types of mutational signatures based on SBS, DBS, indels, and CNAs. Additionally we considered the distribution of SBS density across the genome with respect to transcription, gene expression, DNA replication (both the location and timing), chromatin state (accessibility via DNAse hypersensitivity, presence of active chromatin mark H3K36me3), CTCF binding sites, as well as localization on the X chromosome or in the mitochondrial genome; all of these were previously associated with local mutation rate variability (see Methods). (Fig. 1a).

**Fig. 1.**
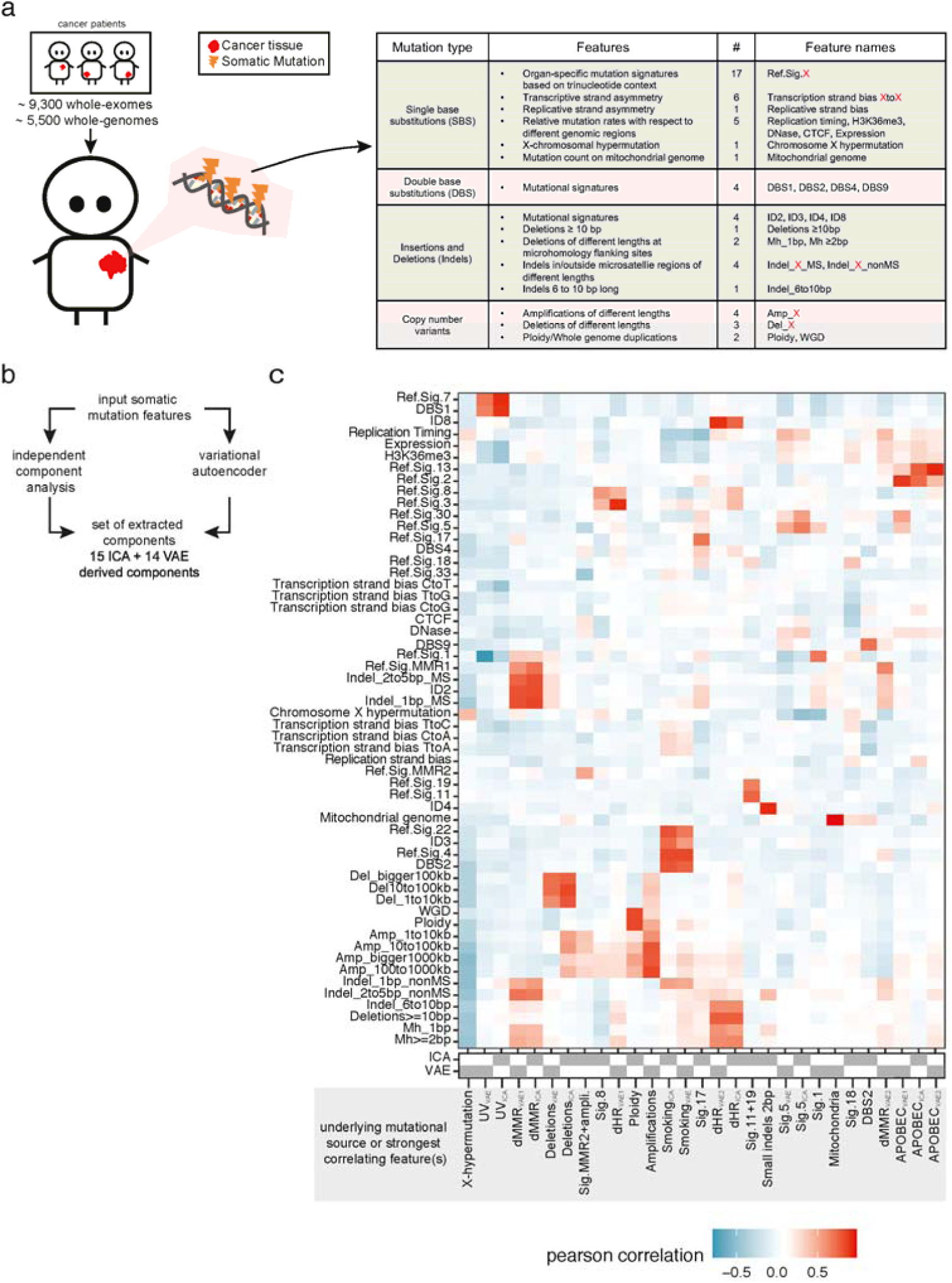
Somatic mutation phenotypes in 15,000 human tumors. **a**, Somatic mutations were extracted from approximately 9,300 whole-exome and 5,500 whole-genome sequenced cancer genomes. **b**, 56 different somatic mutation features were estimated in each cancer genome, covering different types of mutations. **c**, Final set of somatic components was extracted by applying two methods to the input matrix (samples as rows and somatic input features as columns): independent component analysis (ICA) and a variational autoencoder (VAE). 15 ICA-derived and 14 VAE-derived components were extracted. **d**, Overview of extracted somatic components (x-axis) and their Pearson correlation (color code) with the input somatic features (y-axis). Grey strip at the bottom displays whether the component was extracted via ICA or VAE. Components were named based on the underlying mutational process or strongest correlating input feature(s).

To remove the redundancy in these features, we used two different dimensionality reduction techniques – independent component analysis (ICA) and variational autoencoder (VAE) neural network – to deconvolve the (often correlated) mutation features into mutational ‘components’. These components should both better reflect underlying causal mechanisms and increase the statistical power to detect genetic associations by reducing the multiple testing burden. 15 components were derived from the ICA and 14 components from the VAE (see Methods). Thirteen of the 29 components capture known mutagenic mechanisms (Fig. 1c), including UV radiation exposure (UV_ICA_ and UV_VAE_, including CC>TT substitutions), tobacco smoking (Smoking_ICA_ and Smoking_VAE_), deficiencies in MMR (dMMR; dMMR_ICA_, dMMR_VAE1_, and dMMR_VAE2_), deficiency in the repair of DSBs via homologous recombination (dHR; dHR_ICA_, dHR_VAE1_, and dHR_VAE2_), and APOBEC-directed mutagenesis (APOBEC_ICA_, APOBEC_VAE1_, and APOBEC_VAE2_). Many of the components combined different classes of mutational features. For instance, dMMR_VAE2_, has a high correlation with the SNV signature RefSig MMR1^52^, several types of short indels at microsatellite loci and the relative mutation rate with respect to replication timing. The remaining 16 components do not have a known mechanistic cause but can be further described via the features with which they are strongly correlated. For instance, we extracted components covering X-chromosomal hypermutation (X-hypermutation), a component covering mitochondrial SNVs (Mitochondria), and two components related to SNV-signature 5 mutations (Sig.5_ICA_ and Sig.5_VAE_).

### Rare variant association with a combined burden and variance test

To identify genes with rare germline variants that impact somatic mutational processes (Fig. 2a), we defined five different sets of RDGVs using varying approaches and stringency criteria for identifying causal variants, and tested three models of inheritance by also considering RDGVs in combination with somatic loss-of-heterozygosity (LOH)^53^. In total, 15 different models were tested (Fig. 2b top). To increase statistical power, we restricted testing to a set of 892 genes constituting known cancer predisposition genes, DNA repair genes and chromatin modifiers. The combined test SKAT-O^54^, which unifies burden testing and the SKAT variance test^55, 56^, was utilized for testing (Fig. 2b bottom). In brief, the test statistic in SKAT-O is the weighted sum of the test statistic from a burden test and a SKAT test. While in burden testing the variants are aggregated first and then jointly regressed against a phenotype, in SKAT the individual variants in a gene are regressed against the phenotype, and then the distribution of the individual variant score statistics is tested. Importantly, the burden test is more powerful when all RDGVs in a gene are causal, while SKAT is more powerful when some RDGVs are not causal or when RDGVs are causal but with effects in opposite directions^54^. In SKAT-O the parameter contribution of the two tests and corresponds to the smallest reported ρ controls the d p-value^54^, indicating whether the burden or the variance test was used to identify the association.

**Fig. 2.**
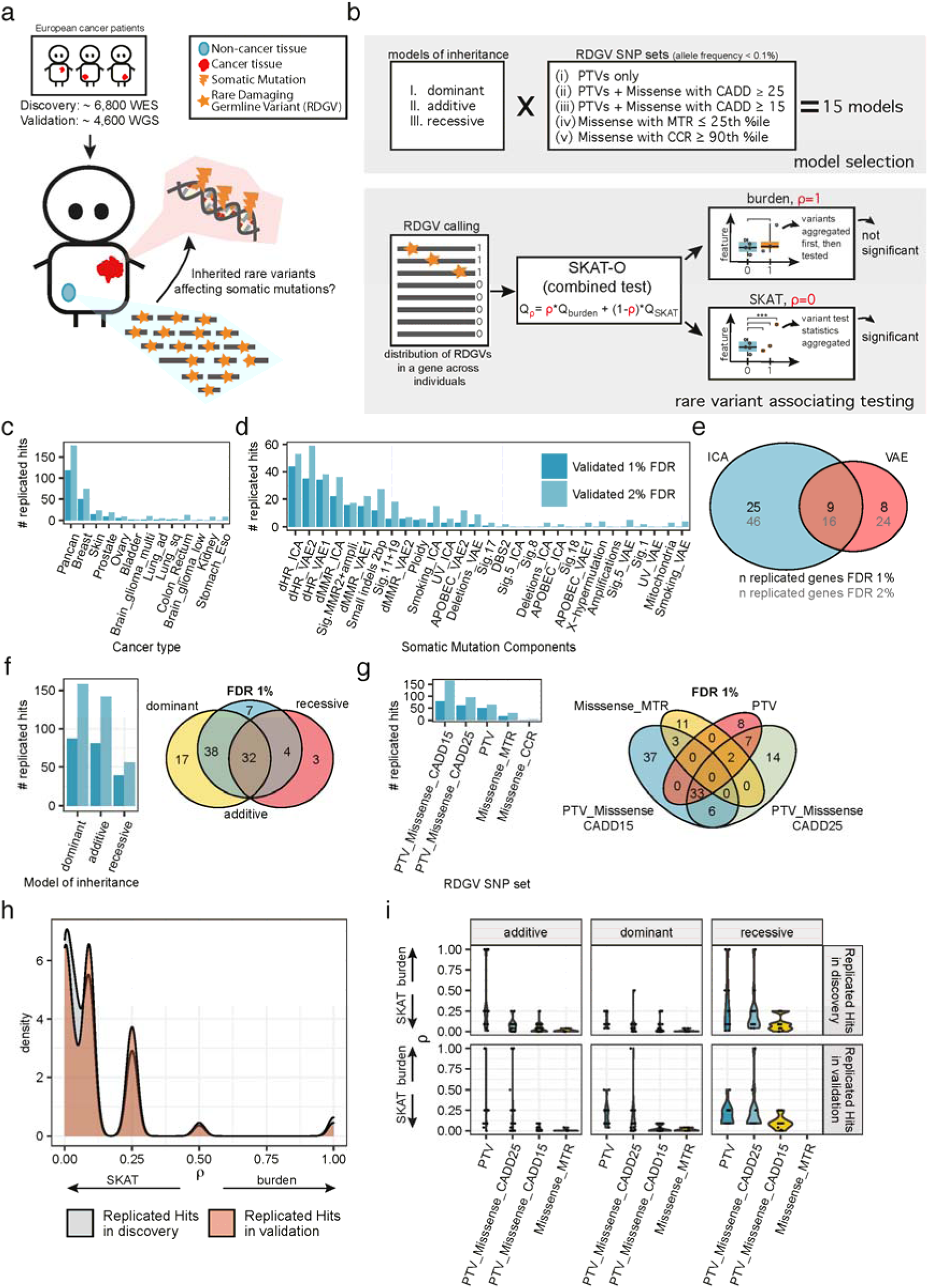
Discovery and validation of rare damaging germline variants (RDGVs) associating with somatic components via a gene-based combined burden and variance test. **a**, Associations were identified in the discovery cohort (TCGA WES) and replicated in the validation cohort (PCAWG + Hartwig WGS). **b**, Associations were tested via 15 models in total, by utilizing 3 models of inheritance and 5 (differently prioritized) SNP sets of rare variants (all with population allele frequency < 0.1 %) (top). The combined test SKAT-O was applied, which calculates a weighted sum between a burden test statistic and the SKAT variance test statistic. When ρ=1, the test reduces to a burden test, and when ρ=0, the test reduces to the variance (SKAT) test. SKAT is more powerful when a fraction of the variants in the SNP set are non-causal, while the burden test has higher power when all variants are causal. **c**, Number of replicated hits at a FDR of 1 % and 2 % across cancer types and **d**, across somatic mutational components. **e**, Overlap of number of genes replicating at a FDR of 1 % and 2 % via the two different dimensionality reduction methods. **f**, Number of replicated hits at a FDR of 1 % and 2 % across models of inheritance (left) and overlap of replicated hits between models at a FDR of 1 % (right). **g**, Number of replicated hits at a FDR of 1 % and 2 % across RDGVs sets (left) and overlap of replicated hits between RDGV sets at a FDR of 1 % (right). **h**, Distribution of ρ values from SKAT-O test (x-axis) for the 207 hits, which replicated at a FDR of 1 %, in the discovery (grey) and validation cohort (red). **i**, Distribution of ρ values from SKAT-O test (y-axis) for the 207 hits, which replicated at a FDR of 1 %, in the discovery (top row) and validation cohort (bottom row), across models of inheritance (columns) and RDGV sets (x-axis).

### 42 genes robustly associated with somatic mutation phenotypes

Testing was performed in the discovery cohort (TCGA) across 6,799 individuals of European ancestry and 12 different cancer types as well as in a pan-cancer analysis (“pancan”) for all 15 models. Genes were only tested via the dominant or additive model when at least 2 individuals carried a RDGV in that gene. For the recessive model, genes were only tested when the gene was biallelically affected in at least 2 samples either by a biallelic RDGV or via a RDGV + LOH (see Methods). In total 594,462 tests were conducted. The tests showed little evidence of inflation when considering models in which at least 100 genes were tested. Overall there was slight deflation (median: 0.78; 1st quartile: 0.55; 3rd quartile: 0.97; max: 2.27) (Extended Data Fig. 1), suggesting conservatively biased test results. Inflated cases were discarded (cut-off at lambda ≥ 1.5; 19 out of 1,909 discarded). We further estimated false discovery rates (FDRs) by randomization. The link between somatic components and individuals was broken down by randomly shuffling the somatic component estimates of the individuals within each cancer type. Empirical FDRs were estimated by comparing the observed p-value distribution against the random one (see Methods and Extended Data Fig. 2). As an additional negative control, we considered a random set of genes, comparing the number of replicated hits at a certain empirical FDR with the random gene set to the number with our candidate gene list (Extended Data Fig. 2). It should be noted that this yields a conservative upper limit since the random gene lists may also include genes which affect somatic mutation processes.

In total, we identified 6,488 associations (out of 591,302 tests) in the discovery phase at an empirical (randomization-based) FDR of 1% (Extended Data Fig. 3). Out of the 6,488 hits, 3,807 had a sufficient number of RDGVs in the matching cancer type (see Methods) to allow re-testing in an independent validation cohort (merged PCAWG and Hartwig) in the matching cancer type, consisting of 4,683 patients of European ancestry. 207 associations replicated in the validation cohort at an empirical FDR of 1 %, covering 42 individual genes, 15 mutational components, 46 unique gene-cancer type pairs, and 65 unique gene-cancer type-component combinations (Fig. 3). We also checked the number of replicated associations at a more permissive FDR of 2 %. At an FDR of 2 %, 12,480 hits were detected in the discovery cohort, 7,290 hits were re-tested in the validation cohort, out of which 356 associations were replicated covering 86 individual genes, 24 mutational components, 105 unique gene-cancer type pairs, and 140 unique gene-cancer type-component combinations (Extended Data Fig. 4). Notably, 7 genes associated across more than one cancer type, of which 3 (*BRCA1*, *EP300*, *MTOR*) associated with the same somatic mutational component across two different cancer types (Extended Data Fig. 5).

**Fig. 3.**
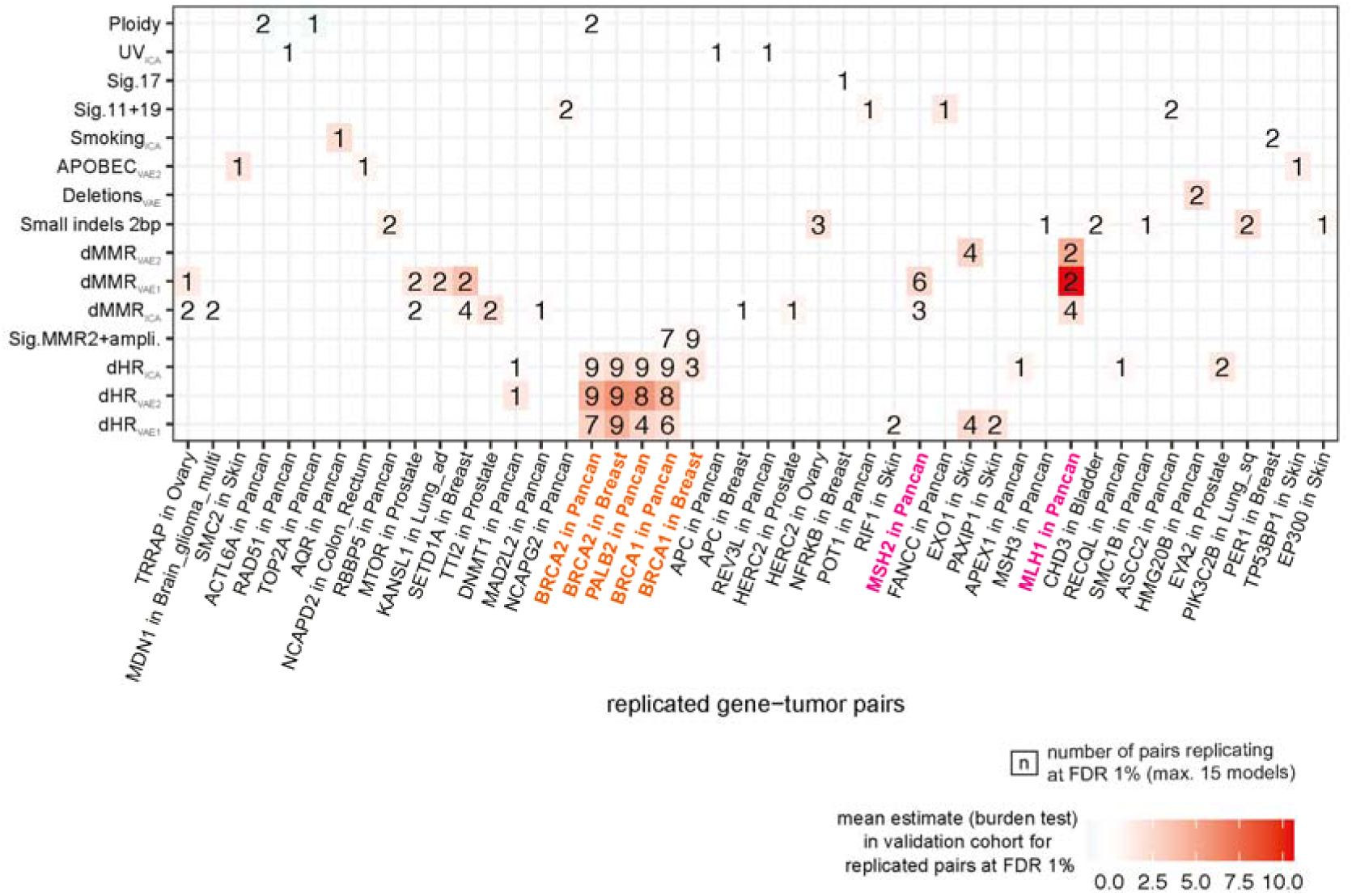
Overview of replicated hits at a FDR of 1%. Showing gene-cancer type pairs (x-axis), the corresponding somatic mutational component (y-axis), and the number of times they replicated at a FDR of 1 % (maximum of 15 models for each gene-cancer type-somatic component tuple). Color represents the mean estimate of the regression coefficient from burden test for all replicated hits at a FDR of 1 %, for the respective gene-cancer type-somatic component combination. Previously associated dHR genes in orange and dMMR genes in pink. Genes on the x-axis were ordered based on hierarchical clustering results using DepMap CRISPR-derived genetic fitness (Chronos) scores (see Supplementary).

**Fig. 4.**
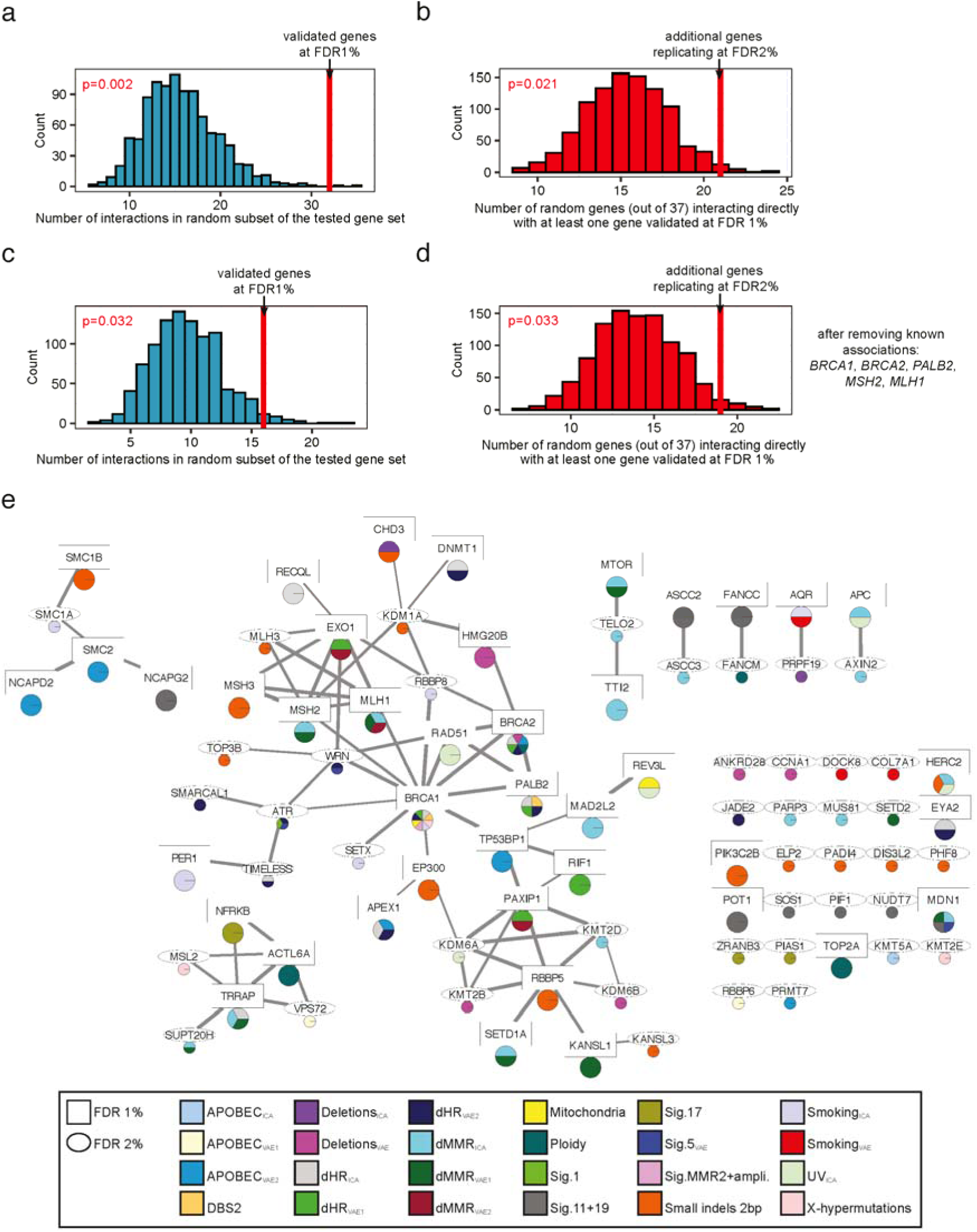
Network analysis supports the role of rare germline variation in somatic mutational processes. All panels in this figure were generated using physical interactions from the STRING database having a combined score ≥ 80%. **a**, Number of physical interactions in a random subset of the tested gene set (controlled for interaction node degree) (x-axis). Red line shows the number of interactions within genes which replicated at a FDR of 1 %. **b**, Number of randomly selected genes from the tested gene set interacting with at least one gene, which replicated at a FDR of 1 % (x-axis), (controlled for interaction node degree). Red line shows the number of genes, out of the ones which additionally replicated at a FDR of 2 %, interacting with at least one gene replicating at a FDR of 1 %. **c**, Same as in panel a, after excluding known genes from the analysis (*BRCA1*, *BRCA2*, *PALB2*, *MSH2*, and *MLH1*). **d**, Same as in b after excluding known genes from the analysis (*BRCA1*, *BRCA2*, *PALB2*, *MSH2*, and *MLH1*). **e**, Visualisation of physical interactions between proteins for genes replicating at a FDR of 1 % (square) and genes replicating at a FDR of 2 % (ellipse). Color code in pie chart shows the somatic components the corresponding gene was associated with (bottom panel). Line width corresponds to combined (experimental, database, and text mining) STRING physical interaction score.

**Fig. 5.**
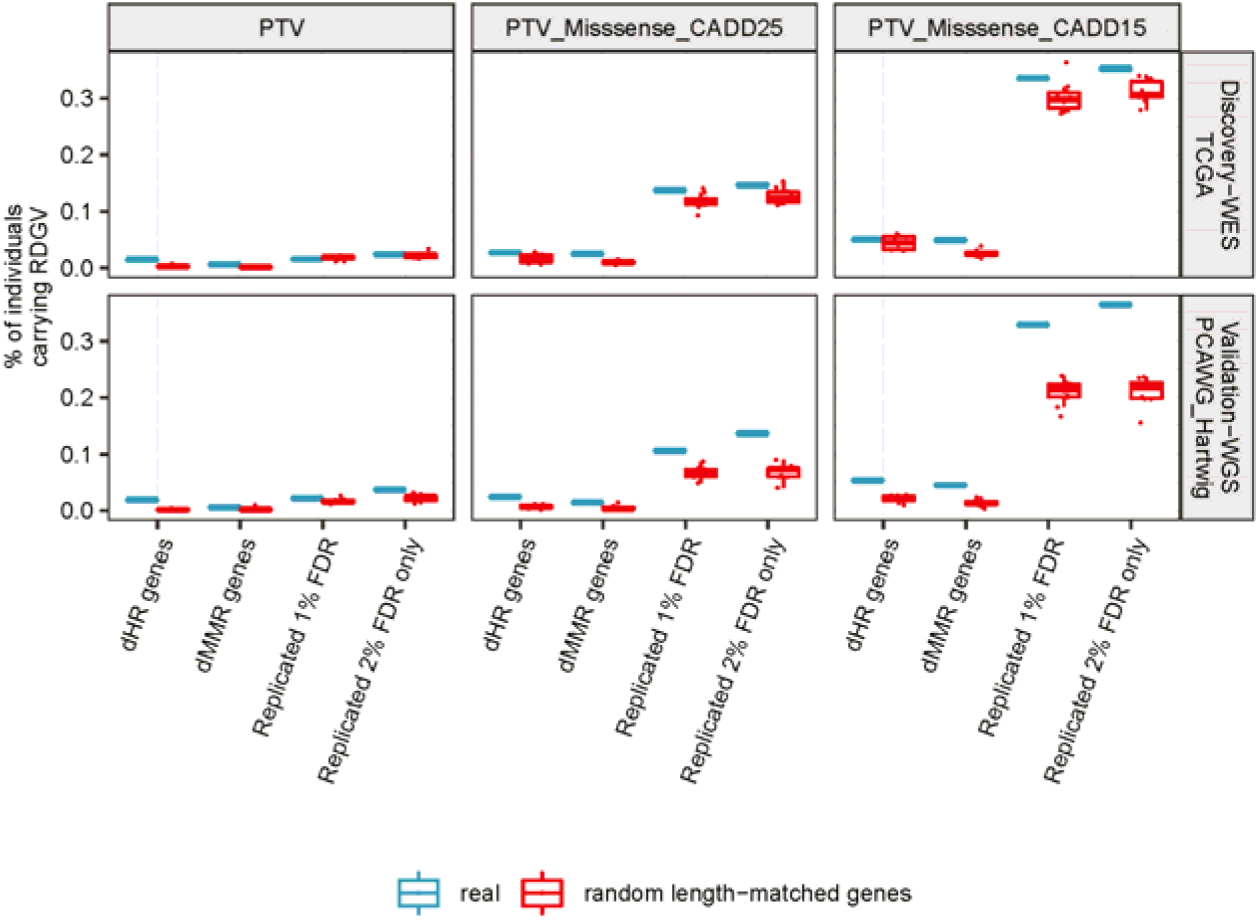
Frequency of RDGVs across cohorts. Showing the frequencies of RDGVs within the individuals (y-axis) of the discovery cohort (TCGA-WES) and validation cohort (PCAWG+Hartwig-WGS) (rows) across different RDGV sets (columns) for different gene sets (x-axis). Known dHR gene set includes *BRCA1*, *BRCA2*, *PALB2*, and *RAD51C*, known dMMR gene set includes *MSH2*, *MSH6*, *MLH1*, and *PMS2*, the replicated 1 % FDR set includes all genes replicating at a FDR of 1 % after excluding known dMMR and dHR genes, and the replicated 2 % FDR only set includes all additional genes which replicated at a FDR of 2 %. Color code for the real gene sets (blue) and length-matched, randomly selected protein-coding gene sets (red). Random selection for length-matched protein-coding genes was performed 10 times, and distribution shown in boxplot.

At an FDR of 1%, most of the replicated hits were identified in the pan-cancer analysis (57 %), followed by breast cancer (24 %), skin cancer (7%), and prostate cancer (4 %) (Fig. 2c), reflecting differential sample sizes between cancer types (Extended Data Fig. 3e). Furthermore, approximately half of the components (15 out of 29) were associated with at least one gene-cancer type pair (Fig. 2d). Many replicated hits were associated with features related with dHR (dHR_ICA_: 21 %, dHR_VAE1_: 17 %; dHR_VAE2_: 16 %), followed by dMMR (dMMR_ICA_: 11 %; dMMR_VAE1_: 7 %), consistent with well-established roles of HR and MMR failures in accelerating mutation rates in tumors^38^. Notably, 25 genes were only identified via an ICA derived component, while 8 genes were only identified via a VAE derived component (Fig. 2e), suggesting a complementary role of the two approaches to summarize mutation processes.

Many of the replicated associations were identified via the dominant (42 %) and (39 %) additive model (Fig. 2f), suggesting that heterozygous variants can alter mutation rates in humans, as was suggested for a model organism^50^. The comparatively lower number of replicated hits of the recessive model can be largely attributed to the fact that RDGV combined with somatic LOH events are considerably less frequent and thus associations could not be tested for many genes (only 4 % of the 591,302 tests performed in the discovery phase came from the recessive model). Considering the proportion of replicated hits to the number of re-tested hits, the validation rate was ∼2.5 times higher via the recessive model (Extended Data Fig. 3e), which was expected since many DNA repair genes are believed to be haplosufficient^57^.

We further considered the number of replicated associations using different approaches and stringency thresholds for declaring a variant to be pathogenic. The highest number of hits replicated using the more permissive thresholds, using protein-truncating variants (PTVs) + missense variants at a CADD^58^ score ≥ 15 (79/207, 38%), followed by PTVs + missense variants at a CADD score ≥ 25 (62/207, 30%) and PTVs only (50/207, 24 %) (Fig. 2g). This suggests that some missense variants that were assigned a lower pathogenicity score – likely due to difficulties in assessing variant pathogenicity *in silico*^59^– can nonetheless bear on somatic mutation phenotypes. We further tested by only considering RDGVs in conserved gene segments (via “constrained coding regions”^60^ and “missense tolerance ratio”^61^ methods), however this yielded few replicated hits (Fig. 2g). It should be noted, however, that some hits were only identified when using the PTV-only set and were not recovered in more permissive RDGV sets.

In summary, with regards to the model of inheritance, RDGV set, and component (mutational process) extraction method, there was no single best model and most models added unique associations to the results.

### More permissive thresholds for variant pathogenicity increase the utility of a variance test over a burden test

The SKAT-O test we employed combines burden testing and a variance test component (SKAT)^54^. Examining the SKAT-O parameter ρ for the 207 validated d that most hits hits, in both the discovery and the validation cohort, revealed that most hits replicated via the variance test (ρ < 0.5 in 393/414 tests) (Fig. 2h). The variance test is the more powerful test of the two when many variants in the tested set are not causal^54^. We hypothesized that a common reason why allegedly pathogenic RDGVs would not be causal is because of inaccurate prediction of damaging variants by *in silico* predictors^62^. If so, at the more stringent settings more hits would replicate via the burden test (which has higher power when many variants in the set are causal), while at the less stringent settings more hits would replicate via the variance test (which is robust to inclusion of non-causal variants). Indeed, several hits replicated via the burden test when using the most stringent RDGV set (PTVs only; Fig. 2i), including *MLH1*, *BRCA1*, and *BRCA2*. For the more permissive RDGV sets, the number of hits replicating via the burden test decreased and all of the replicated hits had a ρ lower than 0.25 (meaning, they used nearly exclusively the variance component) for the RDGV set including missense variants at CADD ≥ 15. The positive control genes *BRCA1* and *BRCA2* licated in the PTV+missense CADD ≥ 15 RDGV set, but with a ρ of 0 (variance test exclusively used), suggesting that this variant set included many non-causal variants.

In summary, many hits were recovered even with more permissive RDGV sets by utilizing the combined testing approach of the SKAT-O method, suggesting the variance (SKAT) component can partially compensate for the inaccuracy of the *in silico* predictors. Most of the replicated hits would not have been identified by use of classical burden testing in a data set of this size.

### Novel genes associating with defects in homologous recombination

Within the set of 207 replicated associations at an FDR of 1%, 117 (57 %) involved associations of *BRCA1*, *BRCA2*, and *PALB2* with various mutational components associated with dHR (Fig. 3), consistent with the known role of these genes in the error-free repair of DSBs. All three genes associated with features of defective HR, such as deletions at microhomology-flanked sites (dHR_ICA_ and dHR_VAE2_) and SNV signature 3 mutations (dHR_VAE1_). In addition, *BRCA1*, but not *BRCA2*, associated with component Sig.MMR2+ampli., reflecting an increased number of amplification events. This is in accordance with a recent report, in which *BRCA1*-type dHR vs. *BRCA2*-type dHR were differentiated via the presence of duplication events^40^.

We also detected additional genes associating with these dHR mutational components. In skin cancer, *PAXIP1*, *EXO1,* and *RIF1* associated with dHR_VAE1_, the component correlating with SNV signature 3 mutations. In support of this, *PAXIP1* and *RIF1* have been implicated in the repair of DNA DSBs^63–65^ and interact with each other^66^. Thus, these associations suggest that individuals carrying damaging variants in either gene have an increase in signature 3 mutations, potentially reflecting a downstream effect of disrupted DSB repair. Additionally, *EXO1* knockout in a cell line model^67^ was reported to result in a mutational signature correlating with signatures 3 (Pearson R=0.71) and 5 (R=0.71), supporting our association observed in tumors.

Furthermore, we identified pan-cancer replicated associations of *APEX1*, RECQL, and DNMT1 with dHR_ICA_ (with *DNMT1* additionally associating with dHR_VAE2_). These associations with a microhomology deletion mutation phenotype are diagnostic of an increased activity of the microhomology-mediated end joining (MMEJ), a highly error-prone DSB repair pathway, suggesting that variants in these genes may disrupt normal functioning of the less error-prone HR and/or NHEJ pathways.

Five additional genes (*ATR*, *JADE2*, *SMARCAL1*, *TIMELESS*, and *WRN*) were identified at a more permissive threshold, associating with at least one dHR-related component (dHR_ICA_ and/or dHR_VAE2_). Notably, ATR and WRN physically interact with BRCA1 (Fig. 4e) and play known roles in repair of DSBs^68–70^, which would support these associations. In particular, pathogenic recessive variants in WRN cause Werner syndrome^71^ and it has been suggested that the WRN helicase is crucial for the repair of MMR-induced DSBs^72, 73^. Additionally, SMARCAL1 and TIMELESS directly interact with ATR (Fig. 4e).

Our analyses therefore replicate well-known associations between rare inherited variants in HR genes and somatic mutational components, as well as identifying new associations with additional genes.

### *MTOR* and interacting protein variants associate with mismatch repair phenotypes

In the context of Lynch syndrome, germline variants in *MLH1*, *MSH2*, *MSH6* and *PMS2*^74^ affect somatic mutation patterns via impairment of the DNA mismatch repair pathway, observed as microsatellite instability (MSI, indels at simple DNA repeats)^75, 76^. MSI was also later associated with mutational signatures derived from SNVs^18^, as well as with a ‘redistribution’ of mutations across replication timing domains^22^. In accordance with this, we detected associations of RDGVs in *MLH1* and *MSH2* with multiple dMMR-related components i.e. those having a high contribution of small indels at microsatellite regions (dMMR_ICA_ and dMMR_VAE1_), and with SNV-derived signature MMR1 mutations and replication timing (dMMR_VAE2_; for *MLH1*).

Beyond the known Lynch Syndrome genes, we also discovered associations between variation in *EXO1*, which has an established role in MMR^77^ and increases the frequency of 1 bp indels when inactivated in cultured cells^78^, and dMMR_VAE1_ and dMMR_VAE2_. However, *EXO1* also associated with dHR-related components, suggesting a more pleiotropic role for the encoded exonuclease in shaping somatic mutational processes in human tumors. Consistent with the association with dHR components, it was reported in yeast as well as human cell lines that *EXO1* processes DSB ends^79^ and is required for the repair of DSBs via HR^80^.

Multiple other genes were associated with dMMR-directed phenotypes (all associated with dMMR_ICA_ and dMMR_VAE1_), including the chromatin modifying enzyme genes *TRAAP* in ovarian and *SETD1A* in breast, and the major growth signalling gene *MTOR* in prostate cancer (and in stomach+esophagus cancer with dMMR_VAE1_ only at a FDR of 2 %). Additionally, *TTI2* in prostate, *APC* in breast, *MAD2L2* in pan-cancer, *HERC2* in prostate, and *MDN1* in brain cancer associated with mutation component dMMR_ICA_. There is additional evidence supporting these associations for some of these genes from prior studies. *MTOR* was identified as one of four genes that regulate MSH2 protein stability^81^. Thus, a possible mechanism explaining the identified association of *MTOR* with dMMR-linked components could be a decreased stability of MSH2 leading to dMMR and consequently, an increased number of indels. A similar mechanism could be speculated for *TTI2*, which binds *MTOR* via the TTT complex (*TELO2*-*TTI1*-*TTI2*) and is important for mTOR maturation^82^. This hypothesis is further supported by *TELO2* associating with the same component (dMMR_ICA_) in kidney cancer at a more permissive FDR of 2% (Extended Data Fig. 4). Furthermore, *SETD2* associated in colorectal cancer with dMMR_VAE1_ at a FDR of 2 %. It has been shown in previous studies^23^, including in cancer genomes^24^, that the encoded methyltransferase SETD2 regulates MMR activity by recruiting the MSH2-MSH6 complex to H3K36me3 marked regions.

Taken together, we recovered known associations of MMR genes with somatic mutational patterns and identified additional genes where germline variants are associated with MMR phenotypes, suggesting that a broad network of genes cooperates to maintain MMR efficiency in human cells.

### MSH3 and additional genes associate with a distinct dMMR phenotype

Interestingly, we identified associations between RDGVs in several genes and a somatic mutational component (Small indels 2 bp) that reflects indels of a size of 2 bp and longer, which is in contrast to the predominantly 1 bp long indels caused by dMMR. Furthermore, this component does not have any contribution from SNV features, indicating that it is specifically capturing indels (Fig. 1c and Supplementary Fig. 6). Among others, the MMR gene *MSH3* associated with this component in the pan-cancer analysis. In contrast to the DNA mismatch repair genes *PMS2*, *MLH1*, *MSH2,* and *MSH6*, germline variants in *MSH3* have not been identified in patients with Lynch syndrome, even though they were reported to increase cancer risk^37^. The MSH2-MSH3 complex has a role in repairing insertion/deletion loops rather than for base-base mismatches^83, 84,85^.

This is in contrast to the MSH2-MSH6 complex, which repairs base-base mismatches and indels shorter than 2 nucleotides^86, 87^. These prior mechanistic studies support our association and suggest that loss of *MSH3* in cancer cells results in an increased rate of accumulation of indels of 2 bp and longer. Other genes associating with this component were *CHD3* in bladder cancer, *HERC2* in ovary cancer, *PIK3C2B* in lung squamous cell cancer, *EP300* in skin cancer (and breast cancer at a FDR of 2 %), *RBBP5* in pancan, and *SMC1B* in pancan. Additionally, *MLH3* associated with the same component at an FDR of 2 %. The MLH3 protein is a paralog of MLH1 that interacts with other MMR proteins (Fig. 4e) and was previously associated with microsatellite instability^88^.

Overall, we detected associations between germline variants in *MSH3* and several other genes and somatic indels of at least 2 bp, suggesting a causal role for MSH3 variants in a specific subtype of MMR failure which does not markedly increase SNV rates.

### Genes associating with a somatic feature enriched in brain and liver cancer

Beyond the dHR and dMMR-related components, the component associated with the largest number of genes was component Sig.11+19, which is enriched for SNV signatures RefSig 11 and 19^52^ (Fig. 2d). This component is enriched in brain and liver cancers (Supplementary Fig. 7). Signature 11 has been reported to be enriched in brain cancers, associated with temozolomide treatment^18^, and is similar to the signature which results from the treatment with the DNA methylating agent 1,2-Dimethylhydrazine^89^. The cause of signature 19 is unknown and it has been mostly identified in brain, liver and blood cancers^52^. At a FDR of 1 %, the genes *ASCC2*, *FANCC*, *NCAPG2* and *POT1* associated with this component in the pan-cancer analysis, as do *NUDT7*, *PIF1*, and *SOS1* at a more permissive 2% FDR. POT1 and PIF1 interact with each other^90^ (Extended Data Fig. 6e) and both have functions in telomere maintenance^91, 92^, but we did not detect any correlation between this component and reported telomere features^93^ (Supplementary Fig. 16).

### Variants in APEX1 associate with increased level of APOBEC-directed mutagenesis

We discovered and replicated associations between *APEX1* and three different somatic components. *APEX1* encodes for a purinic/apyrimidinic (AP) endonuclease that cleaves at abasic sites, which can be formed spontaneously or during base excision repair pathway by a DNA glycosylase^94^. At a FDR of 1 %, *APEX1* associated with dHR_ICA,_ in pancan and at a FDR of 2 % it associated with dHR_VAE2_ in pancan and with APOBEC_VAE2_ in stomach/esophagus cancer. The somatic components dHR_ICA_ and dHR_VAE2_ are enriched for deletions at microhomology-flanked regions. Prior studies showed that the encoded protein APE1 protein plays a role in the repair of DSBs and that depletion of APE1 leads to an decrease of HR-directed repair^95^, suggesting a higher reliance on alternative pathways.

The APOBEC_VAE2_ component is enriched for SNV signature 13 (C>G) mutations^18^. These can be formed when the APOBEC-induced uracil is excised via the uracil-DNA glycosylase UNG and a cytosine is inserted opposite the abasic site by the mutagenic translesion polymerase REV1^96^. Conceivably, a mechanism underlying the higher burden of C>G mutations in tumors of individuals with inherited damaging variants in *APEX1* could be due to a decreased activity leading to a slower repair of the abasic site and consequently, a preference for lesion bypass via the error-prone REV1.

### Network analysis reinforces the role of rare germline variants in somatic mutation processes

The previously known dHR genes encode proteins that physically interact as part of the same protein complexes^97^. Similarly, the products of the known dMMR genes also physically interact^98^. We used protein-protein interactions curated in the STRING^99^ database to test whether the genes identified as having rare germline variants associating with somatic mutational phenotypes also encode physically interacting proteins. Such ‘guilt by association’ network analysis has been used to support associations between somatic mutations and cancer^100, 101^ and between common variants and disease phenotypes^102^ but has not yet been widely adopted for the analysis of rare variants.

We first considered genes associated with somatic mutation phenotypes at a FDR of 1%. These genes are strongly enriched for encoding proteins with physical interactions (Fig. 4a; median difference=17 and P=0.002 by randomisation, controlling for interaction node degree). This also held true after removing genes with previously reported associations (Fig. 4c; median difference=7 and P=0.032 by randomisation).

Secondly, we considered the 44 genes with moderate statistical support of association with somatic mutation phenotypes (those replicating at a FDR of 2%). 21 of the encoded proteins interact with at least one of the proteins encoded by the more stringent FDR 1% genes. This is again higher than expected by chance (Fig. 4b; median difference=6 and P=0.021 by randomisation), further prioritising these 21 genes for additional study. This also held true after removing previously known genes (Fig. 4d; median difference=5 and P=0.033 by randomisation). Similar results were seen using the HumanNet gene network^90^ that incorporates many data sources to predict functionally-related genes (Extended Data Fig. 6).

Thus, genes with replicated associations with somatic mutation phenotypes preferentially encode proteins that physically interact in cellular networks with genes replicating at a more permissive FDR also often connected to the same sub-networks, illustrating the potential for network-based analyses to provide supporting evidence in rare variant association studies.

### Prevalence of damaging germline variants in genes associated with somatic mutational phenotypes

To better estimate the contribution of RDGVs to differences in somatic mutational processes, we counted how many individuals in our dataset had certain RDGVs and compared this to randomly selected protein-coding genes while controlling for covered gene length (Fig. 5). Considering known mutator genes, 44 individuals (0.6 %) had PTVs in Lynch syndrome dMMR genes (*MSH2*, *MLH1, MSH6, PMS2*), and 100 (1.5 %) had PTVs in in dHR genes (*BRCA1*, *BRCA2*, *PALB2*, *RAD51C*) in the discovery cohort (TCGA). Considering only the newly associated genes, 107 individuals (1.6 %) had a PTV in genes that replicated at a FDR of 1 %, and 166 (2.4 %) in genes which replicated at a FDR of 2 %. A similarly high prevalence of damaging variants in newly-discovered genes, relative to known mutator genes, was seen in prioritized missense variants, via the CADD score at stringent (≥ 25) and permissive thresholds (≥ 15; Fig. 5). Additionally, when comparing this with prevalence of deleterious variants in control sets of length-matched genes, there is an excess of damaging missense variants in the known dHR and dMMR genes as well as in the newly-discovered genes at 1% and 2% FDR thresholds (Fig. 5).

Taken together, these results suggest that the novel candidate mutator genes are affected by deleterious variants in a higher fraction of the population of cancer patients than the known human germline dMMR and dHR genes.

## Discussion

We have shown here that rare inherited variants in diverse genes associate with different mutational processes. Our approach incorporated a variance-based test via SKAT-O^54^, two different dimensionality reduction algorithms to extract somatic mutation patterns, the usage of different *in silico* variant prioritization tools^58, 60, 61^, and the use of different models of inheritance for association testing. This experimental design allowed us to identify multiple new replicating associations between genes and somatic mutation phenotypes.

Most of the associations we identified were replicated only via the variance-based test SKAT, which suggests that variants predicted to be damaging still contain many non-causal variants. More accurate variant effect prediction tools should further increase the power of these kinds of analyses^62, 103, 104^. We also found that using two techniques to derive informative somatic mutation components identified more replicated associations than using either approach alone. This is consistent with findings in other fields, where different algorithms have also been found to capture complementary information, for example in gene expression analysis^105^ and identifying genetic variants from genomic data^106^.

We identified novel genes associating with dHR-related repair (e.g. *RIF1*, *PAXIP1*, *WRN*, *EXO1*, and *ATR*) and with components connected to dMMR (e.g. *MTOR*, *TTI2*, SETD2, *EXO1, MSH3,* and *MLH3*). Several novel associations are supported by strong evidence from prior studies such as *EXO1* with dHR^79, 80^ and dMMR^67, 77, 78^, *SETD2* with dMMR^23, 24^ and *MSH3* with a different form of dMMR^78, 83, 84^ On top of the associations with dHR-and dMMR-related components, we also identified an association of *APEX1* with APOBEC-directed mutagenesis (as well as dHR), and additionally several genes associating with a component enriched in brain and liver cancers with an unknown underlying mechanism. ‘Guilt by association’ network analysis has not yet been widely adopted in rare variant association studies but we found that it was useful for both connecting high stringency replicating genes to each other and for connecting lower confidence hits to the high confidence genes. These interactions are useful for prioritising the newly associated genes and provide specific hypotheses connecting them to known germline mutator genes.

Interestingly, the genetic associations distinguish between two different dMMR mutational phenotypes. Firstly, the common dMMR signature, enriched for 1 bp indels and the SNV-signature MMR1; these associations involved e.g. the Lynch syndrome genes *MSH2* and *MLH1*, and some additional genes e.g. *MTOR,* and *SETD2*. Secondly, a distinct set of associations involved a mutational component enriched for 2 bp and longer indels, but did not encompass a notable increase in SNVs, e.g. involving the core MMR gene *MSH3*, and additionally *MLH3*, *EP300*, and *PIK3C2B*.

This study has some limitations resulting from technical factors. The design is likely to result in a conservative bias in the number of replicated hits, because the discovery and validation cohorts were based on different sequencing technologies (WES versus WGS, respectively). WES data yields more noisy somatic mutation features, as it covers ∼2 % of the genome and some features (e.g. replicative strand asymmetry, mutations at CTCF/cohesin binding sites) are measurable at few loci and so enrichments are difficult to estimate due to low mutation counts. Moreover the power to call germline variants at certain loci may be different for WGS and WES data. The TCGA WES data also has batch effects stemming from the different sequencing centers and sequencing technologies^107, 108^. To offset this risk, we only extracted germline variants from regions with enough coverage in each of three sequencing centers as previously^53^. This limited the number of RDGVs extracted, and thus potentially also the number of discoveries.

In order to increase the sample size and thus power, we combined the cancer cohorts that contained both primary and metastatic cancers, as well as treatment-naïve and pretreated. Similarly, in the pan-cancer analyses, we aggregated data from all cancer types, with the result that the distribution of cancer types between the discovery and validation cohort was somewhat different. It is possible that some hits did not replicate due to these differences in cancer type composition.

Our initial set of somatic mutational features was largely motivated by recent reports^14, 16–19, 22, 23, 26–29, 31, 32, 39, 40, 42^. Consideration of additional, complementary features could identify additional associations in future studies. Lastly, our analysis was performed on samples with European ancestry since this was the most numerous group and including sequencing data from more diverse populations is also likely to identify additional associations.

In conclusion, our findings highlight the role of rare inherited germline variants in shaping the mutation landscape in human somatic cells, leading to variability in somatic mutagenesis between individuals. The results support observations from genetic screens in model organisms suggesting that mutational processes can be affected by variation in diverse genes ^50,51^and suggest that low mutation rates in human somatic cells are hard to maintain. Cooperation between many genes is required to guard against genomic instability: the canonical mutator genes (particularly MMR, HR genes) are embedded in a network of regulators and supporting genes required for optimal functioning of the DNA repair systems.

In the future, larger sample sizes with WGS data and better variant pathogenicity prediction tools will enable higher-powered association studies, further elucidating the potentially very numerous set of genes which determine human somatic mutation rates. The identification of additional genes altering human mutation processes may have important implications for understanding, preventing and treating cancer and other somatic mutation-associated disorders.

---

**Extended Data Fig. 1.**
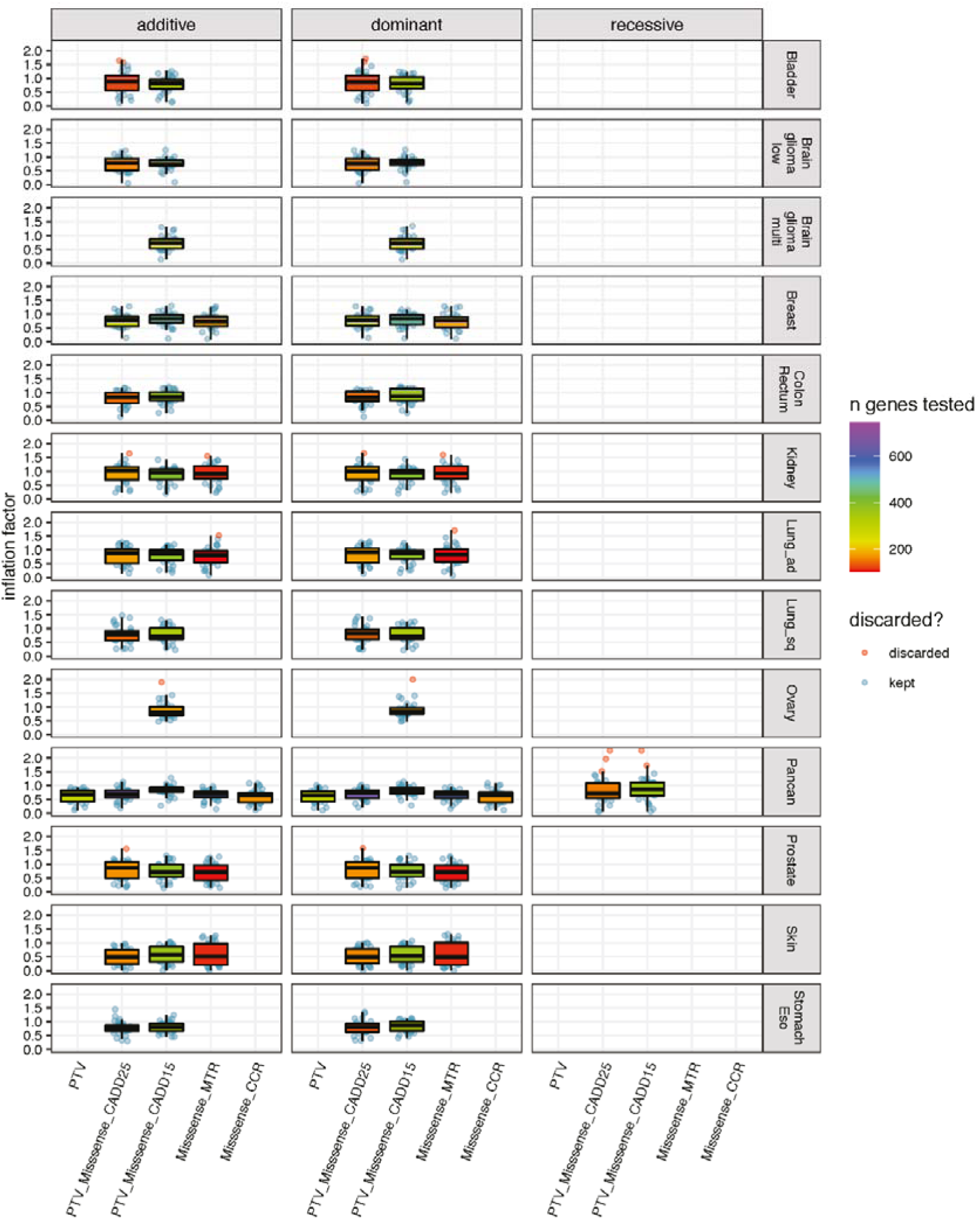
Inflation analysis. Overview of inflation factors (y-axis) across RDGV sets (x-axis), across cancer types (rows), and across models of inheritance (columns). Color code for box plots illustrates the number of tested genes for the respective scenario. Inflation factors were only calculated when at least 100 genes were tested, and inflation factors ≥ 1.5 were

**Extended Data Fig. 2.**
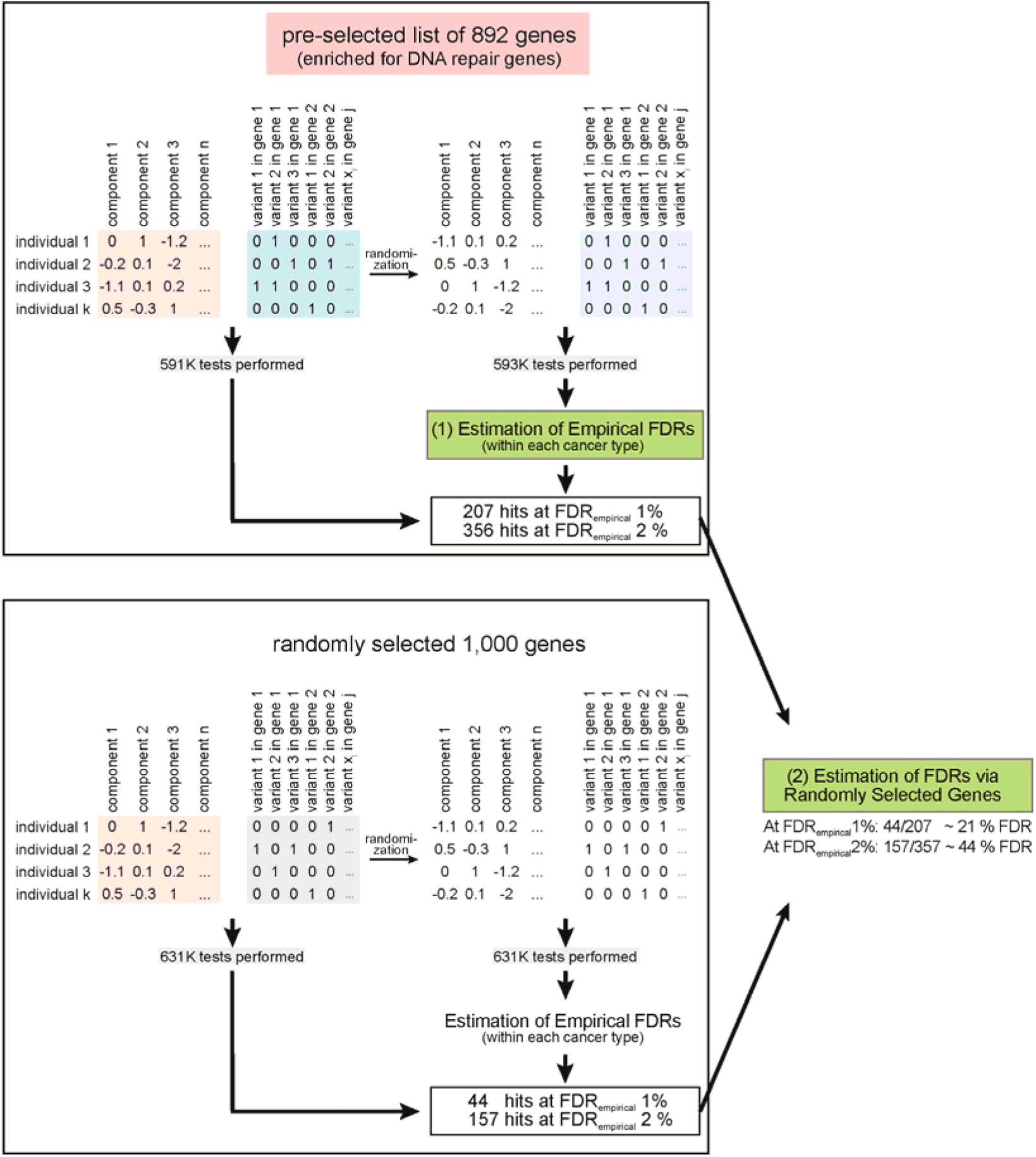
Estimation of false discovery rates. Schematic illustration of the approach. Firstly, testing was performed using the pre-selected 874 genes. Then, randomization was performed shuffling the rows within cancer types, effectively breaking down the link between individuals and somatic components. Testing was performed with the randomized somatic component matrix as well and empirical FDRs were calculated based on the randomization for each cancer type (top half of plot). The same approach was repeated with a random set of 1,000 genes after excluding the pre-selected gene list and any gene interacting with a gene from the pre-selected gene list (bottom half of plot). The number of genes replicating via the randomly selected list of genes at a specific FDR was divided by the number of genes replicating with the pre-selected list to get a conservative FDR estimate.

**Extended Data Fig. 3.**
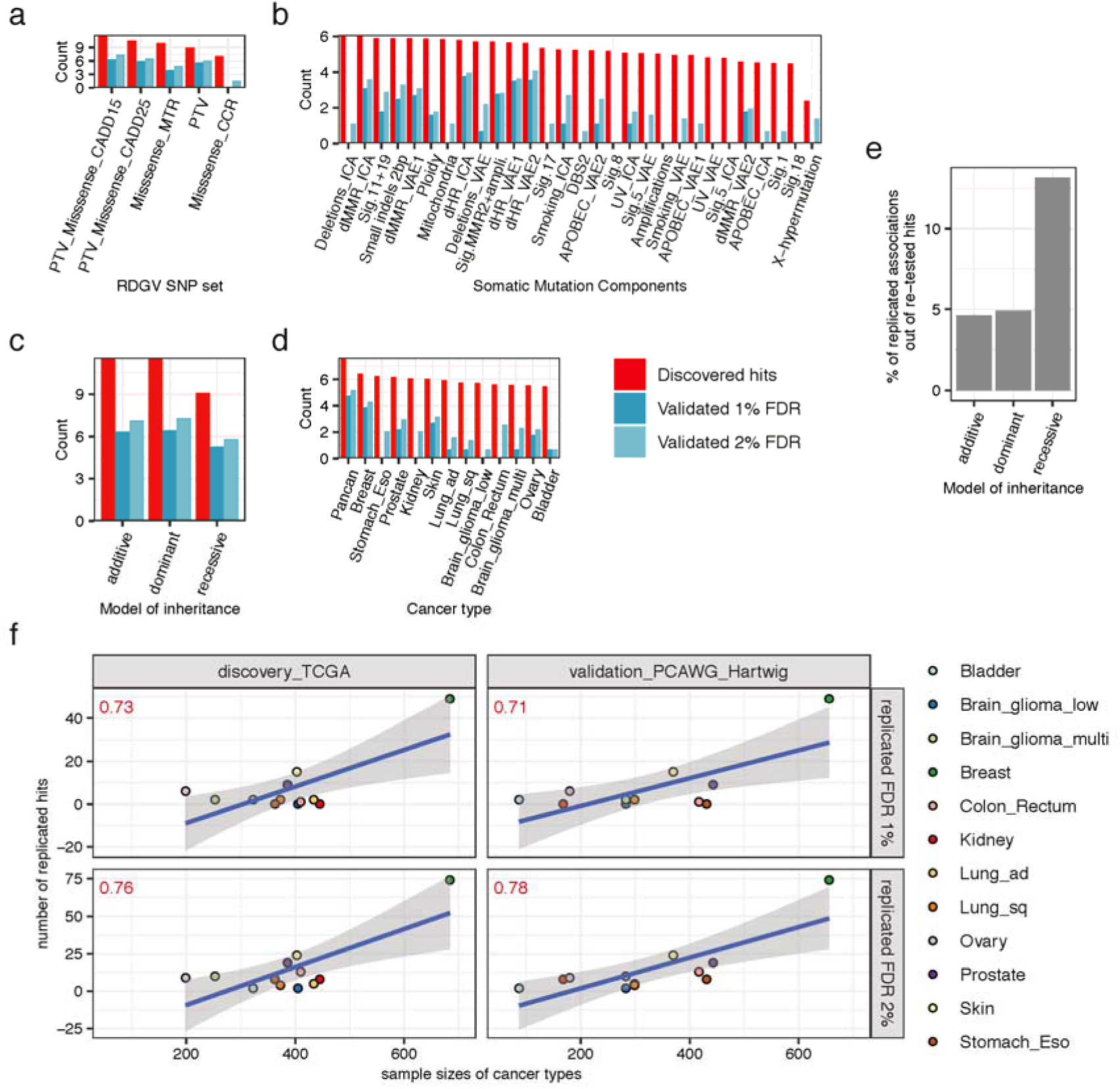
Overview of number of discovered and validated hits. **a**, Number of discovered hits, number of replicated hits at a FDR of 1 % and number replicated hits at a FDR of 2 % across RDGV sets, **b**, somatic components, **c**, models of inheritance, and **d**, cancer types. Log2 counts shown on the y-axis for panels a-d. **e**, Amount of replicated hits out the re-tested discovered hits at a FDR of 1 % across different models of inheritance. **f**, Number of replicated hits (y-axis) versus sample sizes of the corresponding cancer types in which they replicated (x-axis). Columns represent the two cohorts, and rows the applied FDR. Color code for the different cancer types. Pearson correlation shown on the top left corner in red and linear regression fitted through each plot (blue line). Shaded band illustrating 95 % confidence interval. Pancan analysis was excluded.

**Extended Data Fig. 4.**
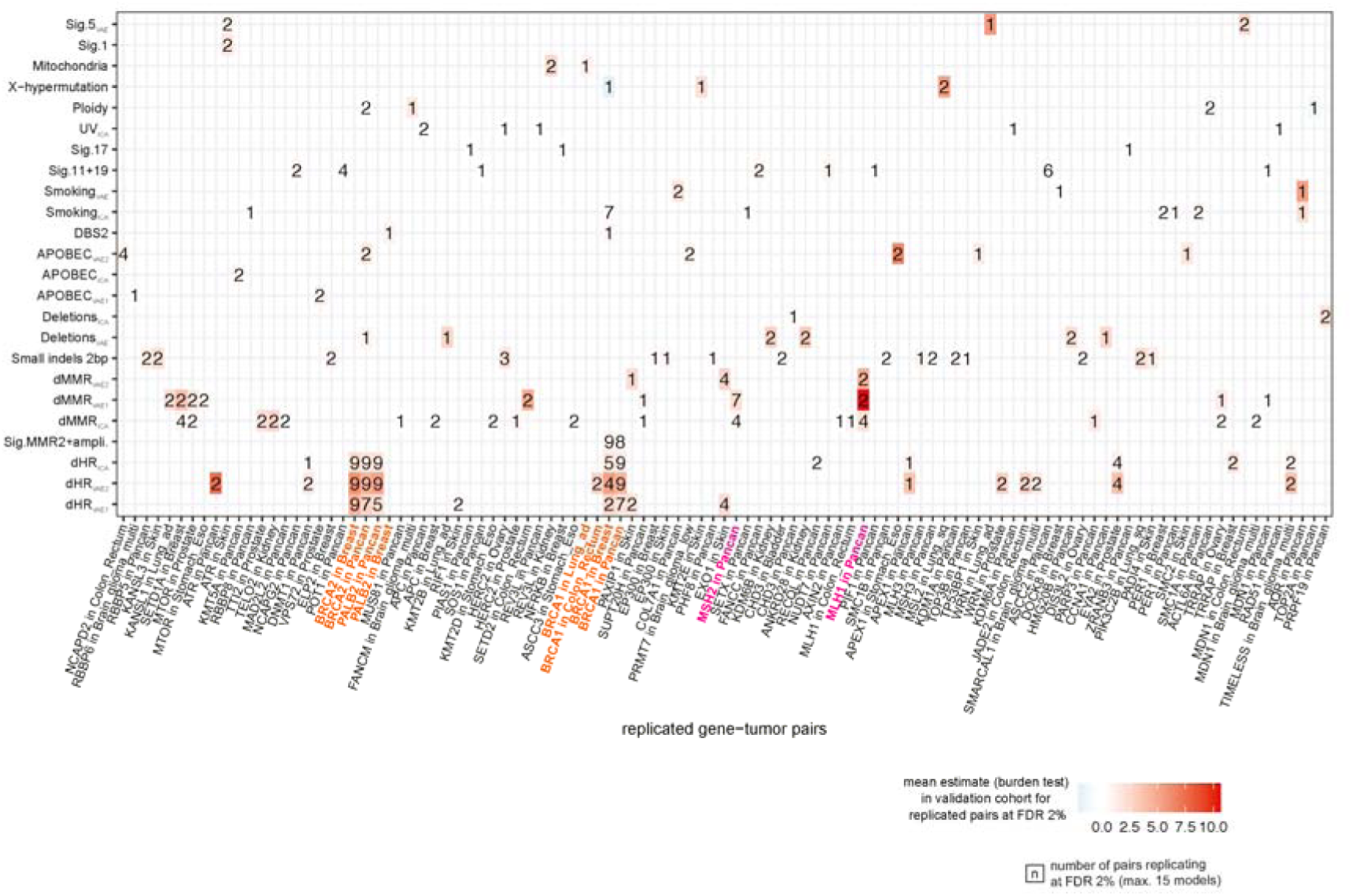
Overview of replicated hits at a FDR of 2%. Showing gene-cancer type pairs (x-axis), the corresponding somatic component (y-axis), and the number of times they replicated at a FDR of 2 % (maximum of 15 models for each gene-cancer type-somatic component tuple). Color code represents the mean estimate of the regression coefficient from burden test for all replicated hits at a FDR of 2 % for the respective gene-cancer type-somatic component tuple. Previously associated dHR genes in orange and dMMR genes in pink. Genes on the x-axis were ordered based on hierarchical clustering results using DepMap CRISPR-derived genetic fitness (Chronos) scores (see Supplementary).

**Extended Data Fig. 5.**
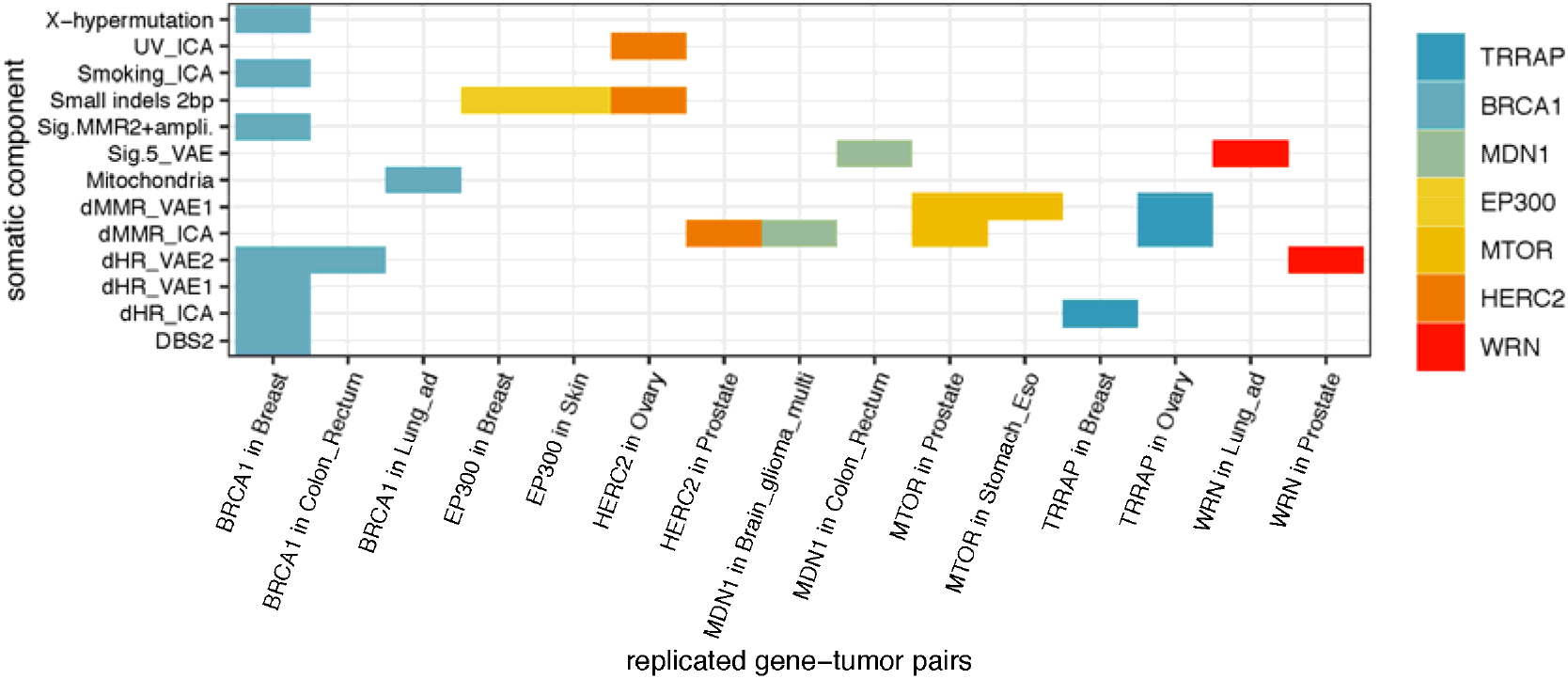
Seven genes associated with a somatic component in ≥ 1 cancer type. Showing gene-cancer type pairs (x-axis) and the corresponding somatic component they associated with at a FDR of 2 % (y-axis). Color code for each gene. Results from pancan analysis excluded.

**Extended Data Fig. 6.**
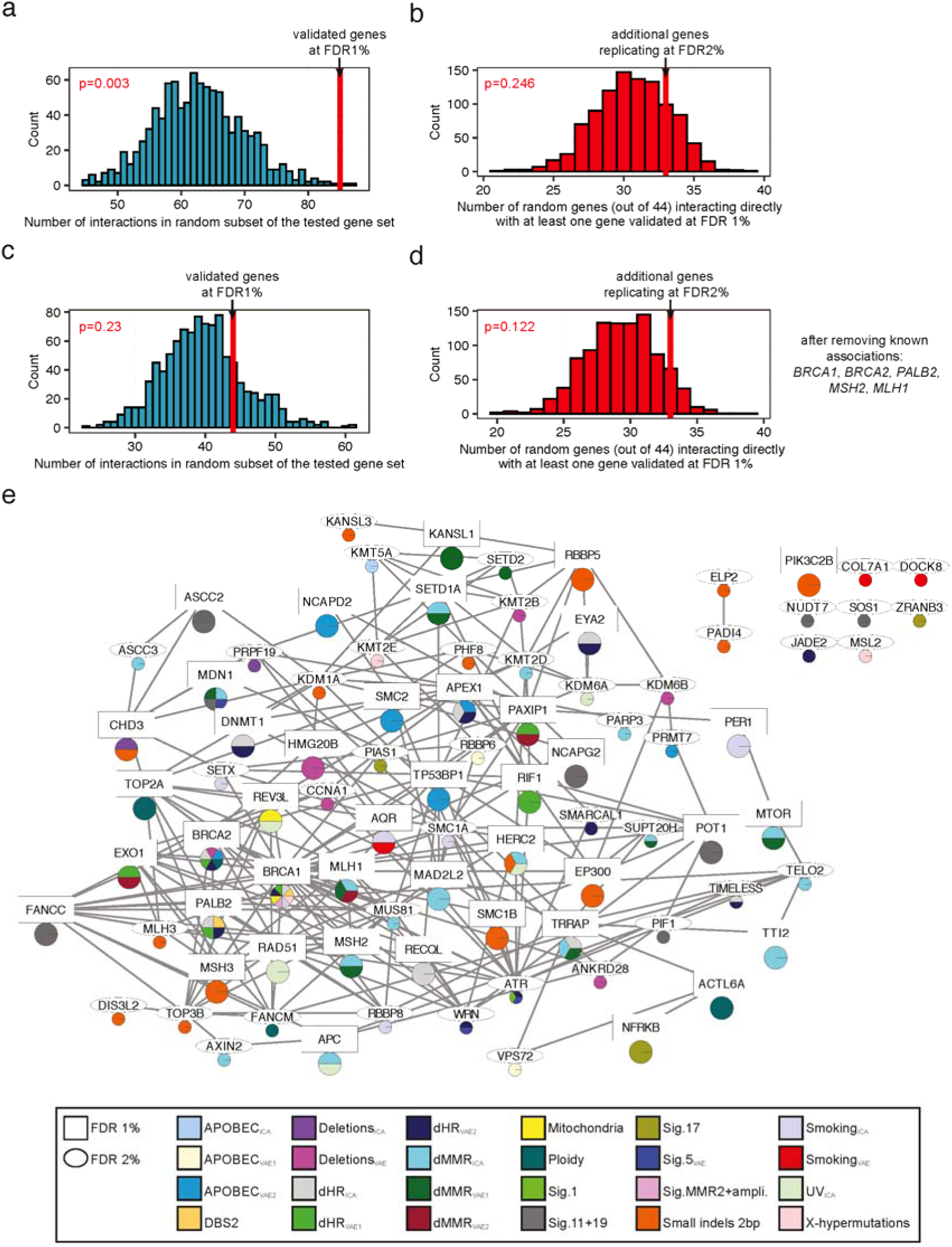
Network analysis (HumanNet) supports the role of rare germline variation in somatic mutational processes. All panels in this figure were generated using the functional gene network from HumanNet. **a**, Number of physical interactions in a random subset of the tested gene set (controlled for interaction node degree) (x-axis). Red line shows the number of interactions within genes which replicated at a FDR of 1 %. **b**, Number of randomly selected genes from the tested gene set interacting with at least one gene, which replicated at a FDR of 1 % (x-axis), (controlled for interaction node degree). Randomization performed 1,000 times. Red line shows the number of genes, out of the ones which additionally replicated at a FDR of 2 %, interacting with at least one gene replicating at a FDR of 1 %. **c**, Same as in a after excluding known genes from the analysis (*BRCA1*, *BRCA2*, *PALB2*, *MSH2*, and *MLH1*). **d**, Same as in b after excluding known genes from the analysis (*BRCA1*, *BRCA2*, *PALB2*, *MSH2*, and *MLH1*). **e**, Visualisation of interactions between proteins for genes replicating at a FDR of 1 % (square) and genes replicating at a FDR of 2 % (ellipse). Color code in pie chart showing the somatic components the corresponding gene associated with. Line width corresponding to interaction score.

## Methods

### Study Design

In this study, the effects of rare damaging germline variants (RDGVs) on different somatic mutational components from cancer genomes were comprehensively analyzed. We utilized genomic sequencing data from three large-scale projects: the Cancer Genome Atlas Program (TCGA)^8^, the Pan-Cancer Analysis of Whole Genomes (PCAWG)^9^, and the Hartwig Medical Foundation (Hartwig)^10^. Associations between RDGVs and somatic features were initially detected in the discovery cohort and hits reaching significance were re-tested in the validation cohort. TCGA WES samples were used as the discovery cohort due to the bigger sample size and WGS samples from PCAWG and Hartwig were aggregated and utilized as the validation cohort.

### Extraction of Somatic Mutational Features and Somatic Components

#### Data Sources in the Discovery Cohort

For the somatic features which were based on SNVs, DNVs, and indels, the somatic calls from the MC3 Project^109^ were used. For the somatic features based on CNVs, TCGA exome data was downloaded from the GDC Data Portal^110^ and processed as described in ref^111^. Copy numbers were identified with the tool FACETS^112^. The tool used as input data the BAM file of the tumor sample, the BAM file of the sample-matched normal sample, and a vcf file of common human SNPs. Furthermore, 93 individuals, which were reported to be positive for human papillomaviruses in head and neck cancer samples^113^, were excluded from the analysis. In total, this yielded somatic calls from 10,033 individuals.

#### Data Sources in the Validation Cohort

Mutation calls for PCAWG were obtained from the ICGC data portal. Somatic mutation calls and copy number calls were obtained from the DKFZ/EMBL variant call pipeline. All samples were downloaded except for ESAD-UK, MELA-AU and all project id’s ending with ‘-US’ in order to prevent an overlap with the discovery cohort. In total, samples from 1,662 donors were downloaded. In short, single nucleotide variants were called via samtools^114^ and bcftools 0.1.19^115^, and indels were called via Platypus 0.7.4^116^. Copy number alterations were estimated with ACEseq v1.0.189^117^ (Supplementary information in PCAWG flagship paper^9^). Data access to the estimated somatic nucleotide variants and copy number variants from Hartwig were acquired as well (https://www.hartwigmedicalfoundation.nl/en/), making up 3,613 samples in total. In Hartwig nucleotide variants were called with Strelka^118^ 1.0.14 and copy number alteration with the Purple tool^10^. BAM files for the melanoma dataset MELA-AU (dataset ID: EGAD00001003388; 183 individuals) and the esophagus dataset ESAD-UK (dataset ID: EGAD00001003580; 303 individuals) were downloaded from the European Genome-Phenome Archive (EGA). Somatic mutations were called via Strelka^119^ 2.9.10 and copy number alterations were extracted as described above with the tool FACETS^112^.

#### Further processing of somatic calls

For all datasets, regions which are known to be difficult to be aligned were excluded as well as regions which have been blacklisted by the UCSC Genome Browser^120^. As described previously^22, 24^ blacklisted regions by Duke and DAC were removed and the CRG75 alignability track was applied to only keep regions, where 75-mers in the genome can be uniquely aligned in the human reference genome hg19.

#### Single Nucleotide Variants - Total Mutation Counts

Based on the number of SNVs in the nuclear genome, 8 different somatic mutational somatic features were estimated: the total number of SNVs, the number of C>A substitutions, the number of C>G substitutions, the number C>T substitutions in regions where the 3’ flanking site was not a G (non CpGs), the number of C>T substitutions in regions where the 3’ flanking site was a G (CpGs), the number of T>A substitutions, the number of T>C substitutions and the number of T>G substitutions. The number of C>T substitutions was divided into two groups (at CpG sites vs. non-CpGs sites) due to the effect of CpG sites on mutation rates (due to DNA methylation)^121^. A pseudocount of 1 was added to each somatic mutational feature and all features were log transformed to the base 2.

#### Single Nucleotide Variants in Mitochondrial DNA - Total Mutation Counts

As other studies have pointed out, WES data can be used to extract mutations occurring in the mitochondrial DNA, due to the large amount of off-target reads^32, 122^. The coverage file of each sample was used to estimate to which extent the mitochondrial genome in each sample was sequenced. Only samples in which at least 50 % of the mitochondrial genome were covered by at least 4 reads were kept for further analysis. Furthermore, following a previous study^32^, only variants were kept which had an allele frequency of at least 3 % to remove potential false-positive calls. For the cancer cohorts Hartwig, ESAD-UK and MELA-AU, which were all based on WGS data, somatic variants in the mtDNA with a frequency of less than 3 % were filtered out as well. After filtering, the total number of SNVs in the mtDNA in each sample was calculated. For PCAWG, mutation calls on the mitochondrial genome were downloaded from the respective study (https://ibl.mdanderson.org/tcma/mutation.html)32,33. At last, a pseudocount of 1 was added to each individual and the feature was log transformed to the base 2.

#### Single Nucleotide Variants - NMF-derived Organ-specific Signatures

First of all, the python tool SigProfilerMatrixGenerator^123^ was used to generate for each dataset a matrix counting all mutations in the 96 possible trinucleotide contexts by considering the adjacent 5’ and 3’ base of the somatic variant (16 trinucleotides for each SNV). Next, the organ-specific signatures, which were derived in the work of Degasperi *et al.*^52^, were fit to each sample. For this step, organ-specific signature exposures were estimated by selecting for each sample the respective organ-specific signature set based on the tissue it was derived from. In cases in which no organ-specific signature set was existing due to its low sample size (e.g. mesothelioma, thymoma, penile, and vulva), the reference mutational signature set was used. In short, this aims to only fit signatures to a sample which were also identified in the according tissue. The tool uses a bootstrap-based method to only assign signatures to a sample when they reach a specific threshold (p < 0.05), otherwise they are set to 0. The goal of this approach is to decrease the probability of overfitting and miss-assignment of signatures^52^. In the discovery cohort the median fraction of unassigned mutations was 47% and in the validation cohort 15%, which is likely due to the low number of somatic mutations in the discovery cohort. To have a common set of signatures, all signature exposures were then converted to the reference signature set via the conversion matrix provided in ref^52^. For further analysis we only kept 17 signatures, which had in the discovery and in the validation cohort an activity of > 5 % in at least one matching cancer type or in the pancan analysis: Ref.Sig.1, Ref.Sig.2, Ref.Sig.3, Ref.Sig.4, Ref.Sig.5, Ref.Sig.7, Ref.Sig.8, Ref.Sig.11, Ref.Sig.13, Ref.Sig.17, Ref.Sig.18, Ref.Sig.19, Ref.Sig.22, Ref.Sig.30, Ref.Sig.33, Ref.Sig.MMR1 and Ref.Sig.MMR2. A pseudocount of 1 was added and each estimated signature count was log transformed to the base 2.

#### Single Nucleotide Variants - Transcriptive Strand Bias

To estimate the transcriptive strand bias, the number of mutations occurring on the untranscribed strand and on the transcribed strand were calculated. This was performed by the python tool SigProfilerMatrixGenerator^123^. Based on the six possible base substitutions, six different somatic features were generated (C>A, C>T, C>G, T>A, T>C, T>G). For each one, the number of base substitutions occurring on the untranscribed strand were divided by the number of mutations occurring on the transcribed strand. A pseudocount of 1 was added to the numerator and denominator before division and the resulting quotient was log transformed to the base 2.

#### Single Nucleotide Variants - Replicative Strand Bias

To estimate the replicative strand bias, replication timing data from lymphoblastoid cell lines was downloaded (http://mccarrolllab.org/resources/)124. The fork polarity, which is a derivative of the replication timing estimate, was estimated as described by Seplyarskiy *et al.*^125^. In brief, the slope/derivative at each coordinate of the replication timing landscape was calculated by considering the region approximately ± 5 kb of the coordinate. The fork polarity value reflects whether the reference strand is more likely to be replicated as the leading strand (fork polarity > 0) or as the lagging strand (fork polarity < 0). Next, the genome was divided into equal sized bins of the length of 10 kb and the average fork polarity in each bin was calculated. Further, the whole genome was split into 10 equal sized bins based on the fork polarity estimate. To calculate the replicative strand bias, we only considered the two lowest bins (reference strand more frequently replicated as the lagging strand) and the two highest bins (reference strand more frequently replicated as the leading strand). From the perspective of the reference strand, we divided the total number of T>C, T>G, G>A, and C>A mutations occurring on the leading strand by the total number of T>C, T>G, G>A, and C>A mutations occurring on the lagging strand. This would mean for instance that a A>G mutation occurring on the leading strand was counted as a mutation occurring on the lagging strand (since T>C on the other strand). We focused on these four mutation types since replicative strand biases have been previously reported for these in connection with a deficiency in DNA mismatch repair^39^. This feature was only calculated in samples, in which at least 20 of the 4 single substitutions types were counted within the covered region. The estimated values were log transformed to the base 2.

#### Single Nucleotide Variants - X-Chromosomal Hypermutation

For generating a somatic mutational feature for X-Chromosomal hypermutation^31^, first of all the total number of single nucleotide variants per megabases (MB) on each chromosome was counted. Next, the number of mutations per MB occurring on the X chromosome was divided by the average number of mutations per MB occurring on the autosomes. A pseudocount of 0.1 was added to the numerator and denominator before division and the resulting quotient was log transformed to the base 2.

#### Single Nucleotide Variants - CTCF/Cohesin Binding Sites

CTCF/cohesin binding sites are often mutated in cancer^29, 30^. To capture this somatic mutational feature, we counted the number of single nucleotide variants occurring in CTCF/cohesin binding site and divided them by the number of mutations occurring in the flanking site (± 500 bp) of the binding site. CTCF/cohesin binding sites were obtained from Roadmap^126^ and averaged over 8 cell types. Genomic regions, that were bound by CTCF in at least one cell type and by cohesin in at least two cell types were set as CTCF/cohesin binding sites. All sites ± 500 bp of the sites that were bound by CTCF in at least one cell type were set as the flanking site. Length of covered genomic regions can be found in Supplementary Table 1. This somatic feature was only estimated in samples which had at least 10 SNVs counted in total within the CTCF/cohesin binding and/or flanking site. At last, we were able to calculate the CTCF somatic feature for 38 % of the samples in the discovery cohort and 98 % of the samples in the validation cohort. The ratio was log transformed to the base 2.

#### Extraction of Genomic Region Densities of Expression, Histone Mark H3K36me3, Replication Timing and DNase I Hypersensitive Sites

Features measuring mutation rate variation with regards to expression, histone mark H3K36me3, replication timing, and DNase I hypersensitive sites were calculated using negative binomial regression to reduce the correlation of these features with each other and to control for mutation substitution types. For this purpose, regional data from a previously published study was used^24^. In brief, levels of histone mark H3K36me3 (averaged over 8 cell types) and DNase I hypersensitive sites were downloaded from Roadmap Epigenomics^126^. Genomic regions with no signal for the corresponding feature were set as ‘bin 0’ and the remaining genomic regions were split into 5 equal sized bins with increasing signal. In this way, genomic regions with the highest amount of histone mark H3K36me3 were put into bin 5, regions with the lowest amount into bin 1 and regions with no signal into bin 0. Replication timing information was derived from the ENCODE project using the average over 8 cell lines. Genomic regions were split into 6 equal sized bins, where bin 1 corresponded to the latest replicating region and bin 6 to the earliest replicating region. Expression levels were based on RNA-seq data, which was obtained from Roadmap^126^ and averaged over 8 cell types as well. Bin 0 represented regions with no expression (RPKM = 0) and the remaining 5 bins were split equally by increasing expression levels. All these genomic masks from ref^24^ were further processed by applying the CRG75 alignability track. For WES data specifically, the masks were intersected with the coverage mask from the MC3 project^109^, since the somatic WES mutation calls were derived from there. Furthermore, the 4 masks (expression, histone mark H3K36me3, replication timing, and DNase I hypersensitive sites) were intersected with each other for the subsequent regression. Several bins extracted from the whole exome mask covered only a small region in the genome (< 5 MB), which was expected since the exonic regions in the genome are known to be enriched for early replicating regions and histone mark H3K36me3. Since we observed that the regression often failed when bin sizes were too small, some bins were merged: replicating timing bins 1 and 2, histone mark H3K36me3 bins 1 and 2, expression bins 0 and 1, and DNaseI hypersensitive site bins 1 and 2. This step was not performed for the whole-genome masks since the covered regions for each bin were big enough. Length of covered genomic regions can be found in Supplementary Table 1.

#### Single Nucleotide Variants - Mutation Enrichment Calculations with regards to Expression, Histone Mark H3K36me3, Replication Timing and DNase I Hypersensitive Sites

The individual features corresponding to the enrichment of mutations in a particular genomic region were calculated via negative binomial regression using the function *glm.nb* from the R package MASS (version 7.3_53.1) in R 3.5.0 The regression was performed for the different features in each tumor sample as follows:

I. mutation count ∼ replication timing + mutation type + offset
II. mutation count ∼ replication timing + DNase + mutation type + offset
III. mutation count ∼ replication timing + expression + mutation type + offset
IV. mutation count ∼ replication timing + H3K36me3 + mutation type + offset

In the discovery cohort (WES only) the mutation type variable had 7 possible encodings (C>A, C>T at CpG sites, C>T at non-CpG sites, C>G, T>A, T>C and T>G), and in the validation cohort (WGS only) the mutation type variable encompassed all 96 possible substitutions within the trinucleotide context (e.g. C>A mutation within ACA context). The offset represents the nucleotide-at-risk and is the natural log of the number of nucleotides covering the respective region. As described previously^24^, the coefficients obtained from the regression for the different genomic regions represent the log enrichment of mutations in each bin in comparison to a reference bin. For replication timing, the latest replicating bin was set as the reference, for expression the lowest expressing bin was set as the reference and for histone mark H3K36me3 and DNase I hypersensitive sites the bins with no signal were set as the reference. This would mean that for instance the coefficient obtained from regression IV. for bin 5 from the histone mark H3K36me3 variable describes the log enrichment of mutations in regions with a high signal of this histone mark in comparison to regions with no histone mark signal, while controlling for replication timing and the mutational context. In this way, we aimed to control for the correlation of expression levels, histone mark H3K36me3 and DNase I hypersensitive sites with replication timing and the mutational context. Especially, for WES data this approach was limited by the reduced covered genomic region and the decreased number of mutations in comparison to WGS data. The regression was only performed in samples, which had at least 30 SNVs counted. The coefficient obtained in regression I. for the earliest replicating bin was extracted for the replication timing feature, the coefficient obtained in regression II. for the bin with the highest amount of signal in DNase I hypersensitive sites was extracted for the DNase I hypersensitive site (DNase) feature, the coefficient obtained in regression III. for the bin with the highest expressing regions was extracted for the expression (Expression) feature, and the coefficient obtained in regression IV. for the bin with highest amount of signal in histone mark H3K36me3 was extracted for the H3K36me3 (H3K36me3) feature. High errors in the regression coefficients (standard error > 100) indicated that the regression failed to converge for the corresponding coefficient and thus, were removed. In the discovery cohort, 7,650 replication timing coefficients, 7,684 H3K36me3 coefficients, 7,471 DNase coefficients and, 7,664 Expression coefficients were extracted in total. In the validation cohort, 5,759 RT coefficients, 5,749 H3K36me3 coefficients, 5,752 DNase coefficients and, 5,759 Expression coefficients were extracted in total.

#### Double Nucleotide Variants - NMF-derived Signatures and Fitting

Double nucleotide variants were extracted with the python tool SigProfilerMatrixGenerator^123^. The tool counted the occurrence of 78 double nucleotide variants (AC, AT, CC, CG, CT, GC, TA, TC, TG, or TT to NN). The matrix was used as an input to extract Double Base Substitution (DBS) signatures using the python tool SigProfilerExtractor^19^. In brief, the tool uses non-negative matrix factorization (NMF) to extract mutation signatures. Since the exact number of mutation signatures is not known, the tool extracted 1 to 25 signatures. For each signature extraction 100 iterations were performed adding poisson noise to the samples during each iteration. For the discovery cohort the optimal solution was 3 signatures and for the validation cohort 11. Next, the tool fitted the established DBS signatures from COSMIC^13^ v3.2 to the extracted de-novo signatures. Then, signature exposures were estimated by fitting the extracted COMISC signatures to each sample. In the discovery cohort the COSMIC^13^ DBS signatures DBS1, DBS2, DBS4, DBS9 and DBS10 were extracted and in the validation cohort the DBS signatures DBS1, DBS2, DBS4, DBS5, DBS6, DBS7 and DBS9 were extracted. The 4 DBS signatures which were found in both cohorts were kept for association testing: DBS1, DBS2, DBS4 and DBS9. Next, a pseudocount of 1 was added to each estimated signature exposure and each estimated exposure was log transformed to the base 2.

#### Insertions and Deletions - Total Mutation Counts

Different insertion and deletion somatic mutational features were generated. First of all, the total number of indels occurring in each sample was counted. Next, the number of indels in microsatellite (MS) regions was counted due to its frequent occurrence in samples with dMMR^18, 127^. For this purpose, the number of indels with a length of 1 bp and the number of indels with a length of 2 to 5bp were counted within and outside MS regions. MS locations were identified via the tandem repeat search tool Phobos (https://www.ruhr-uni-bochum.de/ecoevo/cm/cm_phobos.htm). Next, the total number of indels with a length of 6 to 10 bp was counted. Due to the low number of indels of this length, especially in WES data, this feature was not further split into MS vs non-MS regions. Furthermore, since deletions have often been reported to be predictive of dHR^40^, different deletion features were created. The total number of deletions with a length of bigger than or equal to 10 bp was created. Also, the number of deletions at flanking microhomology sites of either 1 bp or more than 1 bp was counted by using the output matrix from the python tool SigProfilerMatrixGenerator^123^. A pseudocount of 1 was added to each feature and each feature was log transformed to the base 2.

#### Insertions and Deletions - NMF-derived Signatures and Fitting

Small insertion and deletion (ID) signatures were extracted in the same way as described for the DBS signatures. For the discovery cohort the optimal solution was 4 signatures and for the validation cohort 10. The COSMIC^13^ ID signatures were fit to the de-novo signatures and in the discovery cohort COSMIC^13^ ID signatures ID2, ID3, ID4, ID7, ID8 and ID15 were extracted and in the validation cohort ID signatures ID1, ID2, ID3, ID4, ID5, ID6, ID8, ID9, ID10, ID12, ID13 and ID14 were extracted. The 4 ID signatures which were found in both cohorts were kept for further association testing: ID2, ID3, ID4 and ID8. Next, a pseudocount of 1 was added to each estimated signature exposure and each estimated exposure was log transformed to the base 2.

#### Copy Number Variants - Total Mutation Counts, Ploidy and Whole Genome Duplications

Copy number based features were generated by splitting amplification and deletion events by different sizes. The number of amplifications with a size of 1 to 10 kb, 10 to 100 kb, 100 to 1000 kb, and bigger than 1000 kb were counted. Similarly, the number of deletions with a size of 1 to 10 kb, 10 to 100 kb, and bigger than 100 kb were counted. Next, a feature was generated based on the estimated ploidy of the tumor sample from the corresponding copy number detection tool. The number of whole genome duplication events were calculated by dividing the ploidy by 2 via integer division. A pseudocount of 1 was added to the amplification and deletion based features, a pseudocount of 0.1 was added to the WGD feature and no pseudocount was added to the ploidy feature since ploidy can never be 0. At last, each feature was log transformed to the base 2.

#### Generation of the Input Matrix for ICA and VAE

For the ICA and VAE all somatic features described above were used except for the following 9 somatic features: total number of SNVs, total number of indels and total number of the 7 different single mutation substitutions types (Supplementary Fig. 1 and 2). These were excluded since they were already represented by the different NMF-derived signatures. Further, all samples were removed in which more than 20% of the features were not estimated due to low mutation counts. Thus, 9,235/9,425 samples were left in the discovery cohort and 5,597/5,613 samples were left in the validation cohort. Next, missing values were replaced by the median value of the respective columns and each feature was centered and standardized to a mean of 0 and standard deviation of 1. This step was performed for the somatic features, which were extracted from three different cohorts (TCGA, Hartwig, PCAWG) separately to control for potential biases. Then, the three matrices were merged (samples as rows, features as columns).

#### Independent Component Analysis

The ICA was run on the 56 somatic features using the input matrix as described above. Similarly, as for the NMF, the number of ICs needs to be set before running the ICA. The methodology to extract the optimal number of components was adapted from the methodology applied previously^24^ to extract the optimal number of NMF derived components. For the extraction of ICs the R package fastICA (version 1.2.1) in R 3.5.0 was used. The ICA was run by varying the number of extracted components from 2 to 30. For each component extraction the ICA was run 200 times and the seed for the random number generator was changed before every iteration. In each iteration the ICA decomposes the input matrix into a loadings matrix (corresponding to the components and their attributed weight from each somatic feature) and a scoring matrix (also called source matrix; samples projected to new component axes). After 200 iterations, the 200 loadings matrices were combined and clustered using k-medoids clustering with varying k from 2 to 50. Clustering was performed with the function *pam* from the R package cluster (version 2.0.6). For each clustering the average of the mean silhouette indexes of each cluster were saved as well as the lowest and second lowest mean silhouette index of a cluster extraction. Later, extracted summary silhouette indexes for different extracted IC numbers were plotted against the different number of extracted clusters (Supplementary Fig. 3). The optimal number of components was decided visually based on the broken-stick approach (Supplementary Fig. 4). For a given extracted number of ICs, the optimal number of clusters was always times 2 since during each iteration, signs flipped randomly and thus, each component always had a ‘mirrored’ counterpart with opposite signs (Supplementary Fig. 5). In the end, always one component of the mirrored pair was kept. For the ICA, 15 unique ICs (using 30 clusters) were extracted. Correlations were estimated by calculating the Pearson correlation of each input somatic feature with each estimated score of each IC. Contributions were calculated by squaring the estimated loading matrix and dividing the squared loading by the sum of the loadings for the respective IC. Thus, the sum of the contributions (56 somatic input features for each IC) for each IC equals 1 (100 %) (Supplementary Fig. 6).

#### Extraction of Components via a Variational Autoencoder

The architecture of the VAE was adapted from studies from Way *et al.*^105, 128^ (https://github.com/greenelab/tybalt/blob/master/tybalt_vae.ipynb), where they applied a VAE to compress gene expression data to extract biologically relevant representations. The script was modified for our purposes. In short, it is a simple ladder-VAE architecture consisting of one encoding and one decoding layer to generate a generalizable representation of the input and to use this representation to reconstruct the input. Batch normalization was performed in the encoding layer before applying the activation function *ReLu*. In the encoding layer the VAE learned a distribution of means and standard deviations to generate the latent space. This latent representation was then decoded in the decoding layer by applying the *tanh* function as the final activation function. Weights were initialized via the Glorot uniform initializer. We also tested adding an additional layer between the input and the encoding layer and between the latent space and the decoding layer. The extra layer always had 2 times more dimensions than the latent space and involved a batch normalization step before applying the *ReLu* activation function. The reconstruction loss was the sum of the mean squared error and the KL-divergence loss. To encourage learning, the ladder-VAE makes use of a so called *warm* start, meaning that it starts training without the KL divergence loss and linearly increases the contribution of the KL divergence loss after each cycle via the parameter *beta* (mean squared error+*beta*\*KL divergence loss). The linear increase of the contribution of the KL divergence loss was controlled via the parameter *kappa*.

In contrast to a previous VAE architecture^105, 128^, we applied the *tanh* function in the final decoding layer and used the mean squared error as part of the reconstruction loss since our input was not binary. To reconstruct the input via the *tanh* function, all the somatic features were transformed to a range of -1 to 1 prior to running the VAE. The data was split into 90 % training data and 10 % validation data and stratified by gender and cancer type. Performance was evaluated by checking the mean correlation of the reconstructed validation set with the validation input set and by calculating the correlation with selected ICs, which were shown to represent biologically relevant components. For this purpose, we calculated the maximum correlation of the components from the latent space of the VAE to the ICs dMMR_ICA_, dHR_ICA_, Smoking_ICA_ and UV_ICA_ and then calculated the average. To find the optimal hyperparameters we performed a grid search testing over 4,300 hyperparameter combinations (Supplementary Fig. 8). After finding the optimal hyperparameters, the VAE was run for different latent space dimensionalities 5 times with different random initializations (Supplementary Fig. 9). In the end, the results from using a latent space with 14 dimensions was extracted for further downstream analysis using the architecture with no extra layer between input and encoder and with no extra layer between decoder and output (Supplementary Fig. 10 and 12).

The VAE was run in a singularity container. A docker file was generated based on the docker image *tensorflow/tensorflow:1.15.5-gpu-py3-jupyter* and the python modules *scipy*, *scikit-learn*, and *seaborn* were added. The resulting docker image was then uploaded into Docker Hub and run in a singularity container. Python version 3.6.9, keras version 2.2.4 and tensorflow version were used in this environment.

#### Estimation of Tissue Enrichments of Components

Tissue enrichments of individual components (Supplementary Fig. 7 and 13) were calculated as follows. For each component it was tested whether the component scores from one cancer type were significantly different to the scores of the remaining cancer types via a two-sided Welch’s t-test. In addition, Cohen’s d statistic was calculated between the two groups. This test was performed for each cancer type and separately for the two cohorts (TCGA and PCAWG + Hartwig). Cancer types were then grouped into their corresponding tissue of origin and the average Cohen’s d statistic was calculated.

### Identification of Rare Damaging Germline Variants

#### Extraction of Rare Germline Variants in the Discovery Cohort

TCGA bam files were downloaded as described here^111^. Strelka^119^ 2.9.7 was run on TCGA WES normal and tumor samples to extract germline variants. Germline variants called in the tumor samples (will be a mix of germline and somatic mutations) were used later in a downstream step to only keep germline variants which were identified in the normal and tumor tissue. In this way, we aimed to remove potential false-positive germline calls in the normal sample and to remove variants which were selected out in the tumor and thus, irrelevant for our association analysis. Germline variants which were called in the normal sample with the filter PASS were kept as well as variants which were called with the filter LowGQX but had a GQX of at least 10. Variants which were found inside gnomAD^129^ with the filter PASS and had a GQX of at least 10 were kept as well as variants which were not found inside gnomAD^129^, but had a GQX of at least 20. Next, variants were annotated via ANNOVAR^130^ (version 2019-10-24), CADD v.1.6 scoring was added, and only exonic and splicing variants were kept. Furthermore, only variants which had allele frequency of less than 0.1% in gnomAD^129^ (overall and in each sub-population) were kept as well as variants which were not found inside gnomAD^129^. Variants with a frequency equal to or higher than 1 % within the cohort were removed. Additionally, rare germline variants were only kept when they were also found in the matching tumor sample.

#### Generation of a Coverage File for TCGA

We used the same methodology as described in previous work^53^ to only extract genomic regions with sufficient coverage to be sure that regions in which no damaging germline variant was called was not due to lacking coverage. In brief, within each sequencing center (BI, WU, and BCM) 100 coverage files were randomly selected. Genomic regions which were covered by at least 8 reads in 90 % of the samples within each sequencing center were kept. Next, the coverage masks of the 3 sequencing centers were intersected, making up in total a genomic mask of 60 MB in length. Only genomic regions within these sites were kept for further analysis.

#### Extraction of Germline Variants in the Validation Cohort

Germline variants from PCAWG, Hartwig, ESAD-UK and MELA-AU were all processed in the same way if not indicated otherwise. Each cohort was processed at the beginning separately due to the different formats. The files were combined in the end. While germline calls from PCAWG and Hartwig were obtained as described above, germline variants in ESAD-UK and MELA-AU were called via Strelka^119^ 2.9.10 (same approach as in TCGA), and derived from the same datasets from which the somatic calls were obtained as well. Thus, for ESAD-UK and MELA-AU the same approach as for TCGA was applied. For PCAWG and Hartwig, germline calls with the filter PASS by the respective germline detection tool were kept. Next, variants which were found inside gnomAD^129^ and had the filter PASS were kept as well as variants which were not found inside gnomAD^129^ (rare singletons). Variants were annotated via ANNOVAR^130^ (2019-10-24). All variants which were found inside gnomAD^129^ were required to have an allele frequency of less than 0.1 % (overall and in each subpopulation). Exonic and splicing variants were extracted. Furthermore, variants outside the CRG75 alignability mask were filtered out and variants with a frequency equal to or higher than 1 % within each cohort were discarded as well. The rare germline calls from the different cohorts were combined. Further, in all cases in which germline calls were also available for the matching tumor sample, variants were filtered out if they were not found in the matching tumor sample. Germline calls for matching tumor samples were available for PCAWG, ∼80% of Hartwig, and not available for ESAD-UK and MELA-AU.

#### Definition of Rare Damaging Germline Variants

In this study 5 definitions of Rare Damaging Germline Variants (RDGVs) were applied in addition to requiring an allele frequency of < 0.1 % (described above):

I. RDGV= protein truncating variants (PTVs)
II. RDGV= PTVs + Missense variants with a CADD^58^ ≥ 25
III. RDGV= PTVs + Missense variants with a CADD^58^ ≥ 15
IV. RDGV= Missense variants with a ’missense tolerance ratio’^61^≤ 25th percentile
V. RDGV= Missense variants with a ’constrained coding region’^60^ value ≥ 90th percentile

For case I. only PTVs were considered. PTVs comprised in this study frameshift deletions, frameshift insertions, stoploss variants, stopgain variants, startloss variants and splicing variants. Splicing variants comprise the canonical splice variants annotated by ANNOVAR^130^ (version 2019-10-24) and variants with a predicted donor loss or acceptor loss higher than 0.8 by SpliceAI^131^. Pre-computed SpliceAI score files were downloaded from Illumina Basespace and annotations were added to each variant (hg38 for the discovery cohort and hg19 for the validation cohort). For cases II. and III. potentially damaging missense SNVs were added on top of PTVs. Deleteriousness was assigned via the phred-scaled CADD^58^ scores. For case IV. we only considered missense SNVs with a missense tolerance ratio (MTR)^61^ lower or equal to the 25th percentile and for case V. we only considered missense SNVs with a constrained coding region (CCR)^60^ value equal or bigger than the 90th percentile. On top of these variant filtering steps, two additional filtering steps were applied to all five RDGV sets in order to discard potential false-positive RDGVs: the proportion expressed across transcripts (PEXT) metric^132^ and the terminal truncating exon rule^129^.

#### Filtering out Non-Expressed Variants via the PEXT Metric

The PEXT^132^ score was introduced in one of the gnomAD articles and in brief, estimates to which extent a variant is expressed in a tissue based on isoform transcription levels from RNA-seq data. PEXT scores were estimated using over 11,000 tissue samples from GTEx. Thus, PEXT scores were downloaded and added to the variant annotations. Since hg38 was used for the germline calls in the discovery cohort, PEXT annotations were first converted from hg19 to hg38 via the liftover tool from UCSC^120^ (version021620). This step was not necessary for the validation cohort. Variants were only kept when they had a PEXT value higher than 0.1 in the matching GTEx tissue. Matching a cancer type with the most appropriate GTEx tissue was mostly guided by a previous study^133^. For cases in which no matching GTEx tissue was available for a cancer type, the mean PEXT value was used. This filter was applied to all variants not affecting splicing since many splicing variants are close to exon borders and thus, don’t have a PEXT score.

#### Exclusion of Terminal Truncating Exon Variants (with exceptions)

Terminal truncating variants might not have a deleterious loss-of-function effect since they can escape nonsense-mediated decay and still be functional. For these reasons, they have been also removed in the loss-of-function transcript effect estimator (LOFTEE) of gnomAD^129^. Hence, variants occurring in the terminal exon were removed. This filter was not applied in cases in which the variant was predicted to have a deleterious effect by CADD^58^ ≥15 or in cases in which the variant was predicted to have a splicing effect. In this way, we aimed to reduce the risk of losing potentially harmful variants, which as described in the gnomAD flagship paper^129^, can be the case when the C-terminal domain of a protein exerts a crucial function. To identify variants occurring in the last exon, gene coordinates were downloaded from UCSC^120^ using the NCBI RefSeq track^134^. Exon coordinates of the last exon of the longest transcript were kept. These coordinates were then intersected with the variant coordinates to detect variants occurring in terminal exons.

### Detecting and Assigning putative Loss of Heterozygosity (LOH)

#### Detecting and Assigning putative LOH in the Discovery Cohort TCGA

To detect LOH, we considered the copy number calls from FACETS^112^. FACETS calls were available for 9,814 samples. We extracted all ’LOH’ and ’DUP-LOH’ calls and assigned them to genes by intersecting the extracted coordinates with gene coordinates from NCBI Refseq^134^ hg38. We assigned LOH to a gene in samples in which LOH was called via FACETS + the variant allele frequency of the RDGV was not higher in the normal sample than in the tumor sample and the variant allele frequency of the RDGV was not higher than in the tumor and sample-matched normal sample. In this way, we aimed to only consider LOH events, when the putative RDGV of interest got enriched in the tumor via LOH since this was the tested hypothesis for the recessive and additive model. For 441 samples for which we did not have any FACETS calls, we assigned LOH to a gene in a sample when the difference in the variant allele frequency of the putative RDGV between tumor and normal sample was higher than 0.25 and when the variant allele frequency of the putative RDGV was higher than 0.8 in the tumor and sample-matched normal sample.

#### Detecting and Assigning putative LOH in the Validation Cohort

For PCAWG (excluding ESAD-UK and MELA-AU), CNV calls from ACEseq^117^ v1.0.189 were further processed. All passed calls with the assignments ’LOH’, ’LOHgain’ or ’cnLOH’ were extracted and genes were assigned to the LOH events as before (using NCBI Refseq^134^ hg37). We excluded LOH calls when the corresponding RDGV in the respective gene had a lower allele frequency in the tumor than in the sample-matched normal sample and the allele frequency was not higher than 0.8 in both tissues.

For ESAD-UK and MELA-AU, CNV calls were available via FACETS^112^ and LOH was called as described for TCGA. In contrast to the steps performed for TCGA, germline calls from the tumor tissue were not available for ESAD-UK and MELA-AU. Thus, LOH calls were not further filtered.

For Hartwig, CNV calls were provided via the tool Purple^10^. LOH was assigned to locations in which the minor allele ploidy was lower than 0.4. LOH calls were excluded in cases in which the allele frequency of the RDGV was lower in the tumor than in the sample-matched normal tissue and the allele frequency of the RDGV was not higher than 0.8 in the normal and tumor tissue. This was only applicable to the samples in which germline calls from the tumor genome were available (678 samples with germline calls from tumor genomes not available).

### Gene-Based Rare Variant Association Testing

#### Extraction of Common Germline Variants and Sample-level Quality Control

Common variants were extracted from the normal samples to apply some sample-level quality control as well as to prepare the data to perform a PCA for extracting population ancestry. The following steps were performed for the discovery cohort (TCGA) and the validation cohort (PCAWG and Hartwig) separately. Germline variants which were called with the filter PASS were kept. Also, in accordance with the extraction of rare germline variants, variants with the filter LowGQX but a GQX ≥ 10 were kept in the respective cohorts (TCGA, ESAD-UK and MELA-AU). Common variants were extracted by only keeping variants which were identified inside gnomAD^129^ with the filter PASS and with an allele frequency > 5 % within the overall population. In TCGA all variants within the generated genomic mask were retained and in the other cohorts all variants within the CRG75 alignability mask were retained. Loci, in which more than 2 alleles existed, were removed. The total number of common variants inside each sample was calculated and within each cohort (TCGA, Hartwig, PCAWG) samples with an altered number of variants 1.5 standard deviations away from the mean were discarded (214 samples in TCGA, 212 samples in Hartwig, 204 samples in PCAWG) (Supplementary Fig. 17a and 18a). Next, common variants for each cohort were uploaded into PLINKv1.90b6.1 and further processed there. Missing genotypes were set as homozygous for the reference allele. Only variants with a MAF > 5 % were retained and samples with a heterozygosity rate ± 3 standard deviations away from the mean were removed (127 samples in TCGA, 54 samples in Hartwig, 39 samples in PCAWG) (Supplementary Fig. 17b and 18b). For the following steps, variants on the sex chromosomes, on the mitochondrial chromosome and within regions with high amount of linkage disequilibrium (https://github.com/meyer-lab-cshl/plinkQC/tree/master/inst/extdata) were removed. Also, variants extensively deviating from the Hardy-Weinberg-equilibrium with p < 10−6 were excluded.

#### Identification of Duplicated or Related Individuals

The dataset was pruned on the discovery cohort (TCGA) and on the merged validation cohort (PCAWG and Hartwig) separately, applying a window size of 50 bp, a step size of 5 and a r^2^ threshold of 0.2. The identity-by-state (IBS) matrix was calculated for all pairs of individuals within each cohort. Within all pairs of individuals with identity-by-descent (IBD) > 0.185 (0.185 would be the expected value for individuals between third- and second-degree relatives) one individual was removed (542 samples in TCGA, and 479 samples in PCAWG and Hartwig) (Supplementary Fig. 17c, 18c, and 18d).

#### Extraction of European Individuals

To extract individuals of European ancestry the pruned dataset was used and a principal component analysis (PCA) was performed. The PCA was run on the discovery cohort and on the merged validation cohort (Supplementary Fig. 19 and 20). The first ten principal components were used for clustering using the R package tclust (version 1.4.2), which trimmed 1 % of the outlying samples as described previously^53^. Individuals were grouped into k = 10 clusters and European groups were selected based on the reported TCGA/PCAWG annotations. In total 7,864 individuals were retained in the discovery cohort and 4,691 individuals were retained in the validation cohort. The PCA was repeated on the pruned dataset for the individuals of European ancestry in the respective cohorts to extract the PCs, which were used as covariates in the association testing (Supplementary Fig. 21 and 22).

#### Gene-Based Rare Variant Burden Testing

As described above 29 somatic mutational components were extracted from the discovery and validation cohort from the tumor genomes. RDGVs were extracted from the sample-matched normal samples. Gene-based rare variant burden testing was only performed on samples which survived the quality control filters (as described above). We limited the analysis to individuals with European ancestry due to the bigger sample size. In addition, only samples were kept, in which at least 10 SNVs were counted. In total 6,799 samples were left in the discovery cohort for testing and 4,683 samples were left in the validation cohort for testing.

#### Gene Set

For testing, RGDVs occurring in 892 different genes were extracted. The gene set covered DNA damage response genes^135^, known cancer predisposition genes^36^, genes involved in chromatin organization (https://pathcards.genecards.org), genes involved in DNA double strand repair (https://pathcards.genecards.org), genes which were reported to regulate MSH2 stability^81^, and human homologs of genes, in which heterozygous mutations were reported to cause genetic instability in *Saccharomyces cerevisiae*^50^.

Effectively, out of the 892 individual genes 746 genes were tested in the most permissive RDGV set (set III.) in pancan. The remaining genes were not tested in the discovery cohort since not enough RDGVs were identified in these genes to test them.

#### Association Testing via SKAT-O

Association testing was performed in each cancer type separately and with all cancer types together (pancan). The effect of a gene on a somatic component was only tested when a RDGV in that gene was identified in at least two individuals. Testing was performed across 12 cancer types as shown in Supplementary Table 5. Accordingly, depending on the cancer type different numbers of genes were tested in total.

Association testing was conducted via the unified testing approach of SKAT-O^54^. In short, SKAT-O combines the tests SKAT^56^ and burden via a weighted mean:

• *Q*_ρ_ =ρ\**Q_B_* + (1-ρ)*Q_S_*.

Here, *Q*_ρ_ is the final statistic from the weighted mean of the burden statistic *Q_B_* and SKAT statistic *Q_S_*. The parameter ρ influences how strongly each test is weighted. SKAT-O testing was performed via the R package SKAT^54^ 2.0.1. For testing, the covariates were firstly regressed against the somatic components with the function *SKAT_Null_Model*. When applicable, age of diagnosis, sex, ancestry (first 6 PCs) and cancer type were used as covariates. Categorical variables were encoded as dummy variables with the R package fastDummies 1.6.3. Missing age information was imputed by taking the median value in the respective cohort. After initializing the null model, SKAT-O was run by using the function *SKAT* and setting the method to *SKATO*. The function ran SKAT-O with 10 different values of ρ (from 0 to 1) and reported the ρ value which led to the lowest p-value.

Three models of inheritance were tested in total and individual variants were encoded as follows:

I. **Dominant**: no RDGV = 0; RDGV = 1
II. **Additive**: no RDGV = 0; RDGV = 1; RDGV + somatic LOH or biallelic RDGV=2
III. **Recessive**: no RDGV = 0; RDGV + somatic LOH or biallelic RDGV = 1: RDGV without somatic LOH = excluded sample.

Taken together, 3 models of inheritance were tested with 5 different RDGV sets, making up in total 15 models to test across 12 different cancer types and pancan. In total, 15*12*29 = 5,655 model scenarios could have been tested at most. Ultimately, 4,693/5,655 scenarios were tested in the discovery phase.

#### Estimation of Effect Sizes via Burden Testing

Since no effect sizes were reported in SKAT-O, we also performed gene-based burden testing (aggregating variants occurring in the same gene) applying the same models as above. Association testing was performed via linear regression with the *lm* function of the R base package stats in R 3.5.0 as follows:

• Somatic Component ∼ Gene + Covariates

The somatic components were coded as quantitative variables as described above. The gene variable was encoded as a binary categorical variable depending on the model of inheritance (additive, recessive, dominant). When applicable, we controlled for age of diagnosis, sex, cancer type and ancestry (first 6 PCs) as covariates. In total, burden testing was performed for each scenario which was also tested via SKAT-O.

#### Quantile-Quantile Plots for Quality Control

To check for potential biases in testing, we plotted quantile-quantile plots (QQ-plots) for each somatic component tested for each scenario (model of inheritance, RDGV set) in the respective cancer type and calculated the corresponding inflation factor λ. For the QQ-plots, the expected p-value was calculated by ranking all tested genes and dividing the rank of a gene by the total number of genes tested. The idea behind the QQ-plots was that most genes were expected to not have an effect on a somatic component and thus, most p-values would be distributed randomly and fall on a linear line when ordered. The inflation factor λ was calculated to check for inflation, which would be indicated by λ > 1. The inflation factor λ was estimated by dividing the median of the chi-squared test statistic of the p-values by the expected median of the chi-squared distribution, which would be a chi-squared distribution with one degree of freedom. QQ-plots with no inflation would have an inflation factor of λ ≈ 1 and deflated QQ-plots would have an inflation factor of λ < 1. Ultimately, we excluded model scenarios in which at least 100 genes were tested and the inflation factor was ≥ 1.5 (19 ot ouf 1,909).

#### Estimation of False Discovery Rates

We calculated false discovery rates (FDRs) via two approaches: empirical FDR and via a randomized set of genes. To estimate the empirical FDR, the somatic component matrix (somatic components as columns and sample IDs as rows) was randomly shuffled within each cancer type. Importantly, the link between individuals and somatic components was broken down, but the correlation structure between components was conserved. Then, with the randomized somatic component matrix, testing was performed in the same way as it was performed before. We calculated empirical FDR thresholds for each cancer type (or pancan) separately. For instance, the p-value at which 1 % of the associations from the randomized run would have been called as a hit (false discovery) corresponds to a FDR of 1 %.

For our second approach, we repeated the whole analysis using 1,000 random genes. We generated a list of genes, which were not in our pre-selected gene list of 892 genes and in which RDGVs according to RDGV set III. were identified in at least 2 samples. In addition, we discarded all genes which were reported to have a physical interaction with any gene from our pre-selected gene list according to the reported physical interactions from STRING v11.5^99^ with a combined score of at least 50 %. Out of 11,408 remaining genes, 1,000 genes were randomly selected and used for testing. Next, we performed the same steps as it was performed for the pre-selected list of genes, including the calculation of empirical FDRs via randomization and the exclusion of model scenarios with high inflation factors (31 out of 1,885). Based on the conservative hypothesis that there would be no real associations from the random list of genes, we calculated FDRs at different empirical FDR thresholds by dividing the number of hits, which were detected via the random list of genes by the number of genes detected at the same empirical FDR with our pre-selected list of genes. For instance, at an empirical FDR of 1% we identified 44 hits with our random list of genes and 207 hits with our pre-selected list of genes. Thus, we estimated a FDR of 44/207 ≈ 21 % at our empirical FDR of 1 %.

#### Identification of Associations in the Discovery Cohort and Re-Testing in the Validation Cohort

Hits were identified in the discovery cohort when they were significant either at a FDR of 1 % or 2 % based on the estimation of the empirical FDR. These were then re-tested in the matching cancer type based on the tissue of origin (Supplementary Table 5). In total, for 12 individual cancer types a matching cancer type based on the tissue of origin was available in the validation cohort with a sample size of at least 50 samples: bladder cancer, brain glioma multiforme, low-grade glioma, breast cancer colorectal cancer, kidney cancer, lung adenocarcinoma, lung squamous carcinoma, ovary cancer, prostate cancer, skin cancer, stomach and esophagus cancer. Hits which were identified with all cancer types together (pancan) were re-tested in the validation cohort in the same way. We called a hit as replicated when it reached the empirical FDR of either 1 % or 2 % and had the same estimate effect direction as in the discovery cohort. Effect size directions were extracted from the performed burden tests.

#### Network analysis

For the network analysis, we downloaded protein network data from STRING v11.5^99^ involving only physical links, and from HumanNet^90^ v3 the functional gene network (HumanNet-FN). From STRING we only kept interactions which had a combined confidence score (based on experimental, database, and text mining) of at least 80%. The following steps were performed for each protein network separately.

Firstly, we extracted all interactions which involved interactions between genes from our pre-selected gene list of 892 genes. We calculated the total number of interactions our replicated genes had at an empirical FDR of 1 % with each other. It was tested via randomization whether this number was higher than one would expect at random. For this purpose, we selected randomly the same number of genes and calculated the total number of interactions these genes had with each other. We controlled for the total number of interactions each gene had, since some genes (e.g. *BRCA1*) have in general a lot of physical interactions, which would confound our results. To control for this, we counted the total number of interactions our replicated genes had, split them into 10 equal sized bins, assigned all our pre-selected genes a bin, and then randomly selected the same number of genes from each bin. Randomization was performed 1,000 times.

Next, we counted how many genes, which only replicated at an FDR of 2%, had at least one interaction with a gene which replicated at an FDR of 1%. Here, we applied the same approach. We counted the total number of interactions each gene, which only replicated at an FDR of 2 %, had in total and split the number of interactions into 10 equal sized bins. Each gene from our list of genes was assigned a bin and then we randomly selected 1,000 times the same number of genes from each bin and performed the same calculation.

#### Calculation of Frequency of RDGVs in Length Matched Randomly Selected Genes

To calculate the number of RDGVs occuring in a control set of genes, we matched each replicated gene randomly with a gene covering the same length. For this purpose, we intersected the TCGA coverage file with the reported exonic coordinates provided by NCBI RefSeq^134^ track hg38. We only considered protein-coding genes. The covered length of each gene was calculated in kilobases and each replicated gene was randomly matched 10 times with a gene, which covered the same length in our data. Subsequently, RDVGs based on different sets were counted in the replicated gene sets as well as in the length matched control genes. For the validation cohort PCAWG_Hartwig-WGS, the same approach was applied. Here, the coordinates from the CRG75 alignability track were intersected with the exonic coordinates provided by NCBI RefSeq^134^ track hg19 to determine the length of the coding region for a gene.

## Data Availability

Data sources used are detailed in the Methods section.

## Code Availability

Code can be obtained from https://github.com/lehner-lab/RDGVassociation and data visualized at https://mischanvp.shinyapps.io/rare_association_shiny/ once manuscript is accepted for publication.

## Acknowledgements

MVP was supported by a Spanish Ministry of Science and Innovation FPI Fellowship (ref.: PRE2018-084410). Work in the lab of BL is funded by European Research Council (ERC) Advanced (883742) and Consolidator (616434) grants, the Spanish Ministry of Science and Innovation (BFU2017-89488-P, EMBL Partnership, Severo Ochoa Centre of Excellence), the Bettencourt Schueller Foundation, the AXA Research Fund, Agencia de Gestio d’Ajuts Universitaris i de Recerca (AGAUR, 2017 SGR 1322), and the CERCA Program/Generalitat de Catalunya. Work in the lab of FS is funded by the ERC Starting Grant HYPER-INSIGHT (757700), the Horizon2020 grant DECIDER (965193), Spanish Ministry of Science and Innovation grant REPAIRSCAPE (PID2020-118795GB-I00), the EMBO YIP program, and the Severo Ochoa Centre of Excellence award to IRB Barcelona, and the CERCA Program/Generalitat de Catalunya. We thank Dr. Solip Park for method consultations, comments, and discussions.

## Author contributions

MVP designed and implemented the methods, performed all analyses and visualized and interpreted the data. BL and FS conceived the study, designed the methods and supervised the work. All authors drafted the manuscript.

