## Supplementary Note, Figs and Tables for "The impact of rare germline variants on human somatic mutation processes"

**3 Bibliography** **28**

15 Distribution of all 29 somatic components in PCAWG_Hartwig-WGS 13
16 Pearson correlation between IC9 (Sig.11+19) and telomere features 14
17 Identification of individuals with outlying total number of variants, outlying heterozygosity rate or high relatedness in TCGA-WES. . . 15
18 Identification of individuals with outlying total number of variants, outlying heterozygosity rate or high relatedness in PCAWG_Hartwig-WGS . . . . . . . . . . . . . . . . . . . . . . . 16
19 Principal component analysis on common germline variants in TCGA-WES . . . . . . . . . . . . . . . . . . . . . . . . . . . 17
20 Principal component analysis on common germline variants in PCAWG_Hartwig-WGS . . . . . . . . . . . . . . . . . . . . . . . 18

21 Extraction of European individuals in TCGA-WES . . . . . . . . . . 19
22 Extraction of European individuals in PCAWG_Hartwig-WGS . . . 20
23 Hierarchical clustering of replicated genes at FDR of 1 % using cancer dependency map CRISPR screen data. . . . . . . . . . . . . . . 21
24 Hierarchical clustering of replicated genes at FDR of 2 % using cancer dependency map CRISPR screen data. . . . . . . . . . . . . . . 22

### List of Tables

1 Covered Genomic Regions with WES and WGS Masks . . . . . . 23
2 Somatic Component Names . . . . . . . . . . . . . . . . . . . . . . . . 24
3 TCGA Study Abbreviation . . . . . . . . . . . . . . . . . . . . . . . . 25
4 PCAWG Study Abbreviation . . . . . . . . . . . . . . . . . . . . . . . 26
5 Cancer Type Names . . . . . . . . . . . . . . . . . . . . . . . . . . . 26
6 Overview of sample sizes . . . . . . . . . . . . . . . . . . . . . . . . 27

### 1 Supplementary Figures

#### 1.1 Somatic Input Features

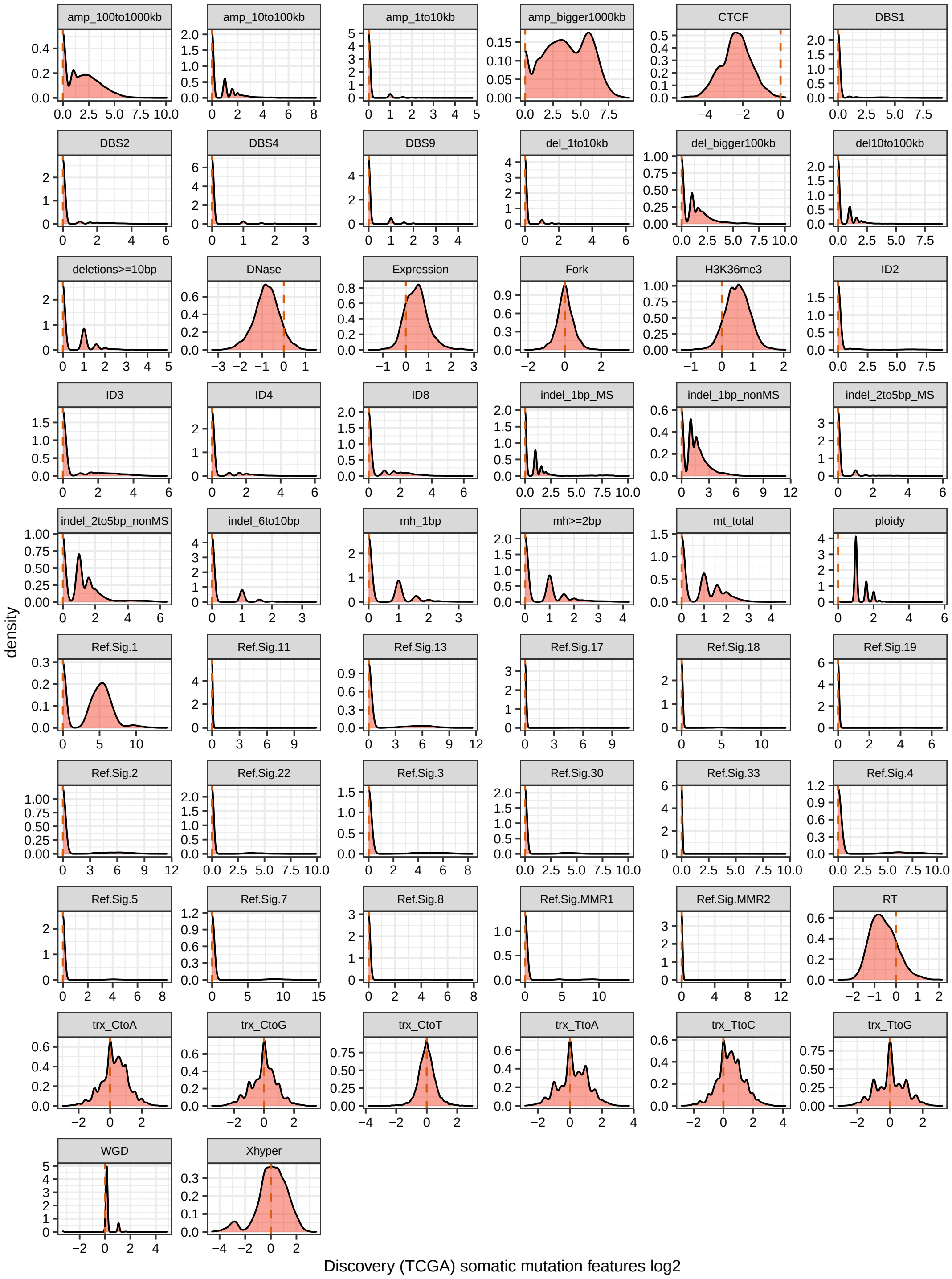

**Figure 1: Distribution of all 56 somatic features in TCGA-WES.** Fork: replicative strand bias, RT: replication timing, trx: transcription strand bias, Xhyper: Chromosome X hypermutation.

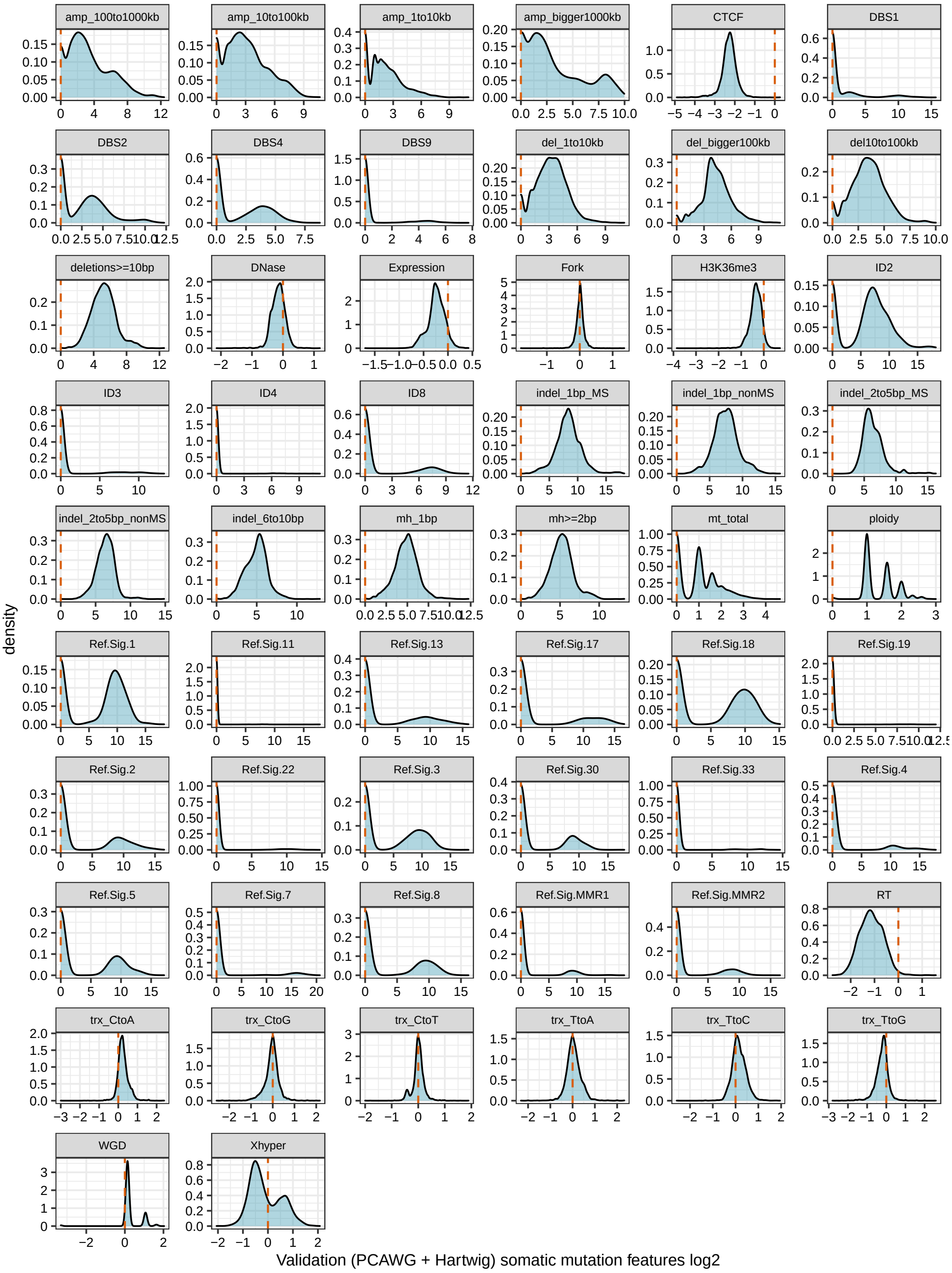

**Figure 2: Distribution of all 56 somatic features in PCAWG_Hartwig -WGS.** Fork: replicative strand bias, RT: replication timing, trx: transcription strand bias, Xhyper: Chromosome X hypermutation.

#### 1.2 Extraction of Independent Components

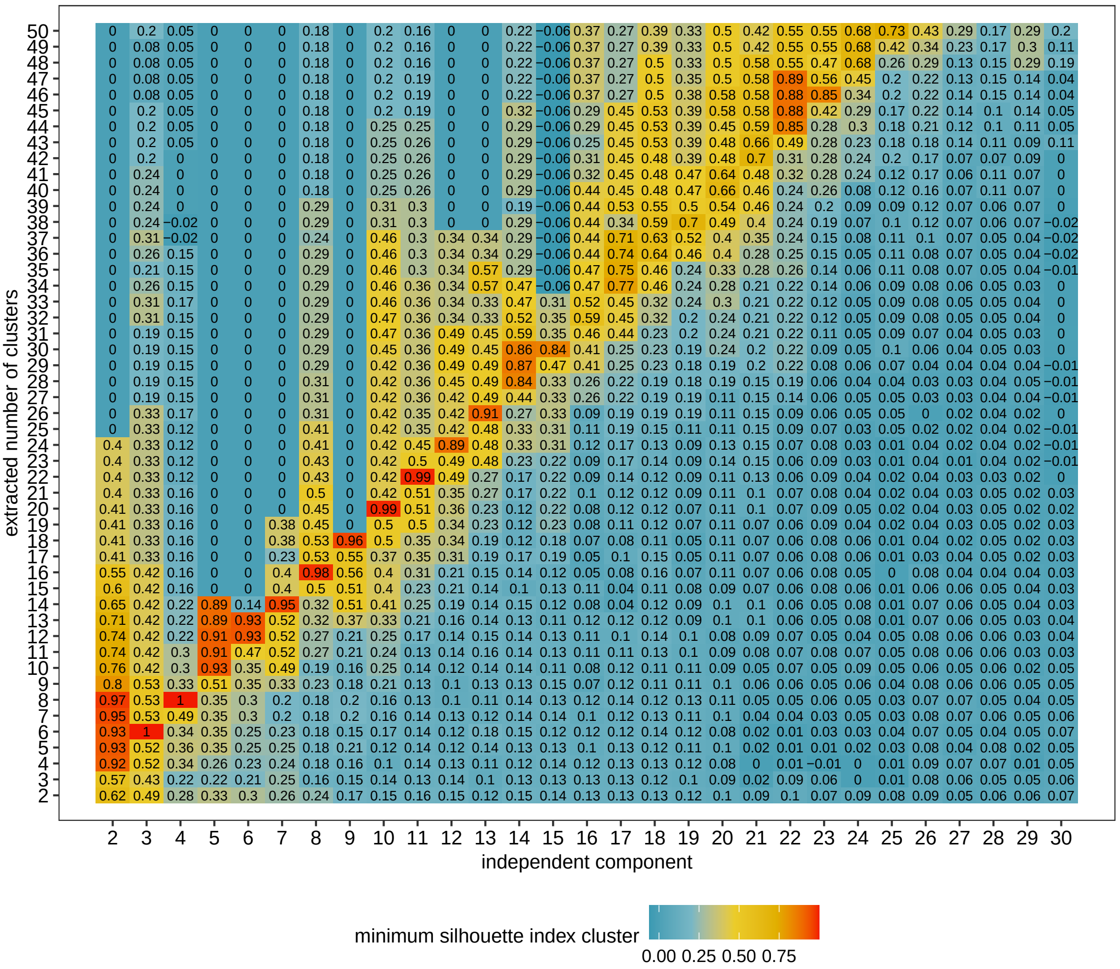

**Figure 3: Finding the optimal number of independent components.** Independent component extraction was run with increasing number of independent components (x-axis). Each extraction was performed 200 times and then clustered using k-medoid clustering with increasing number of clusters (y-axis). Color code and number in each tile shows the minimum silhouette index of a cluster.

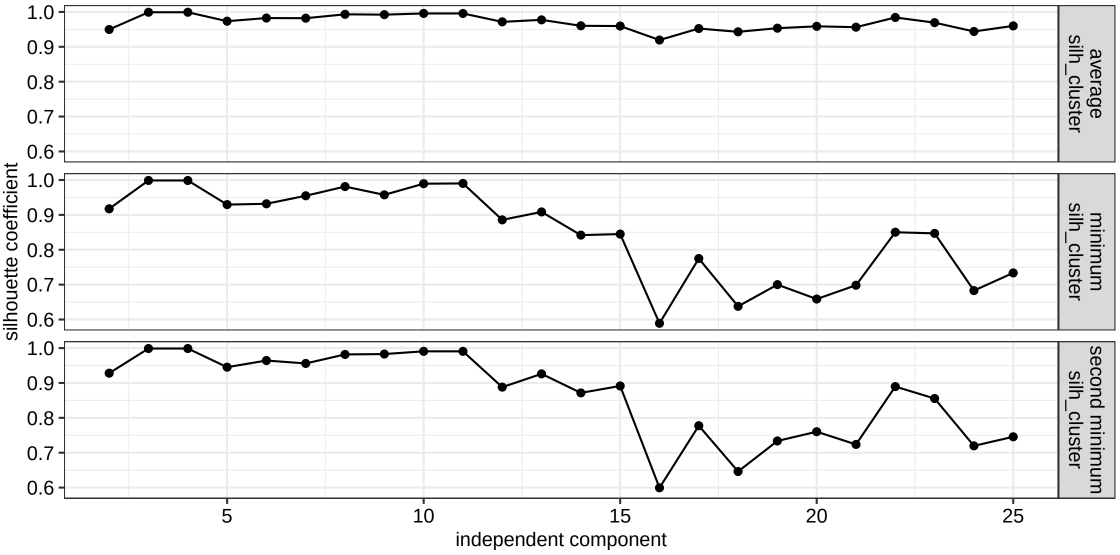

**Figure 4: Selection of 15 independent components for further analysis.** Showing the average, minimum and second minimum silhouette index of the clusters when extracting 2 times more clusters for a set number of components.

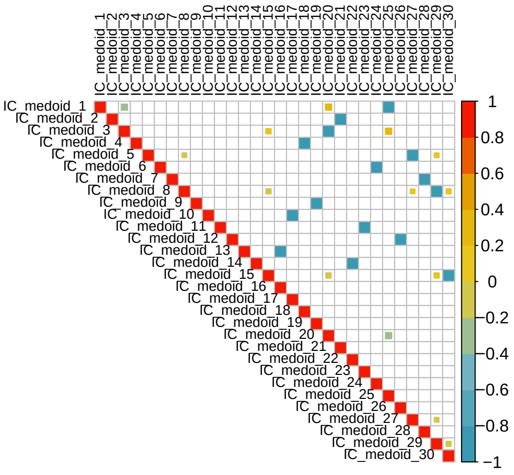

**Figure 5: Pearson correlations between all 30 independent components which were extracted using 15 components and k-medoid clustering with k = 30.** Each component occurred twice with opposite signs.

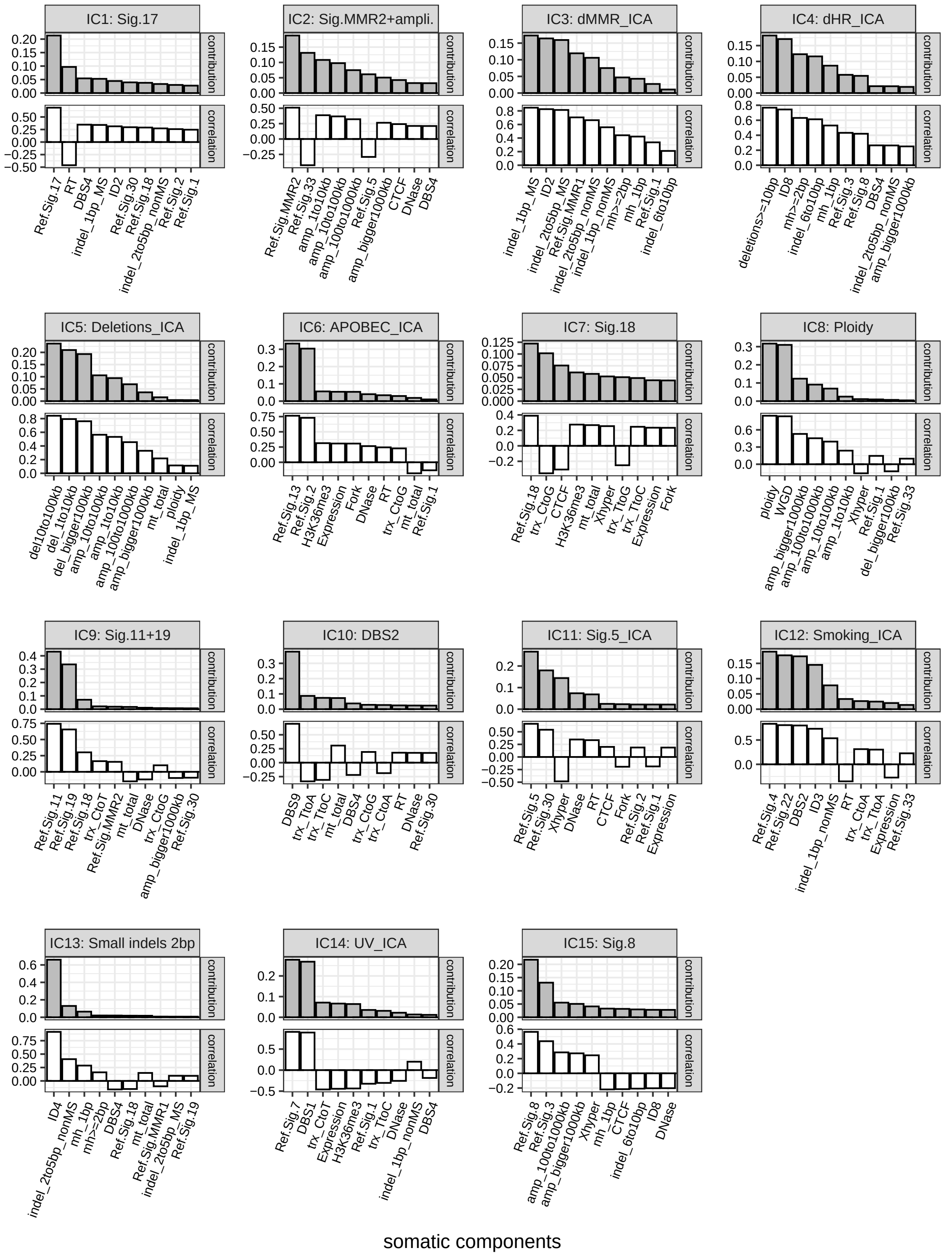

**Figure 6: Overview of strongest contributing features to the independent components.** Showing the Pearson correlation (white bars) and contribution (fraction of 1) (grey bars) of the 10 strongest somatic features to the respective components. Fork: replicative strand bias, RT: replication timing, trx: transcription strand bias, Xhyper: Chromosome X hypermutation. Components were renamed based on strongest correlating somatic features.

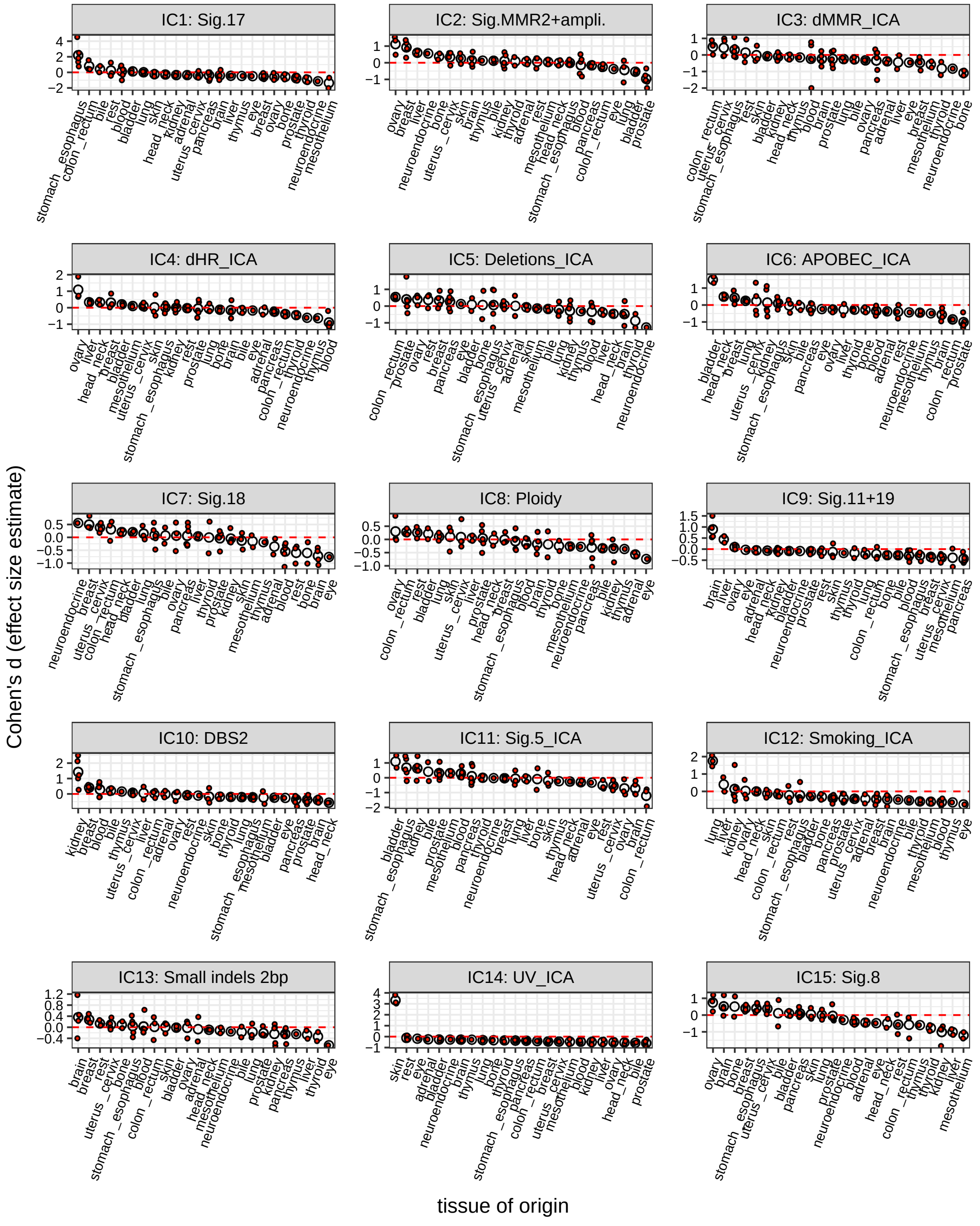

**Figure 7: Several independent component scores were enriched in specific tissue of origins.** Cohen's *d* (effect size estimate) was calculated for each cancer type, grouped by tissue of origin and, then the average value was estimated. Average effect size estimates were ordered by decreasing value for each independent component. Components were renamed based on strongest correlating somatic features.

#### 1.3 Extracting Components using Variational Autoencoders

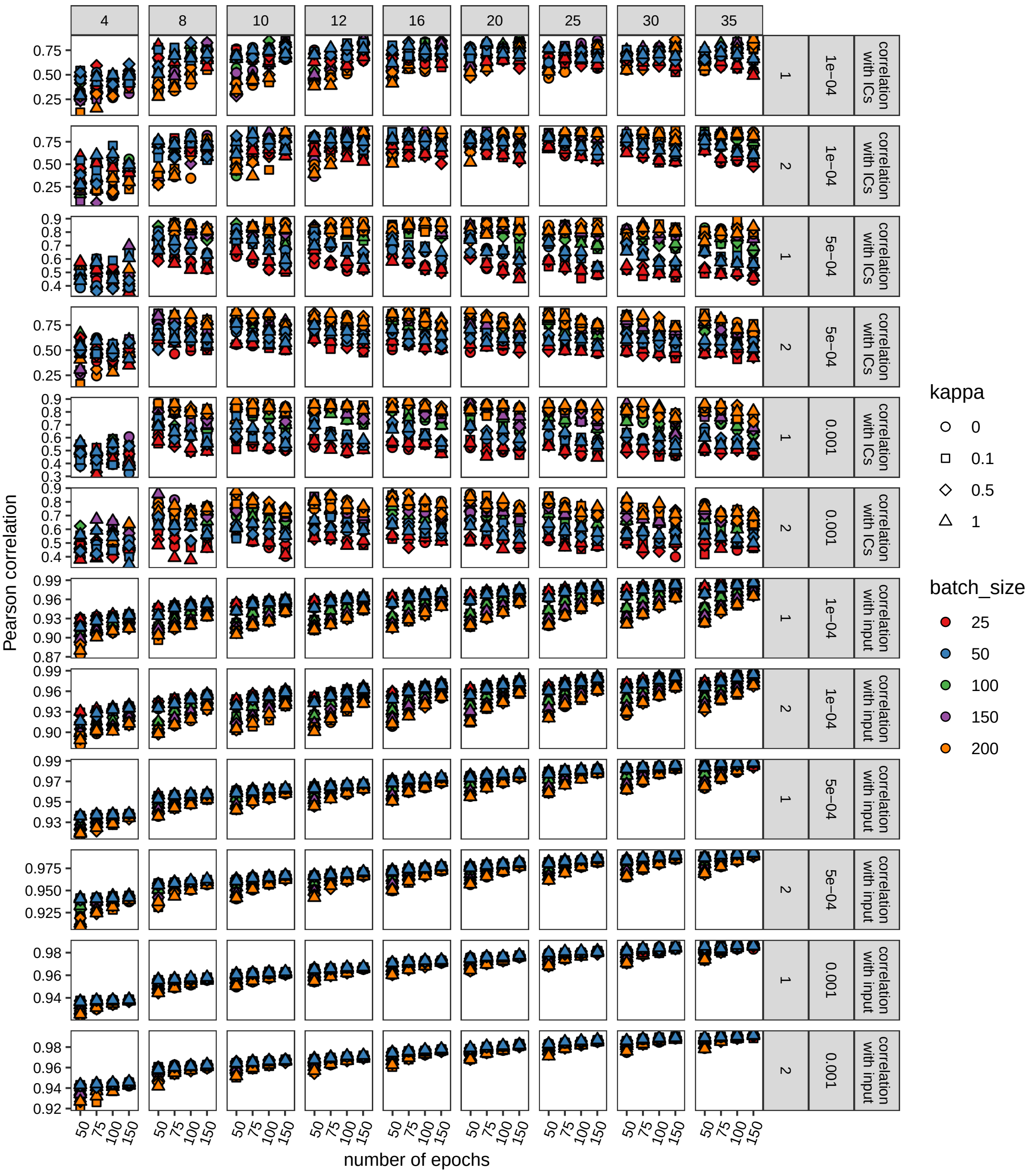

**Figure 8: Finding the optimal hyperparameters for the variational autoencoder.** Increasing number of epochs (x-axis), different kappa factors (point shape), increasing batch sizes (point color), three different learning rates (0.001, 0.0005, and 0.0001), different number of hidden layers between latent space and input/output (either 1 or 2) and different number of components (4, 8, 10, 12, 16, 20, 25, 30, and 35) were tested. Evalutation by measuring the average Pearson correlation with 4 ICs (UV, smoking, dMMR, and dHR ICs) and by calculating the Pearson of the reconstructed input with the initial input (y-axis).

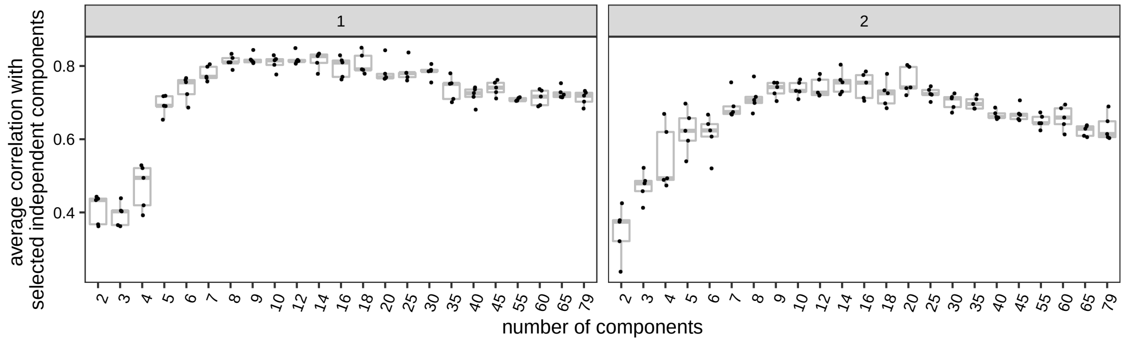

**Figure 9: Correlation with biologically relevant components increased with increasing number of component extractions and quickly reached saturation.** Component extractions were run 5 times for each set component number with different random initiations. Number of components (x-axis) are shown against the average Pearson correlation with 4 biologically relevant IC components (UV, smoking, dMMR, dHR). Facet for either using 1 hidden layer or 2 hidden layers between latent space and input/output.

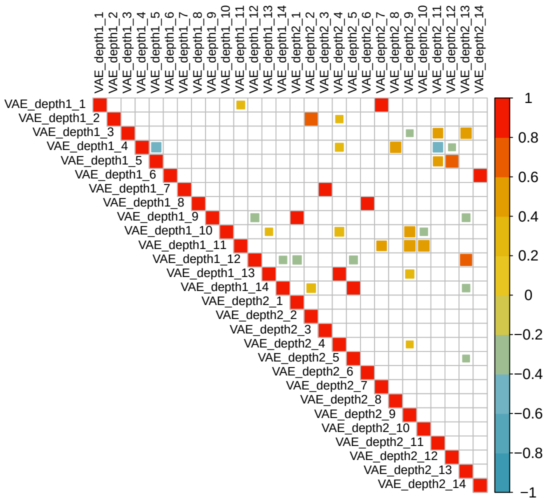

**Figure 10: Number of hidden layers barely made a difference on the extracted components in the latent space of the variational autoencoder.** Pearson correlation between the 14 extracted components with the variational autoencoder using either 1 hidden layer (depth = 1) or 2 hidden layers (depth = 2) between latent space and input/output.

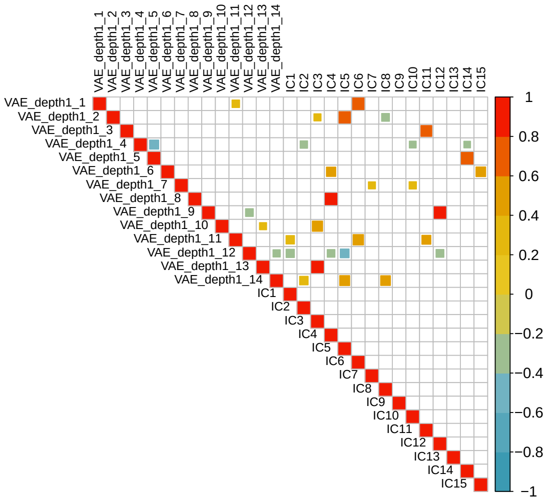

**Figure 11: Some VAE-derived components were not captured in the independent component analysis.** Showing the Pearson correlation between the VAE-derived components using 1 hidden layer (depth = 1) and the ICs.

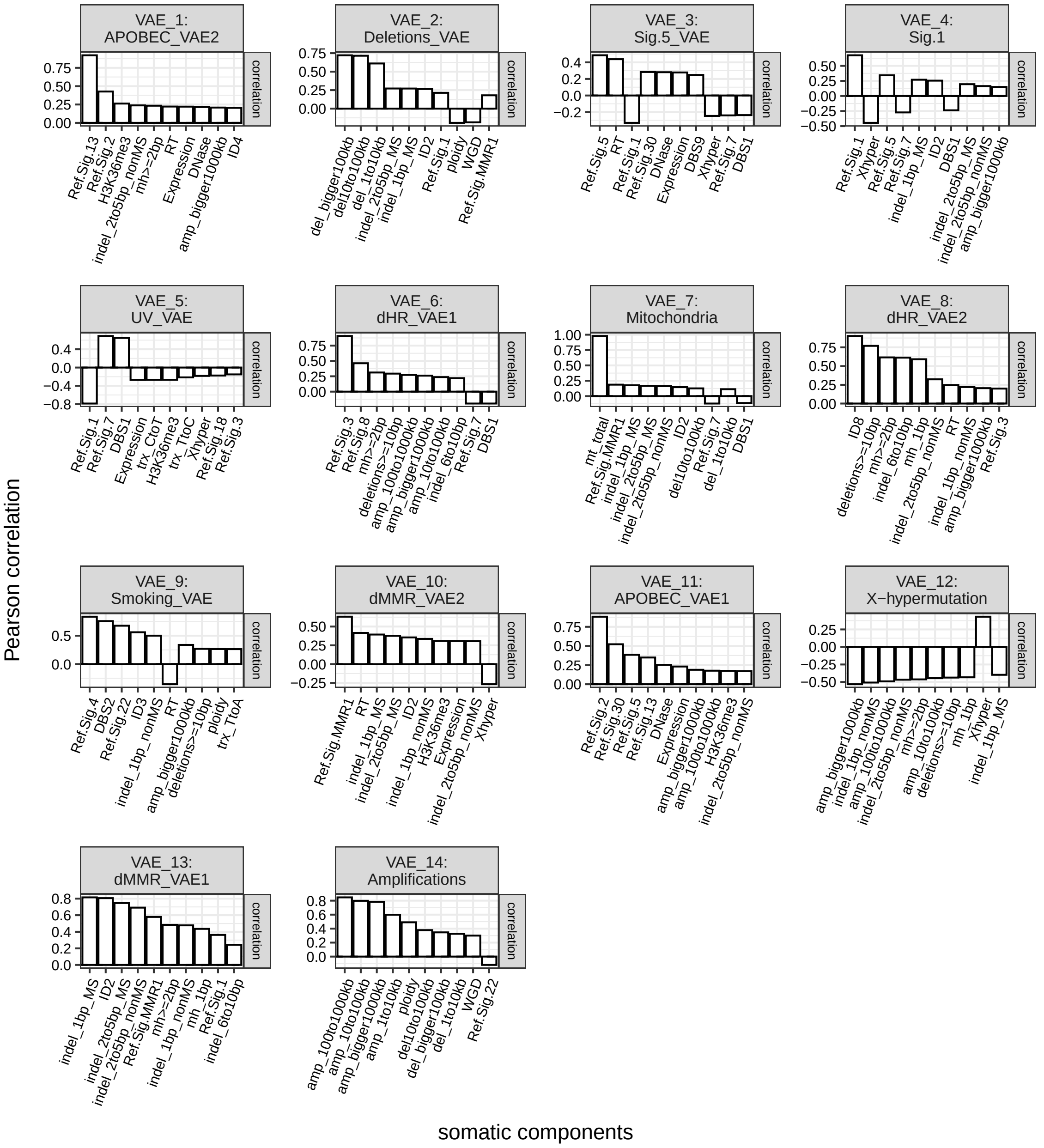

**Figure 12: Overview of strongest contributing features to the variational autoencoder derived components.** Showing the Pearson correlation (white bars) of the 10 strongest somatic features to the respective components, which were extracted via 1 hidden layer between latent space and input/output. Fork: replicative strand bias, RT: replication timing, trx: transcription strand bias, Xhyper: Chromosome X hypermutation. Components were renamed based on strongest correlating somatic features.

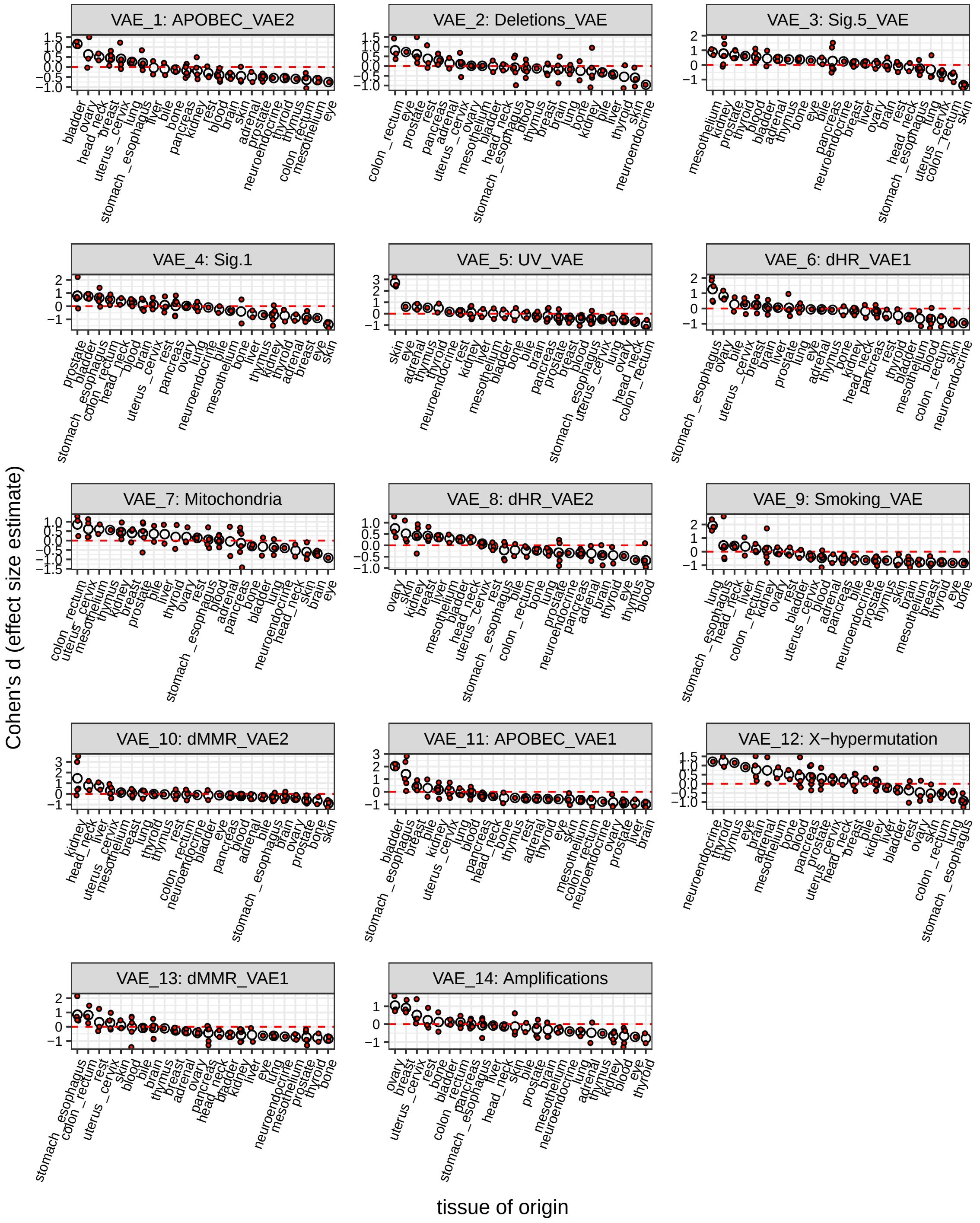

**Figure 13: Several VAE-derived component scores were enriched in specific tissue of origins.** Cohen's *d* (effect size estimate) was calculated for each cancer type, grouped by tissue of origin and, then the average value was estimated. Average effect size estimates were ordered by decreasing value for each VAE-derived component. Components were renamed based on strongest correlating somatic features.

#### 1.4 Overview of 29 Extracted Components

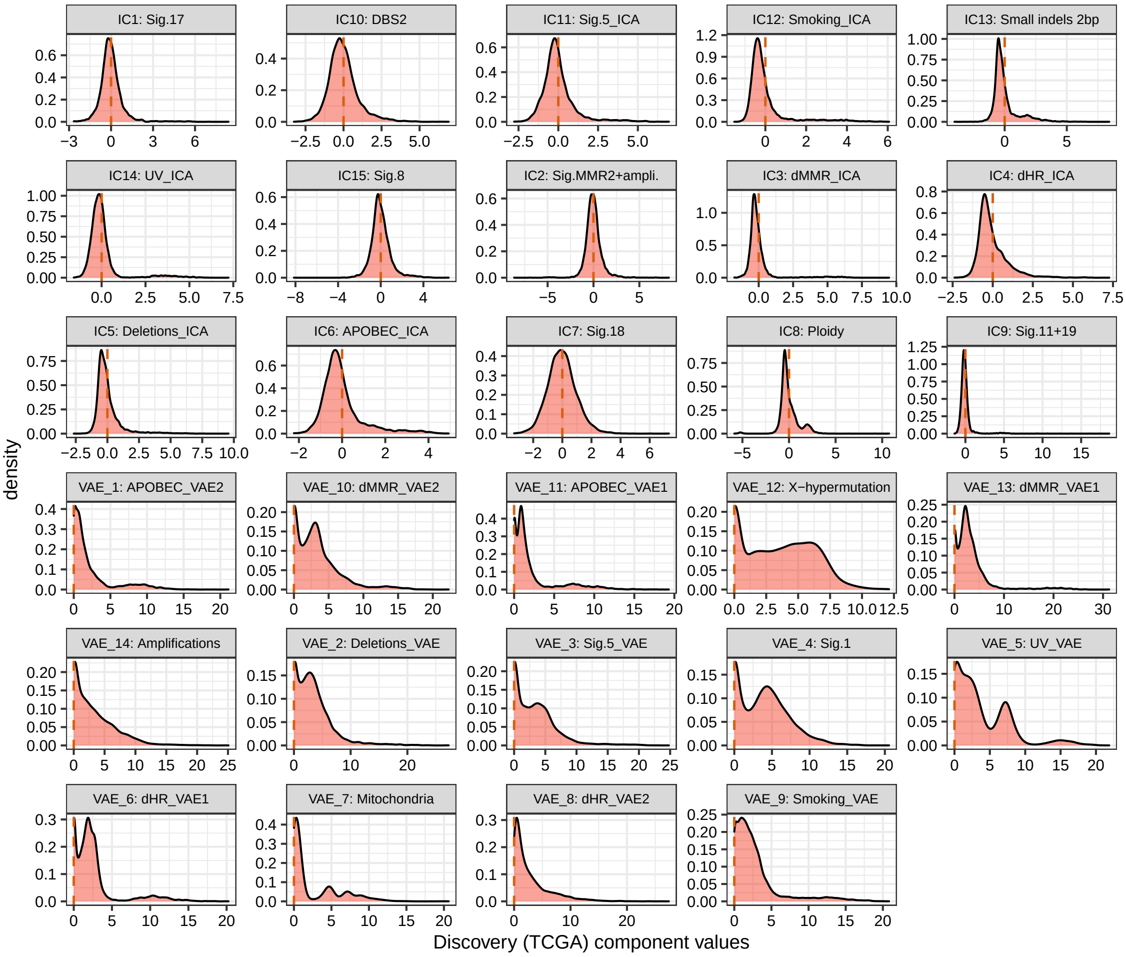

**Figure 14: Distribution of all 29 somatic components in TCGA-WES.**

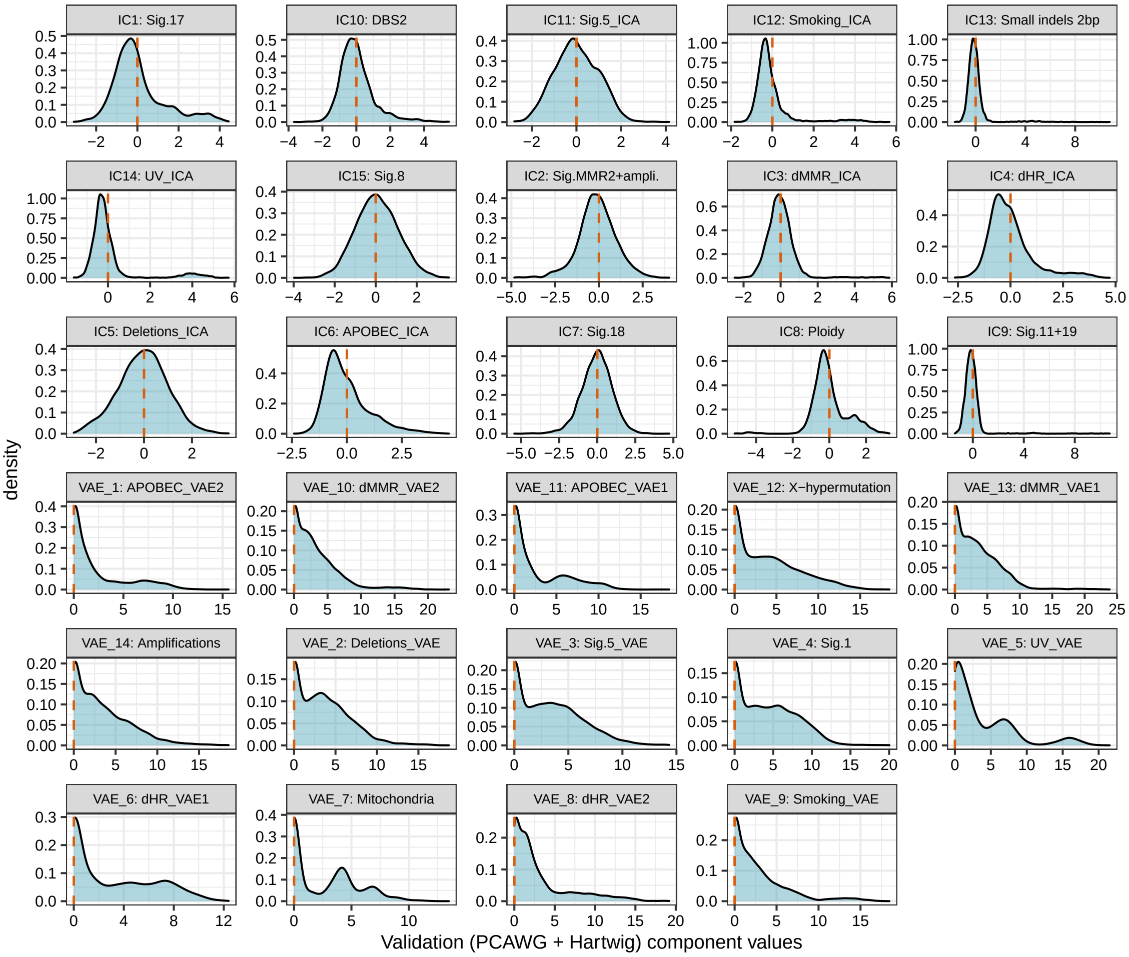

**Figure 15: Distribution of all 29 somatic components in PCAWG_Hartwig -WGS.**

#### 1.5 Correlation between IC9 (Sig.11+19) and telomere features

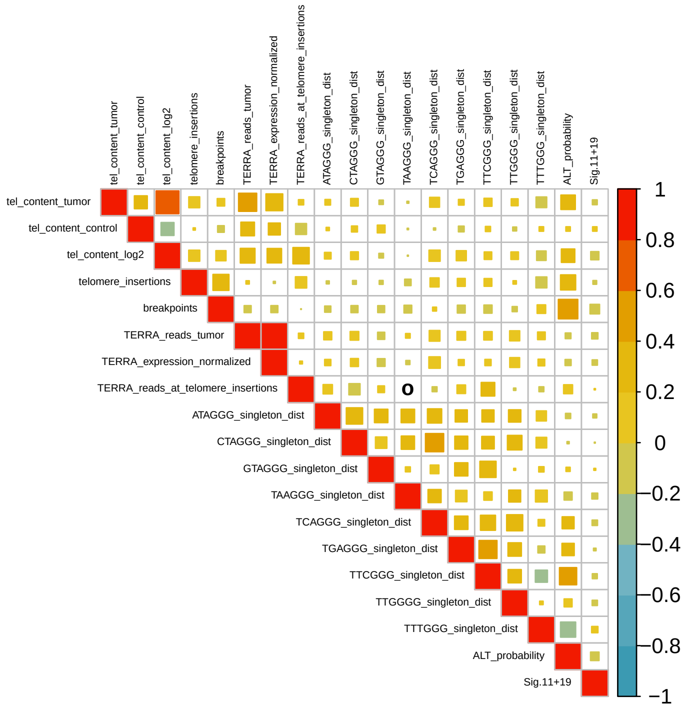

**Figure 16: Pearson correlation between IC9 (Sig.11+19) and telomere features.** Correlations estimated based on 1,254 samples from PCAWG. Telomere features were downloaded from ref[1].

#### 1.6 Sample Level Quality Control and Extraction of Individuals of European Ancestry from Common Germline Variants

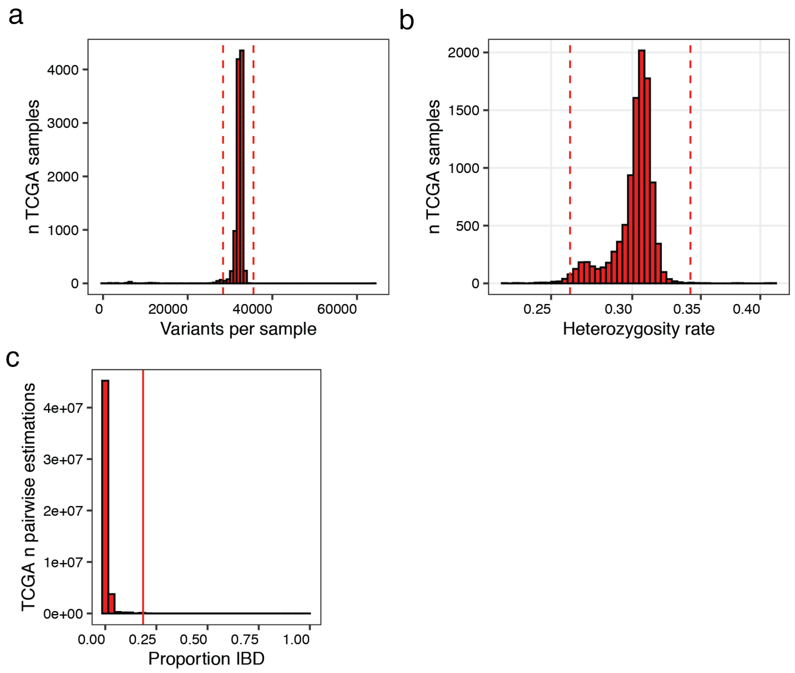

**Figure 17: Identification of individuals with outlying total number of variants, outlying heterozygosity rate or high relatedness in TCGA-WES.** (a) Distribution of total number of variants across samples. Red dashed lines at 1.5 standard deviations away from the mean. (b) Distribution of heterozygosity rate across samples. Red dashed lines at 3 standard deviations away from the mean. (c) Proportion of identity-by-descent (IBD) across all sample pairs. Solid red line at 0.185.

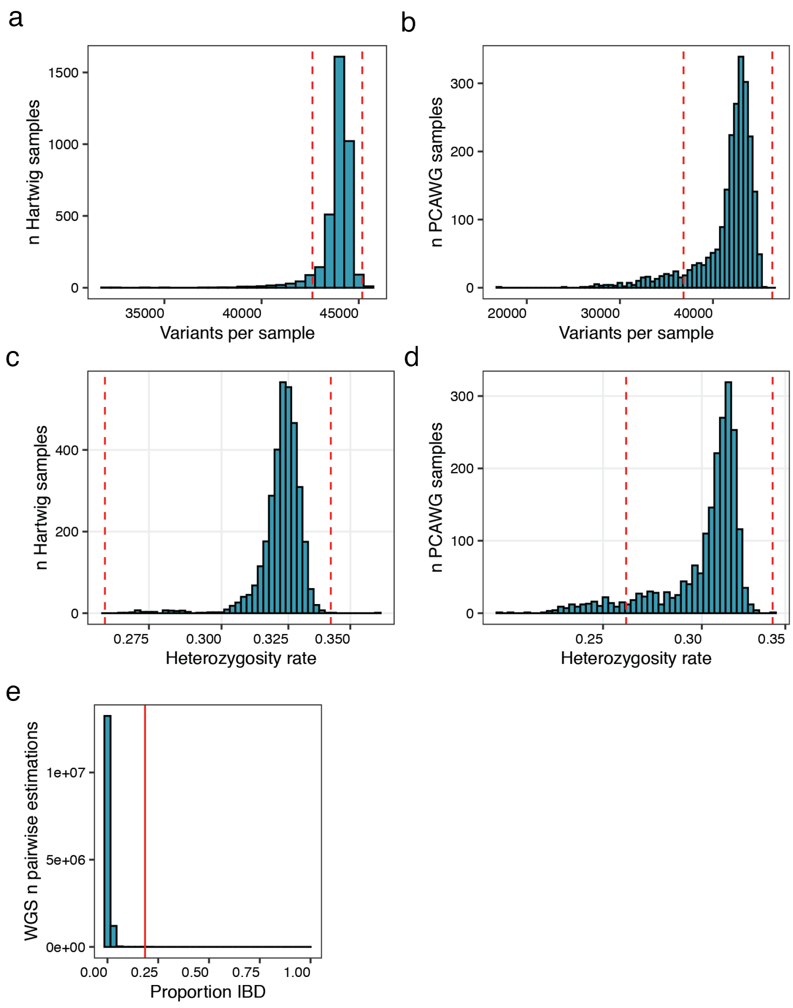

**Figure 18: Identification of individuals with outlying total number of variants, outlying heterozygosity rate or high relatedness in PCAWG_Hartwig -WGS.** Distribution of total number of variants across samples in (a) Hartwig and (b) PCAWG. Red dashed lines at 1.5 standard deviations away from the mean. Distribution of heterozygosity rate across samples in (c) Hartwig and (d) PCAWG. Red dashed lines at 3 standard deviations away from the mean. (e) Proportion of identity-by-descent (IBD) across all sample pairs (PCAWG and Hartwig merged). Solid red line at 0.185.

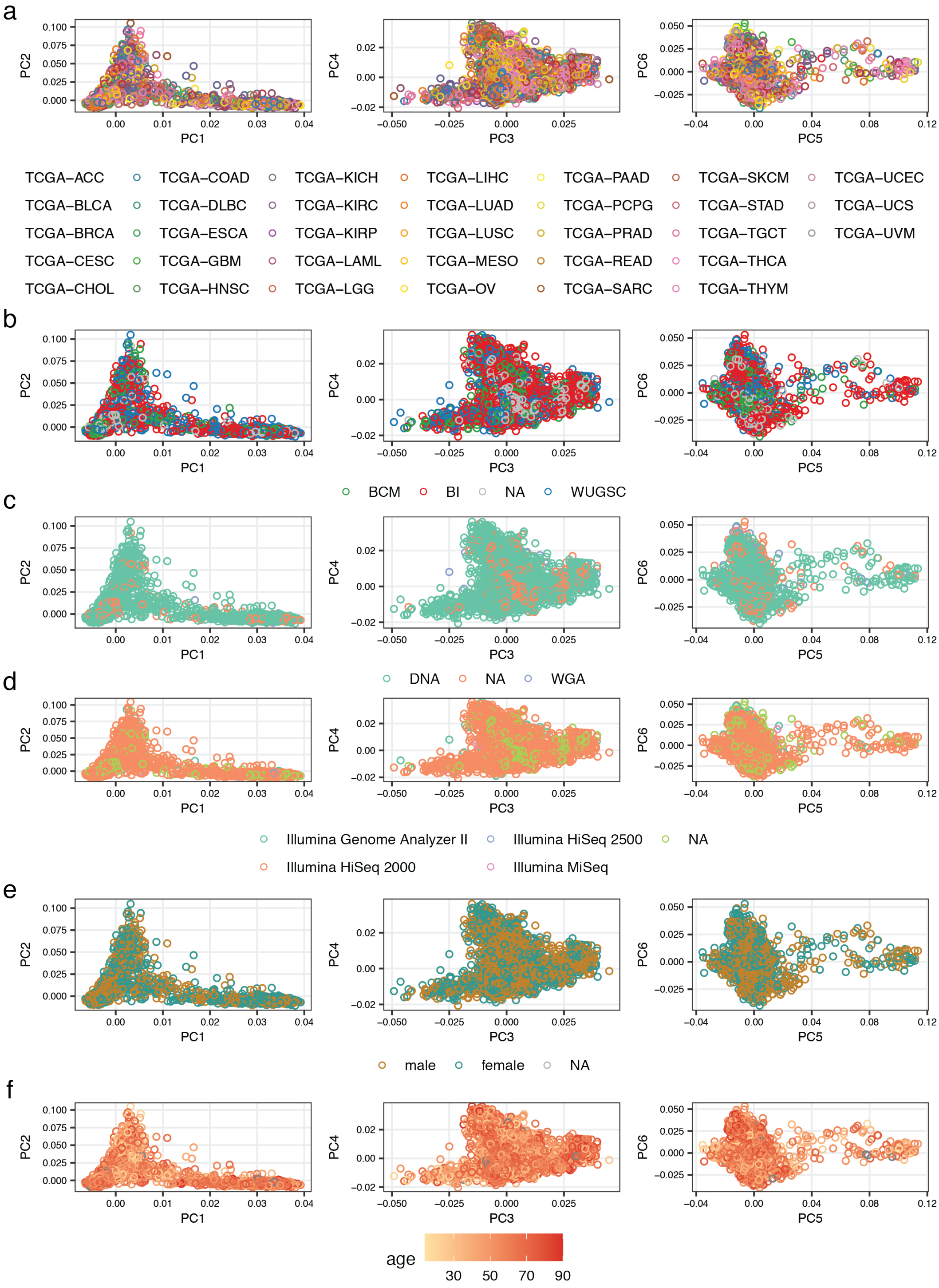

**Figure 19: Principal component analysis on common germline variants in TCGA-WES.** Principal components 1 to 6 color coded by (a) TCGA project id, (b) sequencing center, (c) whole genome amplification (WGA) status prior to sequencing, (d) sequencer, (e) gender, and (f) age of diagnosis.

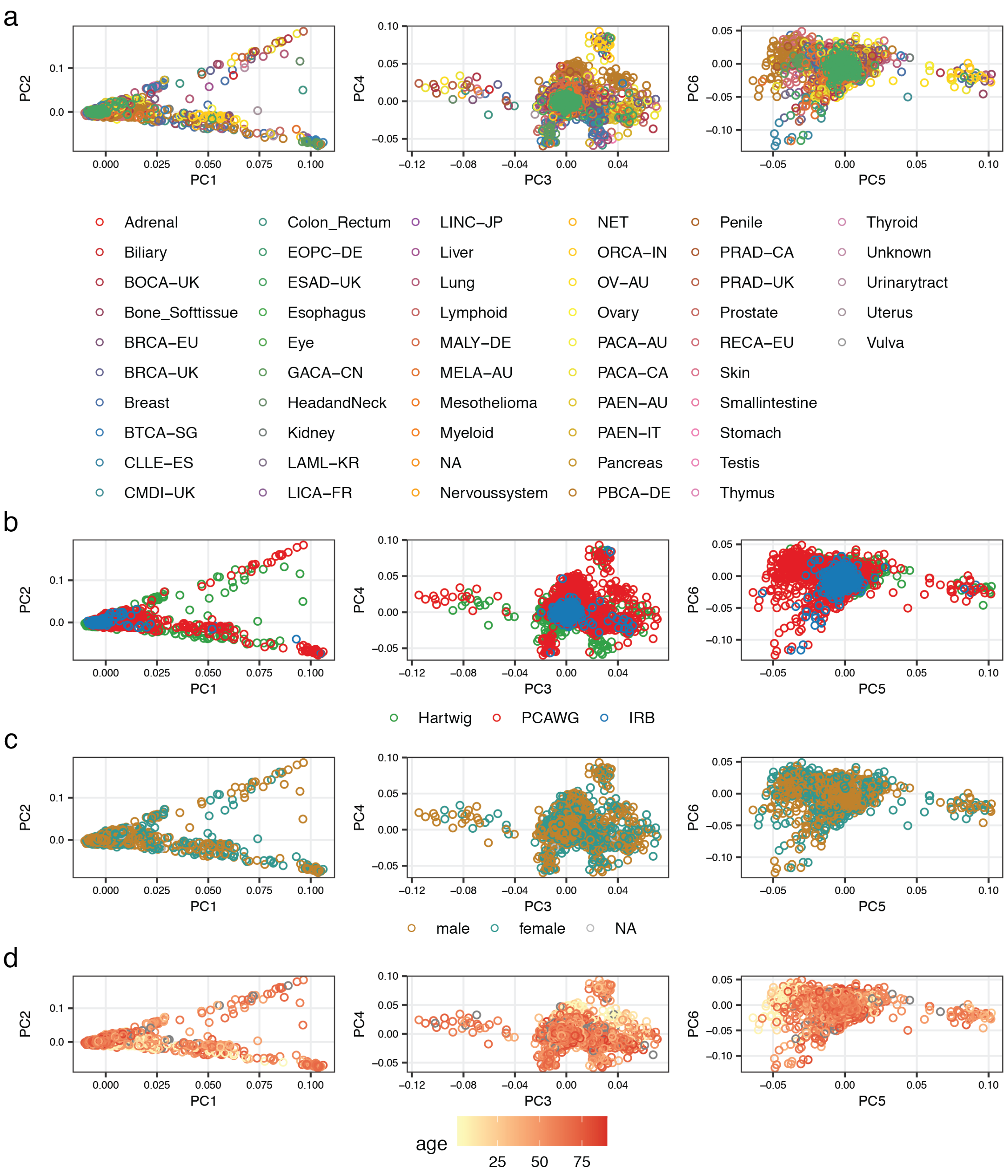

**Figure 20: Principal component analysis on common germline variants in PCAWG_Hartwig-WGS.** Principal components 1 to 6 color coded by (a) PCAWG project id or Hartwig tissue of origin, (b) center/study where germline variants were called, (c) gender, (d) age of diagnosis.

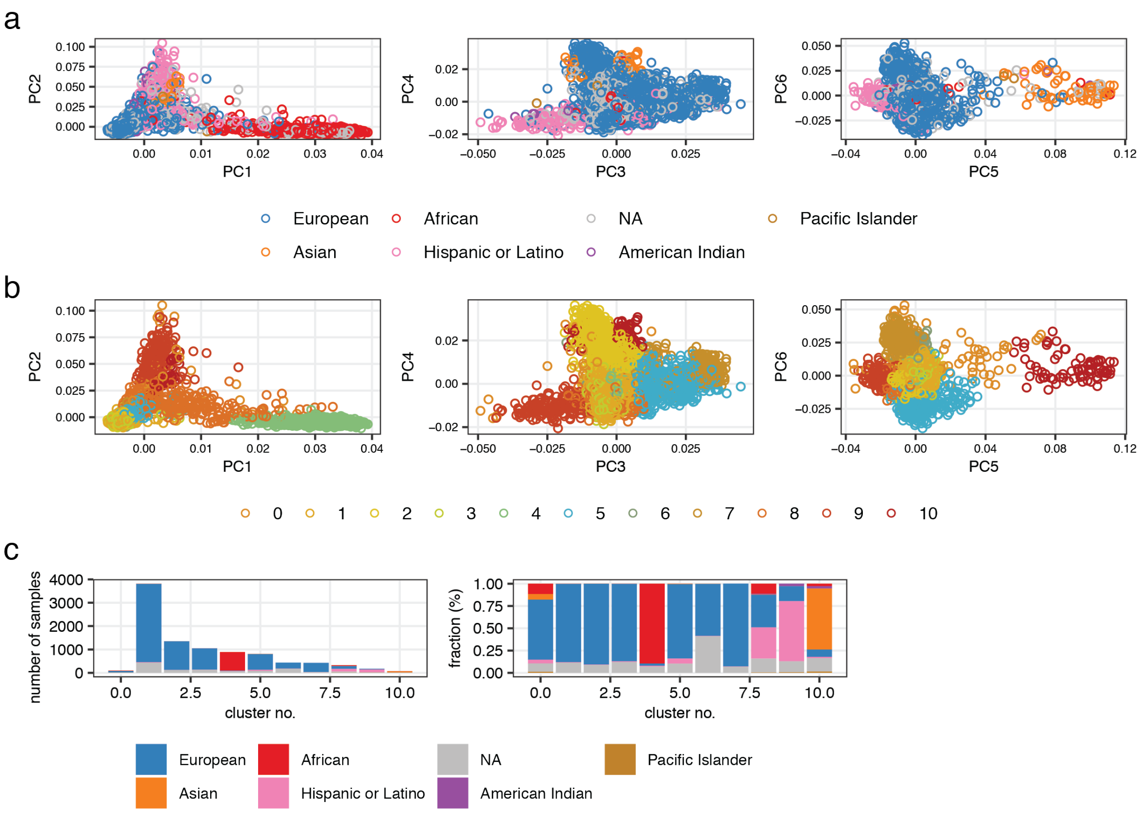

**Figure 21: Extraction of European individuals in TCGA-WES.** Principal components 1 to 6 color coded by (a) reported ethnicities and (b) clustering results using the first 10 principal components. (c) Overview of clustering results. Samples which could not be assigned to a cluster (cluster no. 0) were excluded.

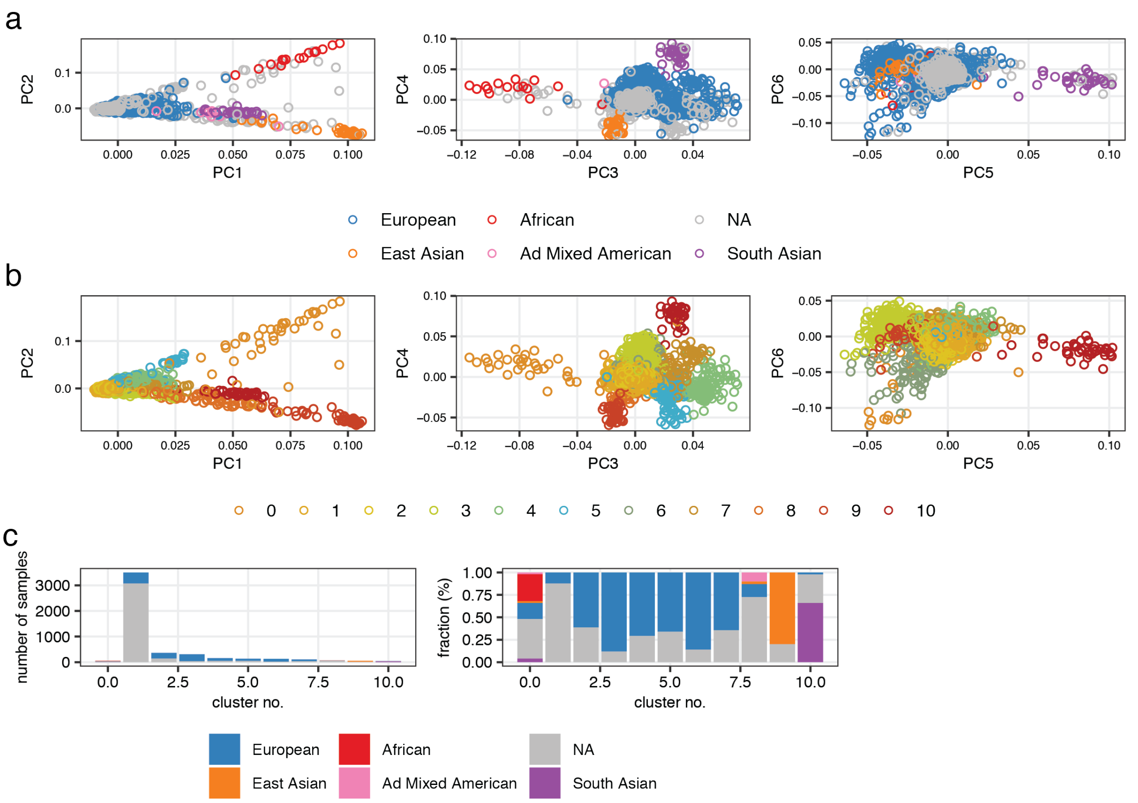

**Figure 22: Extraction of European individuals in PCAWG_Hartwig-WGS.** Principal components 1 to 6 color coded by (a) reported ethnicities and (b) clustering results using the first 10 principal components. (c) Overview of clustering results. Samples which could not be assigned to a cluster (cluster no. 0) were excluded.

#### 1.7 Clustering of Replicated Genes with DepMap

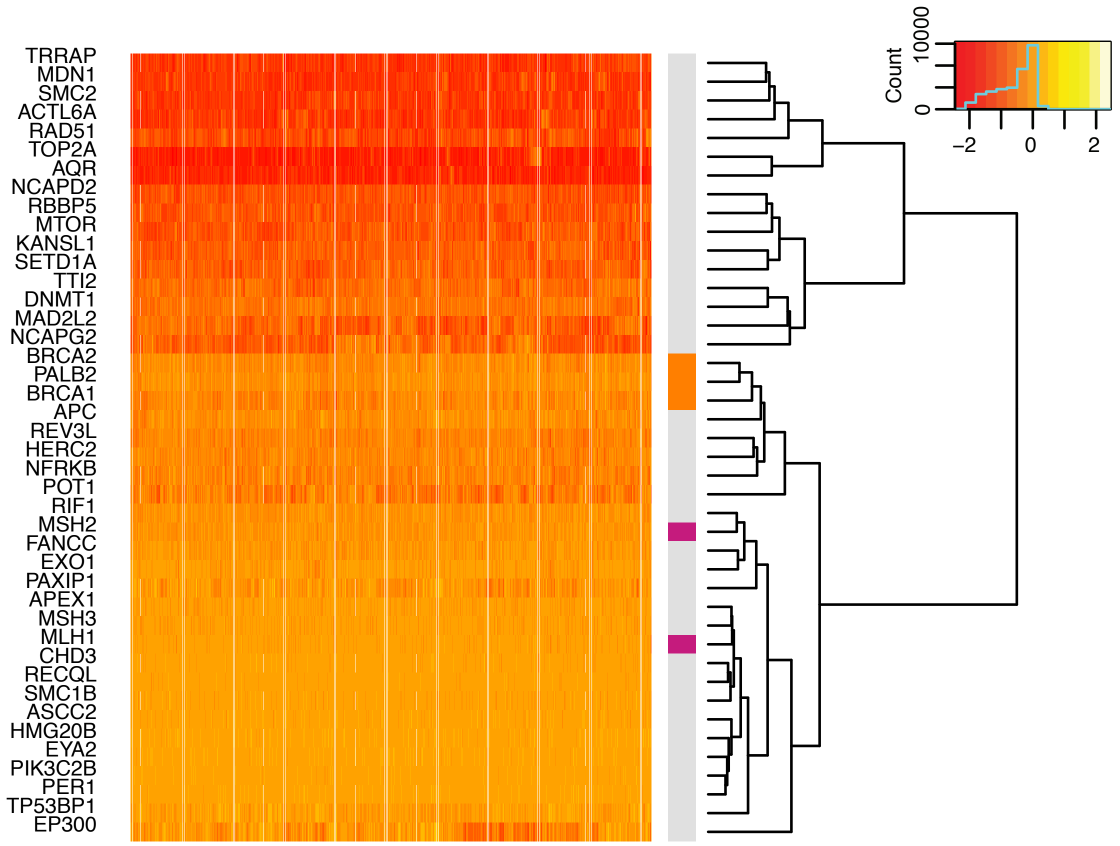

**Figure 23: Hierarchical clustering of replicated genes at FDR of 1 % using cancer dependency map CRISPR screen data.** Estimated fitness effects (Chronos) based on gene knockouts via CRISPR across 1,032 cell lines (columns) from DepMap[2,3] were used as input for clustering. Replicated genes at a FDR of 1 % on y-axis. CRISPR DepMap 21Q3 data was used `https://ndownloader.figshare.com/files/29125323`. Grey strip on right side visualising known dHR genes (orange), known dMMR genes (pink) and novel associations (grey).

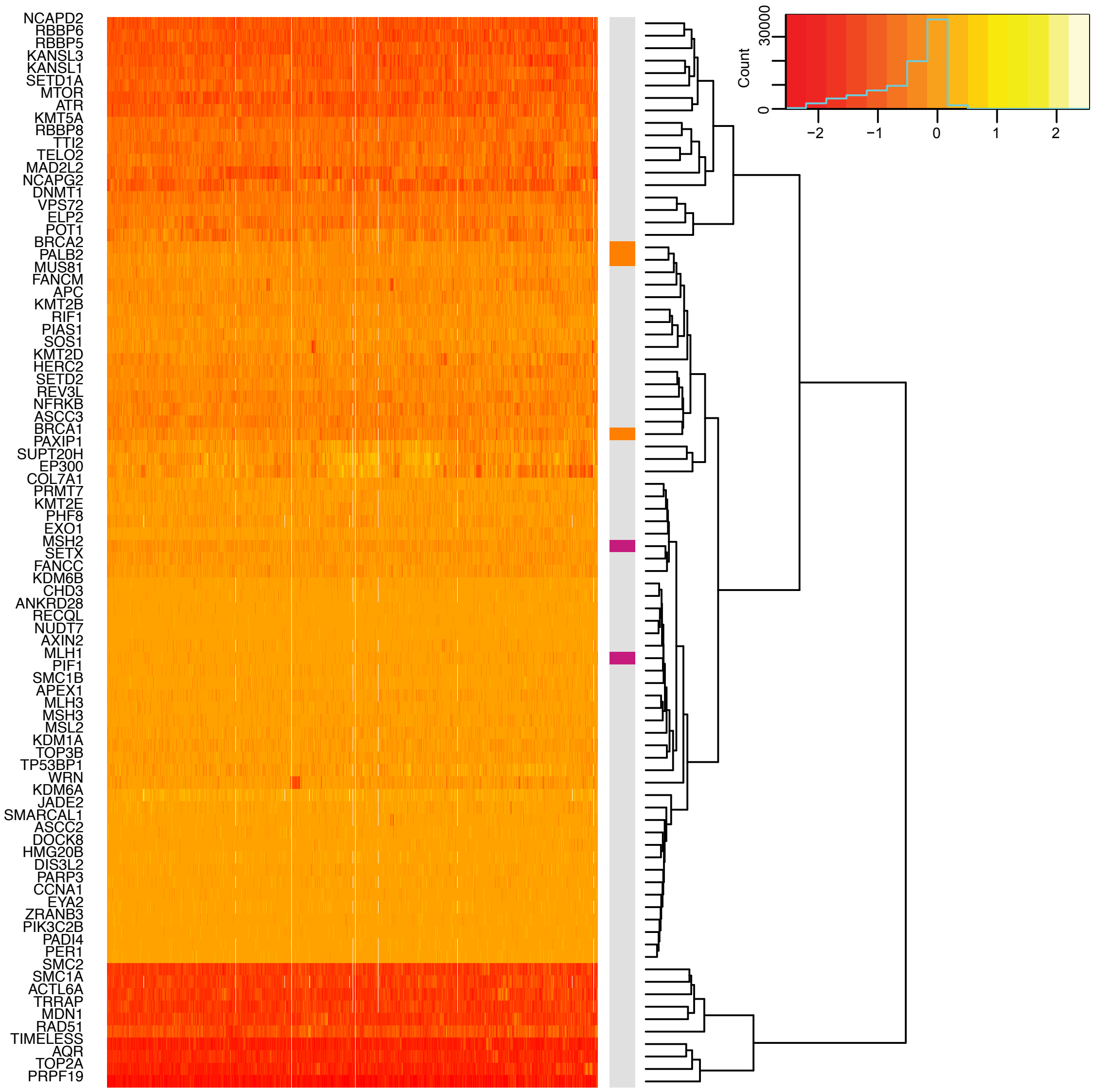

**Figure 24: Hierarchical clustering of replicated genes at FDR of 2 % using cancer dependency map CRISPR screen data.** Estimated fitness effects (Chronos) based on gene knockouts via CRISPR across 1,032 cell lines (columns) from DepMap[2,3] were used as input for clustering. Replicated genes at a FDR of 2 % on y-axis. CRISPR DepMap 21Q3 data was used `https://ndownloader.figshare.com/files/29125323`. Grey strip on right side visualising known dHR genes (orange), known dMMR genes (pink) and novel associations (grey).

### 2 Supplementary Tables

#### 2.1 Genomic Regions

**Table 1: Covered Genomic Regions with WES and WGS Masks.** Estimated lengths of different genomic regions in megabases after applying CRG75 alignability mask on WES and WGS data respectively.

| Region | Bin | Size in WES in Megbases | Size in WGS in Megbases |
|---|---|---|---|
| RT | 1of6 (late) | 1.62 | 345 |
| | 2of6 | 4.03 | 382 |
| | 3of6 | 7.94 | 381 |
| | 4of6 | 12.0 | 379 |
| | 5of6 | 16.7 | 373 |
| | 6of6 (early) | 29.2 | 356 |
| H3K36me3 | 0of5 (no marks) | 71.0 | 3,705 |
| | 1of5 | 6.63 | 143 |
| | 2of5 | 8.54 | 143 |
| | 3of5 | 11.3 | 144 |
| | 4of5 | 16.1 | 147 |
| | 5of5 (high density of marks) | 29.4 | 152 |
| Expression | 0of5 (no expression) | 7.54 | 2,050 |
| | 1of5 | 14.9 | 374 |
| | 2of5 | 17.0 | 425 |
| | 3of5 | 24.2 | 523 |
| | 4of5 | 34.9 | 531 |
| | 5of5 (high expression) | 44.4 | 531 |
| DNase I | 0of5 (no marks) | 95.7 | 3,758 |
| | 1of5 | 6.89 | 137 |
| | 2of5 | 7.68 | 136 |
| | 3of5 | 8.85 | 135 |
| | 4of5 | 9.95 | 134 |
| | 5of5 (high density of marks) | 9.95 | 134 |
| CTCF/cohesin | flanking site $\pm$500 bp | 7.04 | 83.2 |
| | binding site | 1.56 | 17.38 |
| Fork polarity | 1of10 (lagging strand) | 7.71 | 199 |
| | 2of10 | 7.65 | 196 |
| | 9of10 | 6.75 | 195 |
| | 10of10 (leading strand) | 5.54 | 200 |

#### 2.2 Somatic Components

**Table 2: Somatic Component Names.**

| Component | Name |
|---|---|
| IC1 | Sig.17 |
| IC2 | Sig.MMR2+ampli. |
| IC3 | dMMR$_{ICA}$ |
| IC4 | dHR$_{ICA}$ |
| IC5 | Deletions$_{ICA}$ |
| IC6 | APOBEC$_{ICA}$ |
| IC7 | Sig.18 |
| IC8 | Ploidy |
| IC9 | Sig.11+19 |
| IC10 | DBS2 |
| IC11 | Sig.5$_{ICA}$ |
| IC12 | Smoking$_{ICA}$ |
| IC13 | Small indels 2bp |
| IC14 | UV$_{ICA}$ |
| IC15 | Sig.8 |
| VAE_1 | APOBEC$_{VAE2}$ |
| VAE_2 | Deletions$_{VAE}$ |
| VAE_3 | Sig.5$_{VAE}$ |
| VAE_4 | Sig.1 |
| VAE_5 | UV$_{VAE}$ |
| VAE_6 | dHR$_{VAE1}$ |
| VAE_7 | Mitochondria |
| VAE_8 | dHR$_{VAE2}$ |
| VAE_9 | Smoking$_{VAE}$ |
| VAE_10 | dMMR$_{VAE2}$ |
| VAE_11 | APOBEC$_{VAE1}$ |
| VAE_12 | X-hypermutation |
| VAE_13 | dMMR$_{VAE1}$ |
| VAE_14 | Amplifications |

#### 2.3 Cancer Types

**Table 3: TCGA Study Abbreviations.** https://gdc.cancer.gov/resources-tcga-users/tcga-code-tables/tcga-study-abbreviations. List of studies, which were ultimately used in association testing (after filtering steps).

| Study Abbreviation | Study Name |
|---|---|
| ACC | Adrenocortical carcinoma |
| BLCA | Bladder Urothelial Carcinoma |
| BRCA | Breast invasive carcinoma |
| CESC | Cervical squamous cell carcinoma and endocervical adenocarcinoma |
| CHOL | Cholangiocarcinoma |
| COAD | Colon adenocarcinoma |
| DLBC | Lymphoid Neoplasm Diffuse Large B-cell Lymphoma |
| ESCA | Esophageal carcinoma |
| GBM | Glioblastoma multiforme |
| HNSC | Head and Neck squamous cell carcinoma |
| KICH | Kidney Chromophobe |
| KIRC | Kidney renal clear cell carcinoma |
| KIRP | Kidney renal papillary cell carcinoma |
| LAML | Acute Myeloid Leukemia |
| LGG | Brain Lower Grade Glioma |
| LIHC | Liver hepatocellular carcinoma |
| LUAD | Lung adenocarcinoma |
| LUSC | Lung squamous cell carcinoma |
| MESO | Mesothelioma |
| OV | Ovarian serous cystadenocarcinoma |
| PAAD | Pancreatic adenocarcinoma |
| PCPG | Pheochromocytoma and Paraganglioma |
| PRAD | Prostate adenocarcinoma |
| READ | Rectum adenocarcinoma |
| SARC | Sarcoma |
| SKCM | Skin Cutaneous Melanoma |
| STAD | Stomach adenocarcinoma |
| THCA | Thyroid carcinoma |
| THYM | Thymoma |
| UCEC | Uterine Corpus Endometrial Carcinoma |
| UCS | Uterine Carcinosarcoma |
| UVM | Uveal Melanoma |

**Table 4: PCAWG Study Abbreviations.** `https://dcc.icgc.org/projects/details`. List of studies, which were ultimately used in association testing (after filtering steps).

| Study Abbreviation | Study Name |
|---|---|
| BOCA-UK | Bone Cancer - UK |
| BRCA-EU | Breast ER+ and HER2- Cancer - EU/UK |
| BRCA-UK | Breast Triple Negative/Lobular Cancer - UK |
| BTCA-SG | Biliary Tract Cancer - SG |
| CLLE-ES | Chronic Lymphocytic Leukemia - ES |
| CMDI-UK | Chronic Myeloid Disorders - UK |
| EOPC-DE | Early Onset Prostate Cancer - DE |
| ESAD-UK | Esophageal Adenocarcinoma - UK |
| LICA-FR | Liver Cancer - FR |
| MALY-DE | Malignant Lymphoma - DE |
| MELA-AU | Skin Cancer - AU |
| OV-AU | Ovarian Cancer - AU |
| PACA-AU | Pancreatic Cancer - AU |
| PACA-CA | Pancreatic Cancer - CA |
| PAEN-AU | Pancreatic Cancer Endocrine neoplasms - AU |
| PAEN-IT | Pancreatic Endocrine Neoplasms - IT |
| PBCA-DE | Pediatric Brain Cancer - DE |
| PRAD-CA | Prostate Adenocarcinoma - CA |
| PRAD-UK | Prostate Adenocarcinoma - UK |
| RECA-EU | Renal Cell Cancer - EU/FR |

**Table 5: Cancer Type Names.** Cancer type names used in this study and respective cancer types from TCGA, PCAWG, and Hartwig which were assigned to it. EAC: oesophageal adenocarcinoma.

| Cancer Type Name | Discovery | | Validation |
|---|---|---|---|
| | TCGA Cancer Type | PCAWG Project ID(s) | Hartwig Cancer Type(s) |
| Bladder | BLCA | - | Urinarytract |
| Brain_glioma_low | LGG | PBCA-DE | Nervoussystem_Gliomas or _NA |
| Brain_glioma_multi | GBM | PBCA-DE | Nervoussystem_Gliomas or _NA |
| Breast | BRCA | BRCA-EU, BRCA-UK | Breast |
| Colon_Rectum | COAD, READ | - | Colon_Rectum |
| Kidney | KIRC, KIRP | RECA-EU | Kidney |
| Lung_ad | LUAD | - | Lung |
| Lung_sq | LUSC | - | Lung |
| Ovary | OV | OV-AU | Ovary |
| Prostate | PRAD | PRAD-CA, PRAD-UK | Prostate |
| Skin | SKCM | MELA-AU_Cutaneous | Skin_Melanoma or _NA |
| Stomach_Eso | STAD, ESCA (EAC only[4]) | GACA-CN, ESAD-UK | Stomach, Esophagus |

**Table 6: Overview of sample sizes.** Corresponding cancer types for the cancer type names can be found in Table 5.

| Cancer Type Name | Discovery cohort sample size | Validation cohort sample size |
|---|---|---|
| Bladder | 323 | 87 |
| Brain_glioma_low | 405 | 283 |
| Brain_glioma_multi | 253 | 283 |
| Breast | 684 | 656 |
| Colon_Rectum | 410 | 417 |
| Kidney | 445 | 168 |
| Lung_ad | 434 | 299 |
| Lung_sq | 373 | 299 |
| Ovary | 199 | 180 |
| Prostate | 386 | 443 |
| Skin | 403 | 370 |
| Stomach_Eso | 363 | 431 |
| Pancan | 6,799 | 4,683 |
